# Carotenoid-based immune response in sea cucumbers relies on newly identified coelomocytes – the carotenocytes

**DOI:** 10.1101/2025.06.02.657395

**Authors:** Wambreuse Noé, Bossiroy Estelle, David Frank, Vanwinge Céline, Fievez Laurence, Bureau Fabrice, Gabriele Sylvain, Karasiewicz Tania, Mascolo Cyril, Wattiez Ruddy, Eeckhaut Igor, Caulier Guillaume, Delroisse Jérôme

**Affiliations:** Biology of Marine Organisms and Biomimetics Unit, Research Institute for Biosciences, University of Mons, 7000 Mons, Belgium; Belaza Marine Station (IH.SM-UMONS-ULIEGE), 601 Toliara, Madagascar; Direction Générale Déléguée à la Recherche, l’Expertise, la Valorisation et l’Enseignement (DGD REVE), Muséum National d’Histoire Naturelle (MNHN), Station Marine de Concarneau, 29900 Concarneau, France; Flow Cytometry Platform, GIGA Research Institute, University of Liège, 4000 Liège, Belgium; Laboratory of Cellular and Molecular Immunology, GIGA Research Institute, University of Liège, 4000 Liège, Belgium; Mechanobiology & Biomaterials Group, Research Institute for Biosciences, CIRMAP, University of Mons, 7000 Mons, Belgium; Proteomics and Microbiology Unit, Research Institute for Biosciences, University of Mons, 7000 Mons, Belgium

**Keywords:** Immune cell, Hemocyte, Gene expression, Hydrovascular system, Reactive Oxygen Species, Antioxidant, Echinodermata, Deuterostome

## Abstract

Sea cucumbers (Holothuroidea, Echinodermata) are marine deuterostomes possessing a complex innate immune system composed of a wide diversity of immune cells—coelomocytes, making them compelling models for exploring the evolution of immunity. This study investigates the functional specialisation of coelomocytes within the two main echinoderm body fluids, namely the perivisceral fluid (PF) from the perivisceral cavity and the hydrovascular fluid (HF) from the hydrovascular— ambulacral system. Given their specific distribution restricted to the HF, hemocyte-like cells (HELs) are particularly investigated. In echinoderms, hemocytes have been described as reddish cells containing haemoglobin and thus presenting a function in oxygen transport. Using an integrative approach combining cell morphological analyses, pigment profiling and multi-omics technologies, we demonstrate, in the sea cucumber *Holothuria forskali*, that HELs harbour exceptionally high concentrations of carotenoids, primarily canthaxanthin and astaxanthin—potent antioxidant molecules responsible for their pigmentation. Transcriptomics and proteomics analyses reveal that HELs express candidate genes involved in the carotenoid metabolism pathway as well as catalase, an antioxidant enzyme. Additionally, spectral flow cytometry assays reveal that HELs do not produce reactive oxygen species in contrast to most coelomocyte types, reinforcing the hypothesis of their antioxidant function. HELs also contribute to the formation of large red bodies (i.e., coelomocyte aggregates) and increase in concentration following lipopolysaccharide injections, indicating an active role in immune defence. Given these results, we hypothesise that these cells act after the culmination of the immune response, forming an antioxidant shell around the cellular aggregates to mitigate oxidative stress from reactive oxygen species (ROS) produced within the aggregate while encapsulating pathogens, thus protecting the host tissues. The discovery of carotenoid-carrying coelomocytes constitutes the first report of pigmented coelomocytes in sea cucumbers (except respiratory pigments), challenging the long-standing assumption that these cells contain haemoglobin. Therefore, we propose renaming hemocytes into carotenocytes, at least in this species. However, we think that this newly described coelomocyte type has been wrongly identified as haemoglobin-containing cells in many previous studies and could be present in many other holothuroid species. Our findings thus establish a new paradigm in the study of coelomocytes in echinoderms as well as the function of the hydrovascular system, unique to this phylum.

## Introduction

Echinoderms represent a phylum of marine deuterostomes that share many molecular features with chordates (Gross et al., 1999; Hibino et al., 2006; Perillo et al., 2024). These metazoans present a complex innate immune system that primarily relies on coelomocytes – specialised circulating immune cells suspended in the fluids of coelomic cavities (Boolootian and Giese, 1958; Hetzel, 1963; Kindred, 1924; Smith et al., 2018). These include the hydrovascular and perivisceral fluids, which could, roughly speaking, be considered functional analogues of blood (Kindred et al., 1924; Wahltinez et al., 2023). The hydrovascular fluid (HF) fulfils the hydrovascular system, a system unique to echinoderms, which performs a wide range of functions, including locomotion, transport of metabolites and support of body structure (Deridoux et al., 2025; Flammang et al., 1998; Potts, 2003; Strathmann, 1975). The perivisceral fluid (PF) is the fluid surrounding the organs in the general cavity and is known to play critical functions in the transport of metabolites and humoral factors and water balance regulation (La Paglia et al., 2025; Wahltinez et al., 2023). Over the past decades, coelomocytes have been shown to play a wide range of immune functions, including the recognition and elimination of foreign materials and pathogens, phagocytosis, aggregation, encapsulation and the production of a wide variety of humoral factors (Chiaramonte and Russo, 2015; Smith et al., 2018; Xing et al., 1998). In addition, the sequencing of the genome of the sea urchin *Strongylocentrotus purpuratus* has revealed a wide variety of genes encoding pathogen recognition receptors, making these organisms very interesting models for investigating the evolution of innate immunity (Hibino et al., 2006). In this context, numerous studies have examined the transcriptomics response of coelomocytes to various stressors (Dong et al., 2014; Nair et al., 2005; Wambreuse et al., 2025; Wu et al., 2020), highlighting, among other things, the expression of homologues of complement system components (Gross et al., 1999; Xiao et al., 2022), known to interact with the adaptive immune systems of vertebrates and contribute to opsonisation. This example demonstrates that the study of the immune system of echinoderms offers cutting-edge perspectives on the evolution of the immune system in deuterostomes. Nevertheless, while PF coelomocytes have been the subject of in-depth studies (e.g., Dong et al., 2014; Smith et al., 2019), HF coelomocytes have received comparatively little attention. This could be partly due to the technical difficulty of collecting sufficient HF, especially in sea urchins or brittle stars, in which the rigid endoskeleton limits access to the hydrovascular appendages containing a low volume of fluid. Sea cucumbers, on the other hand, with their prominent HF appendages, including the large Polian vesicle(s) (e.g., Caulier et al., 2024, 2020), and their soft bodies, represent an appropriate model for studying the hydrovascular system in general and, in particular, the function of hydrovascular cœlomocytes.

Sea cucumbers are also of considerable interest for a variety of reasons. First, many species play a pivotal role in marine ecosystems by representing important components of benthic macrofauna and by participating in sediment bioturbation (Clements et al., 2024; Purcell et al., 2016). Secondly, some species possess a high commercial value due to their exploitation in traditional Chinese pharmacopoeia and gastronomy (Purcell et al., 2023). To this purpose, some species are farmed under aquaculture conditions with complete life cycle control (Han et al., 2016; Hamel et al., 2022). The model species of this study, the European sea cucumber *Holothuria forskali*, has been considered in particular in the context of integrated multi-trophic aquaculture (IMTA) (David et al., 2024; MacDonald et al., 2013). Thirdly, sea cucumbers are a source of bioactive compounds extensively explored for the development of drugs with antibiotic and anticancer properties, among others (Janakiram et al., 2015; Pangestuti and Arifin, 2018; Santos et al., 2016; Schillaci et al., 2013). For instance, the farmed Chinese sea cucumber *Apostichopus japonicus* is rich in astaxanthin, a powerful antioxidant used in many pharmaceutical products (Hossain et al., 2022). These various characteristics highlight that sea cucumbers warrant research interest from both a fundamental and applied point of view.

Like vertebrates, sea cucumbers have different immune cell types and appear to have the highest diversity among the five echinoderm classes (Smith et al., 2018), with between six and nine types depending on the classification (e.g., six in Chia and Xing, 1996 and Smith et al., 2018; eight in Hetzel, 1963; and nine in Queiroz and Custódio, 2024). The most accepted cell types are phagocytes, spherule cells, progenitor cells, fusiform cells, crystal cells and hemocytes. Nevertheless, it should be noted that the presence of these cells is highly species-dependent and that many different names have also been used to refer to them in the literature (reviewed by Queiroz and Custódio, 2024), which has caused much confusion. Coelomocyte classifications are mainly based on morphological criteria, however, functional information on the different cell types is scarce. For example, only one study provides transcriptomics data on coelomocyte subsets in sea cucumbers (Yu et al., 2023), and these subsets themselves constitute a mixture of “spherical cells” and “lymphoid-like cells” (i.e. not directly related to any previous morphological classification). Among the coelomocyte types, the function and distribution of hemocytes, a type of coloured cell thought to contain haemoglobin (Kindred, 1924), remain particularly enigmatic. While previous reports claim that they have a function in oxygen transport and are limited to the holothuroid orders Molpadida and Dendrochirotida (Baker and Terwilliger, 1993; Fontaine and Hall, 1981; Hetzel, 1963; Kindred, 1924), recent research has shown that they have a wider distribution than previously described, also occurring in the order Holothuriida (formerly within a larger taxon, the Aspidochirotida) (Caulier et al., 2024). Furthermore, it has been shown that these cells can participate in the encapsulation process, and it is suggested that their haemoglobin would release reactive oxygen species (ROS) during the immune response (Caulier et al., 2020, 2024; Jobson et al., 2022). These studies have also shown that the hydrovascular fluid of *H. forskali* is particularly rich in this cell type (Caulier et al., 2024), thus offering a promising opportunity to deepen our knowledge about these pigmented coelomocytes.

The present study, therefore, seeks to investigate the differences in the immune response of circulating coelomocytes between the two main body fluids of sea cucumbers, namely the hydrovascular and the perivisceral fluids, using an integrative approach combining morphological analyses, pigment profiling and multi-omics tools. With their particular distribution, mainly localised in the HF, the function of cells similar to hemocytes (i.e., colourful reddish coelomocytes) is specifically studied. These cells will be referred to as hemocyte-like cells (HELs) in this study to avoid any confusion and prior functional conclusions related to the presence of haemoglobin. Overall, this study provides new insights into the immune response of sea cucumbers and establishes new paradigms on the function of HELs, which we describe here as a new functional coelomocyte type.

## Results

### 1. Perivisceral and hydrovascular fluids have distinctive coelomocyte populations

A total of 12 cellular elements could be distinguished in the HF and PF of *H. forskali* (**Fig. 1**), including three types of spherule cells (morula cells, large spherulocytes and small spherulocytes), two types of phagocytes (petaloid and filiform phagocytes), small round cells (SRCs), hemocyte-like cells (HELs), fusiform cells, crystal cells, giant cells, minute corpuscles and spermatozoa. The different types of spherule cells were divided according to their size and the size of their granules: Morula cells were the largest, measuring 14.6 ± 2.8 µm (mean ± SD) with granules larger than 3 µm in diameter (**Fig. 1B and R**); large spherulocytes measured 10.8 ± 1.9 µm (**Fig. 1C, N and O**) while small spherulocytes measured 6.5 ± 1.1 µm (**Fig. 1D and R**), both with granules between 1 and 2 µm in diameter. Petaloid and filiform phagocytes, measuring 23.1 ± 11.5 µm (including pseudopodia), could be identified based on the morphology of their pseudopodia, harbouring lamellipodia (**Fig. 1E and P**) and filopodia (**Fig. 1F and Q**), respectively. However, many phagocytes presented pseudopodia of various shapes and were difficult to assign to one or the other subtype. Therefore, all phagocytes were considered as one cell type in the cell count. The small round cells were smaller and had an undifferentiated appearance, measuring 5.3 ± 1.6 µm (**Fig. 1G and T**). Hemocyte-like cells (HELs) were also small cells with highly variable diameters ranging from 1.5 to 8 µm, with a mean size of 3.3 ± 1.5 µm (**Fig. 1H and U**). These were mainly recognised through their reddish colour (**Fig. 1H**), and their propensity to form uniform cell aggregates (**Fig. 1U**). Fusiform cells were distinguished based on their two opposite pseudopodia, which are sometimes branched at the end. These cells measured 27.3 ± 8 µm long (**Fig. 1I and S**). Crystal cells were recognised through their typical prismatic crystalline inclusion; it should be noted that the SEM image only represents a putative crystal cell, as the crystalline inclusion might also correspond to a methodological artefact, and only a few of them were observed (**Fig. 1J and V**). They measured 8.2 ± 1.6 µm. In addition to conventional coelomocyte types, two cell types were observed only a few times; they correspond to giant cells measuring 44.1 ± 5.4 µm with a light orange pigmentation (**Fig. 1K**) and minute corpuscles, which appeared empty with a very regular spherical shape, and a small size of 2.6 ± 0.4 µm (**Fig. 1L**). It is uncertain whether these cell types are coelomocyte types or results of tissue contamination, or cell debris release. Finally, spermatozoa could also be distinguished only in some male individuals and were sometimes very abundant in the two body fluids. They were identified mainly by their flagella (**Fig. 1M and W**) and their characteristic acrosome that was conspicuous in SEM preparations (**Fig. 1W**).

**Fig. 1.**
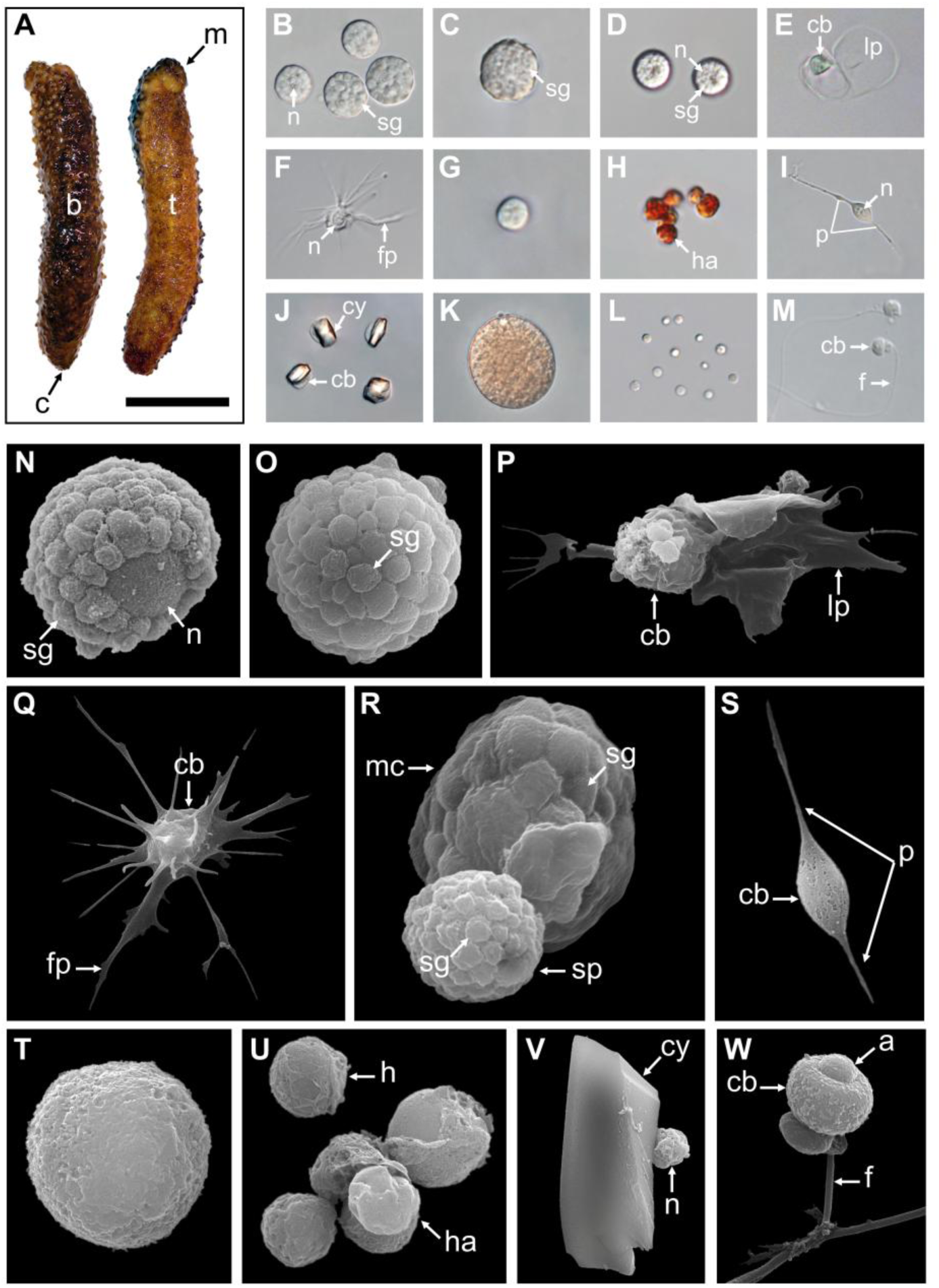
Cellular elements found in the perivisceral and hydrovascular fluids of *Holothuria forskali*. **A.** Aboral and oral views of a specimen of *H. forskali*. **B-M.** Light microscopy on coelomocytes: **B.** Morula cells. **C.** Large spherulocyte. **D.** Small spherulocytes. **E.** Petaloid phagocyte. **F.** Filiform phagocyte. **G.** Small round cell (SRC). **H.** Hemocyte-like cells (HELs). **I.** Fusiform cell. **J.** Crystal cell. **K.** Presumed giant cell. **L.** Presumed minute corpuscles. **M.** Spermatozoon. **N-V.** Scanning electron microscopy on coelomocytes: **N.** and **O.** Large spherulocytes. **P.** Intermediate phagocyte. **Q.** Filiform phagocyte. **R.** Side-by-side, a Morula cell and a spherulocyte. **S.** Fusiform cell. **T.** Small round cell (SRC). **U.** Hemocyte-like cells (HELs). **V.** Presumed crystal cell. **W.** Spermatozoon. Legend: a – acrosome; b – bivium; c – cloaca; cb – cellular body; cy – crystal; f – flagellum; fp – filipodia; h – hemocyte-like cell (HEL); ha – HEL aggregate; lp – lamellipodia; m – mouth; mc – Morula cell; n – nucleus; p – pseudopodia; sp – spherulocyte; sg – secretory granules; t - trivium. The scale bar (in A) represents 8.5 cm in A; 38 µm in B; 19 µm in C; 24 µm in D; 35 µm in E; 31 µm in F; 17 µm in G; 17 µm in H; 34 µm in I; 32 µm in J; 35 µm in K; 31 µm in L; 26 µm in M; 3 µm in N; 3.4 µm in O; 5 µm in P; 6.2 µm in Q; 3 µm in R; 8.6 µm in S; 2.1 µm in T; 4 µm in U; 4.4 µm in V; 2.5 µm in W.

Only phagocytes, spherulocytes (large and small merged), Morula cells, small round cells, HELs, fusiform cells and crystalline cells were considered in cell concentration and proportion assessment as they were almost always present and easily distinguishable under light microscopy. In total, HF contained 1.40 ± 1.0 × 10^7^ cells per ml, which was significantly higher than PF, containing 3.28 ± 0.8 × 10^6^ cells per ml (**Table 1**; Wilcoxon signed-rank test; p = 0.03; W = 31; **Table S1** shows the full statistical analysis results). This difference was mainly driven by the high concentration of HELs in HF, with an average concentration of 1.20 ± 1.0 × 10^7^ cells per ml and a proportion of 80.6 ± 5.1 %. Importantly, in PF, HELs were particularly poorly represented, with a proportion of 0.8% ± 0.7% and were only observed in four of the six individuals considered for this assessment. Although HELs were overall almost absent in the PF of *H. forskali*, they could reach 20% in some individuals examined. In the PF, spherulocytes were the dominant cell type with a concentration and proportion of 1.2 ± 1.0 × 10^6^ cells per ml and 36.0 ± 5.9 %, respectively. Concentrations and proportions of HELs and spherulocytes were significantly different between the two fluids (p = 0.031; W = 21; **Table 1**). Phagocytes were the second dominant cell type in the two body fluids, and their concentration was similar at around 10^6^ cells per ml, with a proportion of 10.5 ± 11.3% in the HF and 30.0 ± 10.1% in the PF. Next, SRCs and Morula cells had similar concentrations between the two fluids at around ∼1.2 × 10^6^ cells per ml and ∼6 × 10^6^ cells per ml (**Table 1**). Finally, fusiform and crystal cells were both poorly represented, with less than 10^5^ cells per ml. Interestingly, crystal cells were significantly more concentrated in the PF at 2 ± 0 × 10^4^ cells per ml against 3.3 ± 8.2 × 10^3^ cells per ml in the HF (p = 0.037; W = -15). Overall, proportions were significantly different between the two fluids in all the cell types (p < 0.05) except for fusiform cells. However, these different proportions can be attributed to the high concentration of HEL, which only appears in HF.

**Table 1.** Comparison of concentration and proportion for each cell population between the hydrovascular fluid (HF) and perivisceral fluid (PF), in specimens of normal condition (no injection). Results are formulated as mean ± SD (n = 6) and p-values show significant differences between the two fluids (Wilcoxon signed rank test; * p-value ≤ 0.05; n.s. means not significant; na means not applicable; see **Table S1** for full results).

| Cœlomocyte types | Concentration (cells ml<sup>-1</sup>) |  |  | Proportion (%) |  |  |
| --- | --- | --- | --- | --- | --- | --- |
|  | HF | PF | P-value | HF | PF | P-value |
| **Phagocyte** | 9.28 ± 6.18 10<sup>5</sup> | 1.04 ± 0.45 10<sup>6</sup> | n.s. | 10.52 ± 11.33 | 30.98 ± 10.07 | * |
| **Spherulocyte** | 3.27 ± 1.44 10<sup>5</sup> | 1.19 ± 0.41 10<sup>6</sup> | * | 2.91 ± 1.45 | 36.04 ± 5.90 | * |
| **Morula cell** | 1.05 ± 1.28 10<sup>5</sup> | 1.41 ± 1.16 10<sup>5</sup> | n.s. | 1.07 ± 1.12 | 5.05 ± 3.14 | * |
| **Small round cell** | 5.78 ± 4.16 10<sup>5</sup> | 7.37 ± 1.61 10<sup>5</sup> | n.s. | 4.49 ± 2.69 | 23.44 ± 7.36 | * |
| **Hemocyte-like cell** | 1.20 ± 0.96 10<sup>7</sup> | 2.33 ± 2.34 10<sup>4</sup> | * | 80.62 ± 5.14 | 0.75 ± 0.71 | * |
| **Fusiform cell** | 5.67 ± 5.99 10<sup>4</sup> | 1 ± 1.37 10<sup>5</sup> | n.s. | 0.38 ± 0.35 | 3.09 ± 3.75 | n.s. |
| **Crystal cell** | 3.33 ± 8.16 10<sup>3</sup> | 2 ± 0 10<sup>4</sup> | * | 0.01 ± 0.03 | 0.64 ± 0.15 | * |
| **Total (all types)** | 1.40 ± 0.99 10<sup>7</sup> | 3.28 ± 0.83 10<sup>6</sup> | * | n.a. | n.a. | n.a. |

### 2. Coelomocytes aggregate upon fluid collection

Coelomocytes appeared to display high mobility after fluid collection and gathered, leading to cell aggregation. In both body fluids, this process began with a phagocyte settling on the slide and extending its pseudopodia (**Fig. 2A and D**). Then, some previously immobile coelomocytes started to achieve amoeboid movements and join the settled phagocytes, especially spherule cells that could suddenly migrate toward the aggregate by forming characteristic small spherical protrusions followed by an undulation throughout the cell membrane, giving them a figure-eight appearance (**Fig. S1B**; **Video S1**). In some cases, when they encountered an aggregate, spherule cells were seen to perform a sudden cell lysis, releasing their cytosol content, including numerous secretion granules within the aggregate (**Video S1 arrow; Fig. S1A; Fig. 2E**). Other cells were passively captured in the newly formed aggregate when encountering it through the fluid flow. All this process results in the formation of early, disorganised and loose stage I aggregates (**Video S2; Fig. 2A and 2E**). These aggregates, by a retrograde movement of the phagocytic pseudopodia, while capturing additional cells, became more compact, and if some early aggregates were close together, they could merge by a reciprocal tractive force of the pseudopodia (**Fig. 2B and 2F**). This results in aggregates at stage II with many pseudopodia scattered around it (**Video S3; Fig. 2B**; **Fig. 2G**). As the aggregates grew, the pseudopodia became more stretched, and some were gradually retracted within the aggregates. Aggregates also acquired a spherical shape and slowly detached from the slide, resulting in stage III aggregates (**Video S4; Fig. 2C and 2H**). These aggregates were capable of autonomous lateral movement on the slide. While the aggregation process was the same in the two body fluids, many HELs were passively captured in early aggregates in the HF. Most HELs were isolated, but many formed small aggregates composed solely of HELs. **Figure 2I** shows a large HEL aggregate surrounding an early aggregate, itself fusing with another early aggregate, all capturing isolated HELs and other coelomocytes (**Video S5**). In the upper right of the video, a small HEL aggregate remains immobile without the action of a phagocyte or another early aggregate. To confirm that HELs participate in the aggregate, even passively, a quantitative particle analysis was performed by automatically counting cells whose surface area is between 4 and 50 µm^2^ (corresponding to a diameter between 2.2 and 8 µm), targeting isolated HELs. This analysis shows that the number of isolated HELs decreased significantly throughout the time-lapse, indicating that HELs join the aggregates over time (**Fig. 2J**; Mann-Kendall trend test: z = -7.9; n = 51; p = 2.8 × 10^-15^). However, this does not necessarily mean that this decrease in free HELs results from an active movement of these cells towards the early aggregates. Indeed, the time-lapse video suggests that it was rather the fluid flow that caused their movement, but that once they encountered an aggregate or another cell, they strongly adhered to it (**Video S5**). Similarly, the mean aggregate size was followed during time-lapse, and it appears that it increased significantly over time (**Fig. 2F**; z = 7.3; n = 51; p = 1.8 × 10^-13^). This can be explained by the fact that isolated cells join aggregates and small aggregates merge together, resulting in fewer but larger aggregates. The automated numbering of HELs and measurement of aggregate size can be found in **Fig. S2**.

**Fig. 2.**
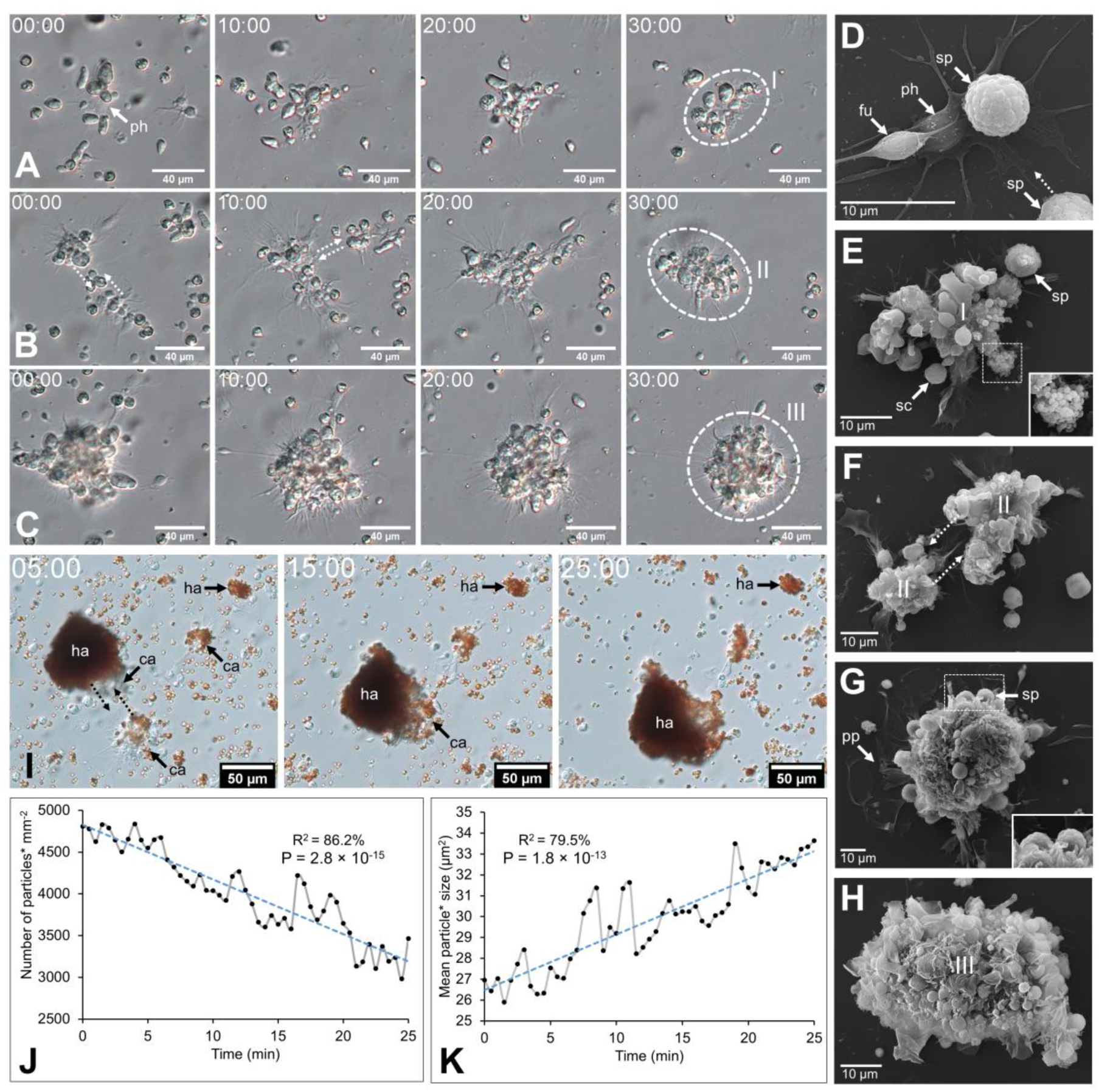
Coelomocyte aggregation just after fluid collection in *Holothuria forskali*. **A-C.** Time-lapse images of early aggregate formation in the perivisceral fluid. **A.** Formation of stage I aggregate is driven by a retrograde movement of cells encountering a phagocyte (ph) and their pseudopods (**Video S2**). **B.** Maturation of a stage I aggregate in stage II early aggregate (**Video S3**). **C.** Maturation of a stage II aggregate in a stage III aggregate (**Video S4**) **D-H.** SEM images of coelomocytes aggregating in the perivisceral fluid (PF), following what is observed by time-lapse imaging. **D.** Adherent phagocytes interact with other cells and lead to the formation of the initial aggregate (i.e., only a few cells). **E.** Other cell types are recruited, notably spherule cells, which release secretory granules through cell lysis (close-up view; **Video S1**). **F.** Fusion between two aggregates of stage II. **G.** Large compact aggregate of stage II, still showing numerous pseudopodia (close-up view of spherule cells taking part in the aggregate). **H.** Maturation of the aggregates, which become more compact and finally detach from the slide by retracting their pseudopod. **I.** Time-lapse images showing early aggregate formation in the hydrovascular fluid with HELs and HEL aggregates being passively captured in the early aggregates (**Video S5**). **J.** and **K.** Respectively, the number of small particles (i.e., the asterisk means cells with an area between 4 and 50 µm^2^; size targeting HELs; **Fig. S2B**) and the mean size of particles (i.e., the asterisk means cells and cell aggregates with an area superior to 4 µm^2^; **Fig. S2C**) per minute during 25 minutes (coefficient of determination – R^2^ and Mann-Kendall trend test p-value are included in the graphs). Legend: ca – coelomocyte aggregate; fu – fusiform cell; ha – HEL aggregate; ph – phagocyte; pp – pseudopodia; sc – small round cell; sp – spherule cell.

### 3. Only hemocyte-like cells increase one day after immunostimulation with lipopolysaccharides

Significant changes in cell populations following lipopolysaccharide injections were only found in the HF (**Fig. 3**). Firstly, in terms of concentration, the total number of coelomocytes and HELs differed significantly, with a clear increase in LPS-injected individuals (**Fig. 3A**; Mann-Whitney U test; p < 0.01). In PF, although an overall rise in the concentration of all coelomocyte types was observed in LPS-injected individuals, none were significant (**Fig. 3B**; p > 0.05). Regarding proportions in HF, most cell types differed significantly, except Morula cells and crystal cells (**Fig. 3C**; p < 0.05). Only HELs increased in proportion in LPS-injected individuals, with a proportion of 85.3 ± 6.6% in LPS-injected against 41 ± 25.1% in control individuals (p = 0.0047). As only HELs change in concentration in the HF, this increase explains why the proportion of other cell types is significantly lower. Finally, the proportions in the PF were stable between control and LPS-injected individuals, except for Morula cells and SRCs, which showed a slight increase and decrease in LPS-injected individuals, respectively, although these were not significant (**Fig. 3D**; p > 0.05). Proportion and concentration values, as well as the results of the statistical tests, can be consulted in **Table S2**.

**Fig. 3.**
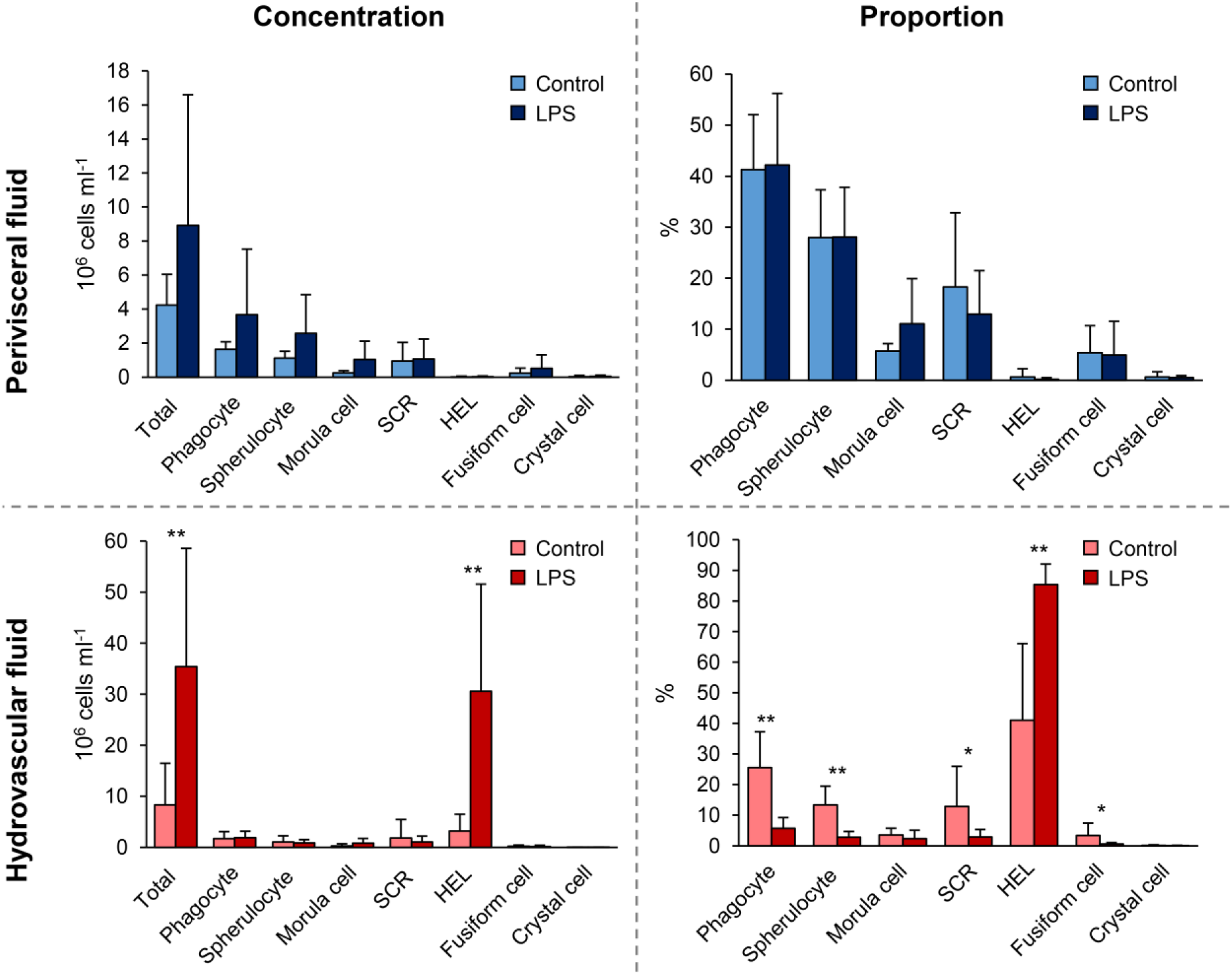
Comparison of cell population concentrations and proportions 24 hours after injections of lipopolysaccharides between control-injected individuals (artificial sterile seawater; control) and lipopolysaccharide-injected individuals (same solution containing 5 mg ml^-1^ of LPS). Scale bars represent the SD (n = 7 in the LPS condition and n = 6 in the control condition), and asterisks show significant differences (Mann-Whitney U test; * p ≤ 0.05; ** p ≤ 0.01; results of the statistical analysis are shown in **Table S2**).

### 4. Coelomocytes have a specific gene expression profile following a lipopolysaccharide challenge

#### 4.1. Transcriptome metrics and quality control

A total of 14 cDNA libraries were sequenced, including 6 HF cell samples, 6 PF cell samples, 1 stone canal sample and 1 podia sample sequenced for other projects (**Table S3**). They generated 3.33 billion reads with 93.0% clean reads and a Q20 of 99.0%. The *de novo* assembly yielded a total of 167,199 unigenes ranging from 30,861 to 73,673 unigenes for individual samples and with an average length, a N50 and a GC content that were 2,017, 3,807 and 38.9%, respectively.

The unigene length distribution showed that most unigenes were between 300 and 3,000 bp in length (76.2%; **Fig. S3A**). To obtain an initial indication of the function of the unigenes, the transcriptome was aligned with seven functional databases; 45.5% had at least one annotation, with Nr having the highest proportion of annotated unigenes (41%), followed by KEGG (31.7%) and InterPro (30.7%; **Fig. S3 B**). For the Nr annotation, the distribution of species shows a majority of echinoderm species (< 77.8%), with the most represented species being *Apostichopus japonicus* (60.4%; **Fig. S3C**). To assess the completeness of the transcriptome, the BUSCO statistic analyses were used; while individual transcriptomes showed a variable degree of completeness, the merged transcriptome showed a high percentage of complete orthologs (97.2%), most of which were duplicated (75.6%) (**Fig. S3D**).

#### 4.2. Differential gene expression

Three differential analyses were performed to compare expression between control and LPS-injected individuals; one considered only HF samples (HF analysis), one only PF samples (PF analysis) and one both (merged fluid analysis). The merged fluids analysis yielded a total of 17,646 DEGs of which 10,113 genes were up-regulated and 7,533 down-regulated; the PF analysis yielded 5,524 DEGs of which 2,702 genes were up-regulated and 2,822 down-regulated and the HF analysis yielded 6,420 DEGs of which 4,124 genes were up-regulated and 2,296 down-regulated (**Fig. 4A**). Comparing the three analyses in a Venn diagram shows that 2450 DEGs were common between PF and HF analysis with only 2 not found in the merged fluid analysis (**Fig. 4B**). A principal component analysis (PCA) was performed on the DEGs from the merged analysis (filtered by quantile-quantile to keep only the 25% most informative DEGs), showing a clear distinction between control and LPS-injected individuals with a higher variability within LPS-injected individuals (**Fig. 4C**). For each analysis, a heat map was produced based on the DEGs (**Fig. 4D-F**). While the merged and PF analyses show a good clustering between control and LPS-injected individuals, in the HF analysis, the 4-LPS-HF sample forms an individual branch, possibly due to a sex influence, as it was the only male in the LPS-injected condition (**Table S3; Fig. S4**). Furthermore, in the merged analysis group, clustering reveals a strong influence of individuality rather than sex or body fluids (i.e., the HF-PF distinction). To obtain a functional overview of the DEGs, the twenty most enriched KEGG pathways were selected for each analysis and a comparison was performed to reveal potential specific pathways in the immune response of each body fluid (**Fig. 4F**). Of the twenty most enriched pathways, 14 were common to all three analyses and 16 were common to the PF and HF analyses. A ranking was established based on the q-value (adjusted p-value), representing the significance of the enrichment for each pathway. All three analyses presented the same most enriched pathways, namely ”Nod-like receptor signalling pathway”, followed by “PI3K-Akt signalling pathway” in the merged and HF analysis and “Necroptosis” in the PF analysis. Surprisingly, the “PI3K-Akt signalling pathway” was not among the twenty most enriched pathways in the PF analysis. In addition, the “RIG-I-like receptor signalling pathway”, which was the third most enriched pathway in the PF analysis, was only the seventeenth most enriched pathway in both the merged and HF analyses. The most represented KEGG category in all three analyses was “Immune system”, with the following pathways: “NOD-like receptor signalling pathway”, “Toll and Imd signalling pathway”, “RIG-I-like receptor signalling pathway”, “IL-17 signalling pathway” and “Toll-like receptor signalling pathway” and “C-type lectin receptor signalling pathway”. The pathways with the highest rich factors (RF) were “Apoptosis - multiple species” in the merged and HF analyses and “RIG-I-like receptor signalling pathway” in the PF analysis. Note that the RF are generally higher in the merged analysis, which is logical since it includes a much larger number of DEGs (see **Fig. 4A**). Regarding the number of DEGs in the pathways, although the up and down-regulated ratio was predominantly between 0.8 and 1.2, it could vary significantly. The highest ratio (i.e., higher expression in the LPS condition) corresponds to “ECM-receptor interaction” and “Pathogenic *Escherichia coli* infection” (1) in the merged analysis; to “Hepatitis B” in the PF analysis (1.4) and to “Influenza A” in the HF analysis (2.2). Overall, these results show a clear response in gene expression to LPS injection, with a similar gene expression between the two fluids, which mainly corresponds to “immune system” pathways.

**Fig. 4.**
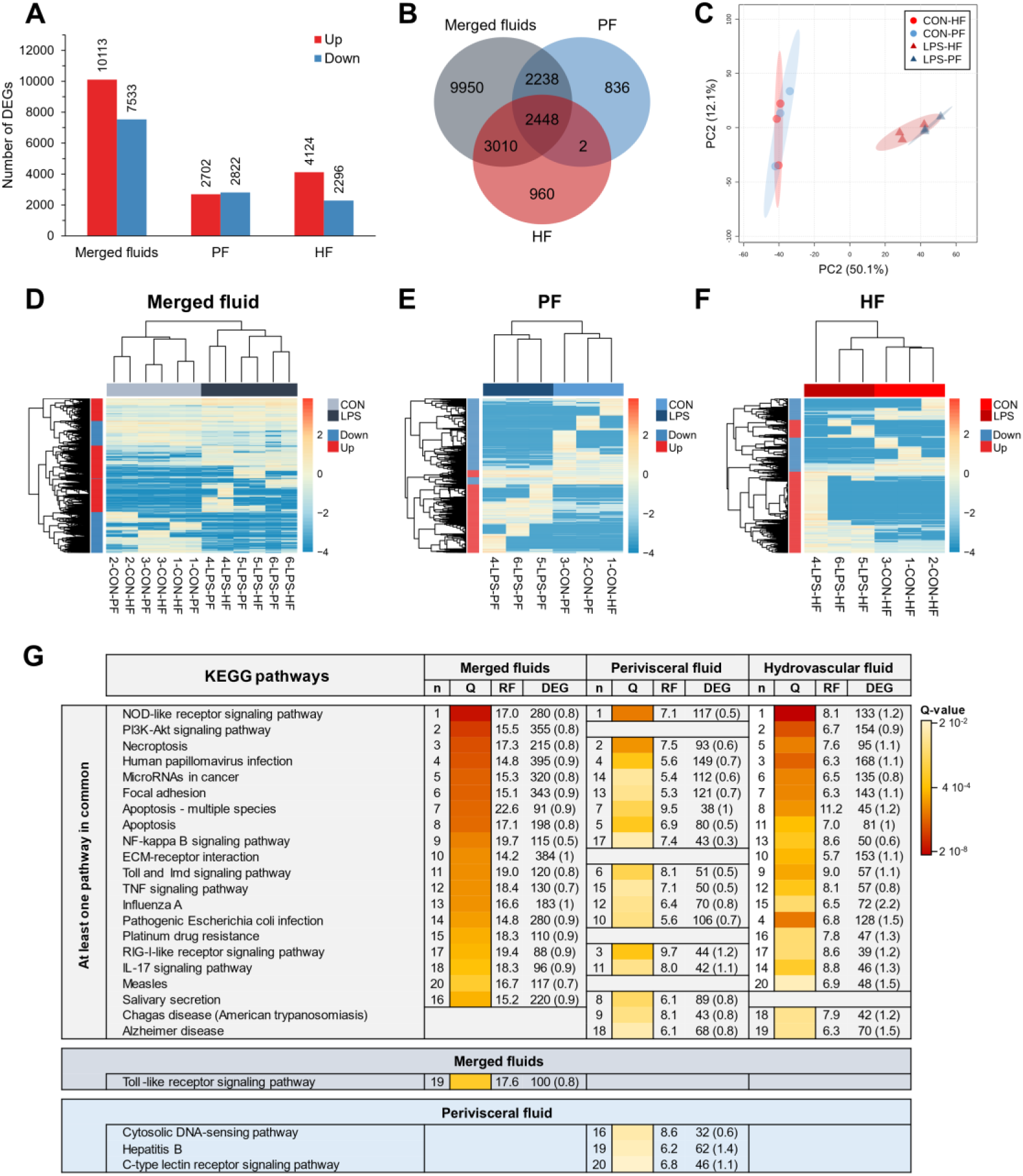
Differential expression analyses between control-injected (CON) and lipopolysaccharide-injected (LPS) individuals of *Holothuria forskali*. **A.** Number of up and down-regulated genes between CON and LPS individuals in the three analyses carried out: the first considering all body fluids; the second only within perivisceral fluid (PF) sample, and the third only within hydrovascular fluid (HF) samples (the exact number of differentially expressed genes (DEGs) for each analysis is included in the graph; the full list is shown in **Table S4**). **B.** Venn diagram of shared and unshared DEGs between the three analyses. **C.** Principal component analysis of DEG gene expression considering the analysis of CON vs LPS (i.e., considering all body fluids). **D-F.** Heatmaps based on the three differential expression analyses following the same order as in A. **G.** The twenty most enriched KEGG pathways in the three analyses; Pathways were ordered so that the degree of enrichment can be compared between the three analyses; n represents the ranking of the pathway; the q-value (Q) represents the significance of the enrichment according to a colour scale; the rich factor (RF) represents the percentage between the number of DEGs out of the number of unigenes in a given pathway; DEG represents the number of DEGs annotated in the pathways and, in brackets, is indicated the ratio between up and down-regulated within these DEGs. The full list of KEGG enrichment analyses between CON and LPS individuals can be found in **Table S5**.

#### 4.3. Sex-specific differential expression

Since a sex influence was suggested in the expression of coelomocytes, an additional expression analysis was performed to assess the influence of sex on gene expression (**Fig. S4**). A large number of differentially expressed genes (DEGs) were identified (**Fig. S4A**), with a clear distinction between male and female samples (**Fig. S4. B and C**). This could be partly explained by the presence of spermatozoa in the fluids of male individuals, probably resulting from contamination during the collection of coelomic fluids (**Fig. S4B**). Enrichment analyses revealed that many immune pathways were significantly differentially expressed between the two sexes (**Fig. S4D-F**). However, it cannot be excluded that this difference is due to the presence of spermatozoa in the male samples.

### 5. Coelomocytes exhibit distinct gene expression profiles in hydrovascular and perivisceral fluids

To compare expression between PF and HF samples, three differential analyses were performed; one considered only control samples (CON analysis), one only LPS-injected samples (LPS analysis) and one both (merged condition analysis). The merged analysis yielded a total of 2,853 DEGs, of which 2,070 genes were up-regulated and 783 down-regulated in the HF; the CON analysis yielded 179 DEGs, of which 141 genes were up-regulated and 38 down-regulated in the HF, and the LPS analysis yielded 2,773 DEGs, of which 2,124 genes were up-regulated and 649 down-regulated in the HF (**Fig. 5A**). The Venn diagram shows that only 43 DEGs were shared by all three analyses, which is mainly explained by the low number of DEGs in the CON analysis (**Fig. 5B**). Similarly, only 45 DEGs were shared between the CON and LPS analyses, but the LPS and merged analyses shared 1,552 DEGs. The PCA based on the merged analysis showed a clear distinction between the HF and PF samples, however, this separation was less pronounced under control conditions, which tended to cluster more closely along the PC1 (**Fig. 5C**). While the heatmaps of merged and LPS analyses show a relatively good distinction between PF and HF, clustering in the CON analysis shows an inaccurate separation (**Fig. 5D-F**). In the merged analysis, PF sample 3-CON-PF is in the HF cluster and sample 4-LPS-HF forms an isolated branch; in the CON analysis, several samples alternately form individual isolated branches, and only PF samples 3-CON-PF and 1-CON-PF fit the expected clustering correctly; and in the LPS analysis, sample 4-LPS-HF forms an isolated branch but PF samples form a good cluster. With regard to the comparison of the twenty most enriched pathways between the three analyses, only three pathways are shared. However, 12 pathways are shared between the merged analysis and the HF analysis, consistent with the higher number of DEGs that are shared between these two analyses. The three most enriched pathways were “Proteoglycans in cancer”, “Hypertrophic cardiomyopathy” and “Dilated cardiomyopathy” in the merged analysis; “NF-kappa B signalling pathway”, “HIF-1 signalling pathway” and “Estrogen signalling pathway” in the CON analysis; and “Adherens junction”, “Maturity onset diabetes of the young” and “Human papillomavirus infection” in the LPS analysis. In terms of KEGG categories, the most represented system is the “Endocrine system” (7 terms), followed by the “Digestive system” (4 terms) and the “Immune system” (3 terms). The pathways with the highest richness factors (RF) were “Maturity onset diabetes of the young” in the merged analysis (1.5%) and LPS analysis (21.2%), and “Antigen processing and presentation” (1.4%) in the CON analysis. In terms of the ratio of up-regulated to down-regulated genes, many pathways had only up-regulated genes (represented by “up” in **Fig. 5G**). In the merged analysis, only one pathway had only up-regulated genes, it was “Maturity onset diabetes of the young” and the minimal ratio was “ECM-receptor Interaction” and “MicroRNA in Cancer” (both at 1.1). In the CON analysis, 19 out of 20 pathways contained only up-regulated genes, the last being “Cell adhesion molecules” with a ratio of 4. In the LPS analysis, six pathways contained only up-regulated genes, the lowest ratio being “PI3K-Akt signalling pathway” (0.9%).

**Fig. 5.**
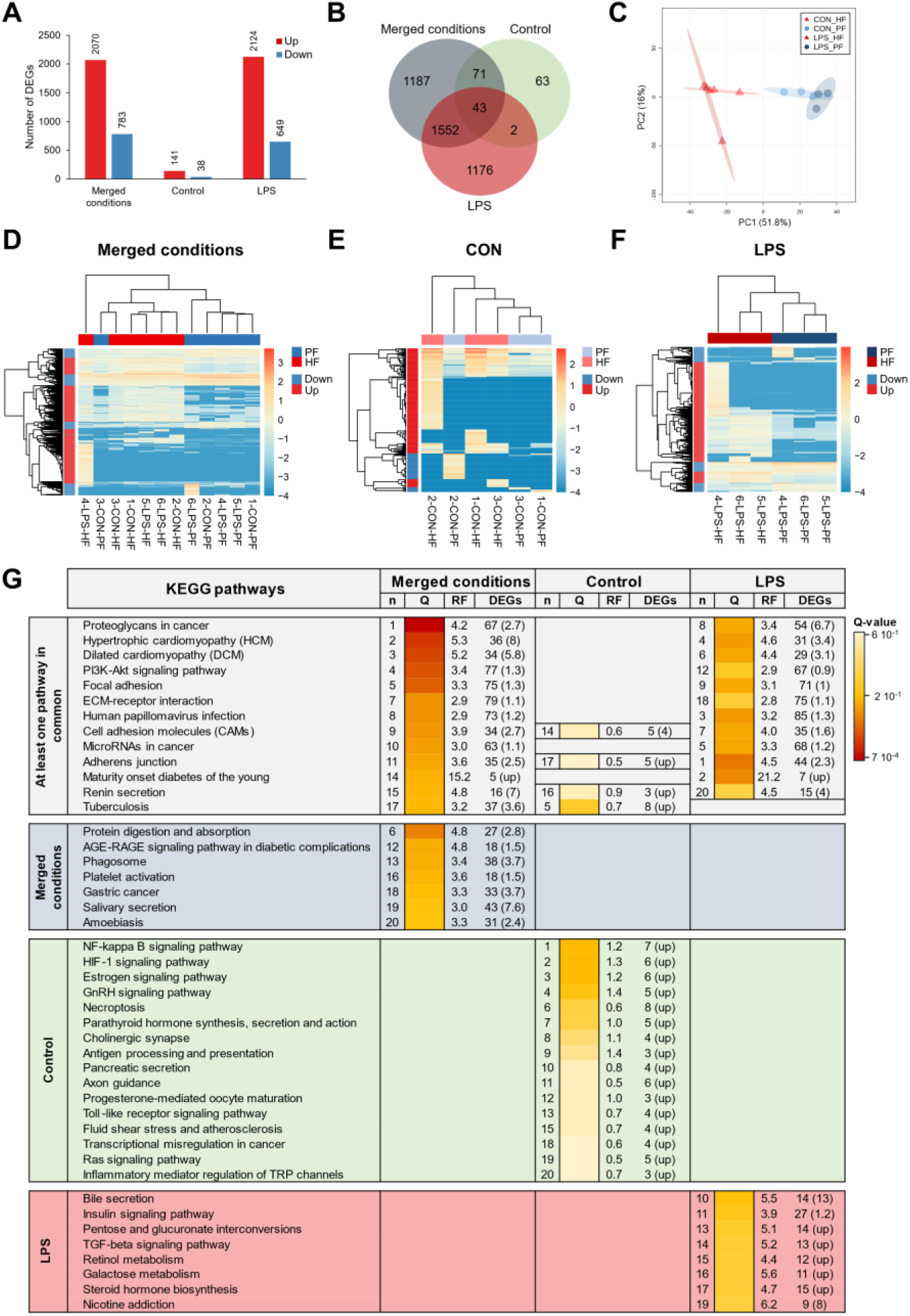
Differential expression analyses between the perivisceral fluid (PF) and the hydrovascular fluid (HF) in *Holothuria forskali*. **A.** Number of up and down-regulated genes between the PF and the HF in the three analyses carried out: the first considering all conditions; the second only within control individuals (CON) and the third only within LPS individuals (the exact numbers of differentially expressed genes (DEGs) are included in the graph; the fill list is shown in **Table S6**). **B.** Venn diagram of shared and unshared DEGs between the three analyses. **C.** Principal component analysis of DEG gene expression considering the analysis between PF vs HF (i.e., considering the two conditions). **D-F.** Heatmaps based on DEGs of the three differential expression analyses following the same order as in A. **G.** Twenty most enriched KEGG pathways in the three analyses; pathways were ordered so that the degree of enrichment can be compared between the three analyses; n represents the ranking of the path; the q-value (Q) represents the significance of the enrichment according to a colour scale; the rich factor (RF) represents the percentage between the number of DEGs out of the number of unigenes for a given pathway; DEG represents the number of DEGs annotated in the path and, in brackets, the ratio between up and down-regulated within these DEGs (“up” means that all genes in this pathways were up-regulated). The full list of KEGG enrichment analyses between the PF and HF can be found in **Table S7**.

### 6. Coelomocytes contain carotenoid pigments, which are more abundant in the hydrovascular fluid

A comparative pigment analysis was carried out between the two fluids to characterise the nature of the red pigmentation of the HELs. First, a liquid-liquid extraction was carried out to verify the pigment preference for polar (water-methanol) or apolar (chloroform) solvents. In addition to the two body fluids (i.e., HF and PF), the analysis includes a mammal blood sample (*Bos taurus*) as a positive control, given the hypothesis that hemocyte pigmentation is due to haemoglobin. The results show that reddish-orange pigmentation is mainly observed in the non-polar phase of the HF samples, with some PF samples showing a slight pink colouration in their non-polar phase (**Fig. 6A**). The blood sample, in contrast, shows no colouration in the two phases, with most of the colour visible in the interphase, corresponding to pelleted cellular debris in the polar phase.

**Fig. 6.**
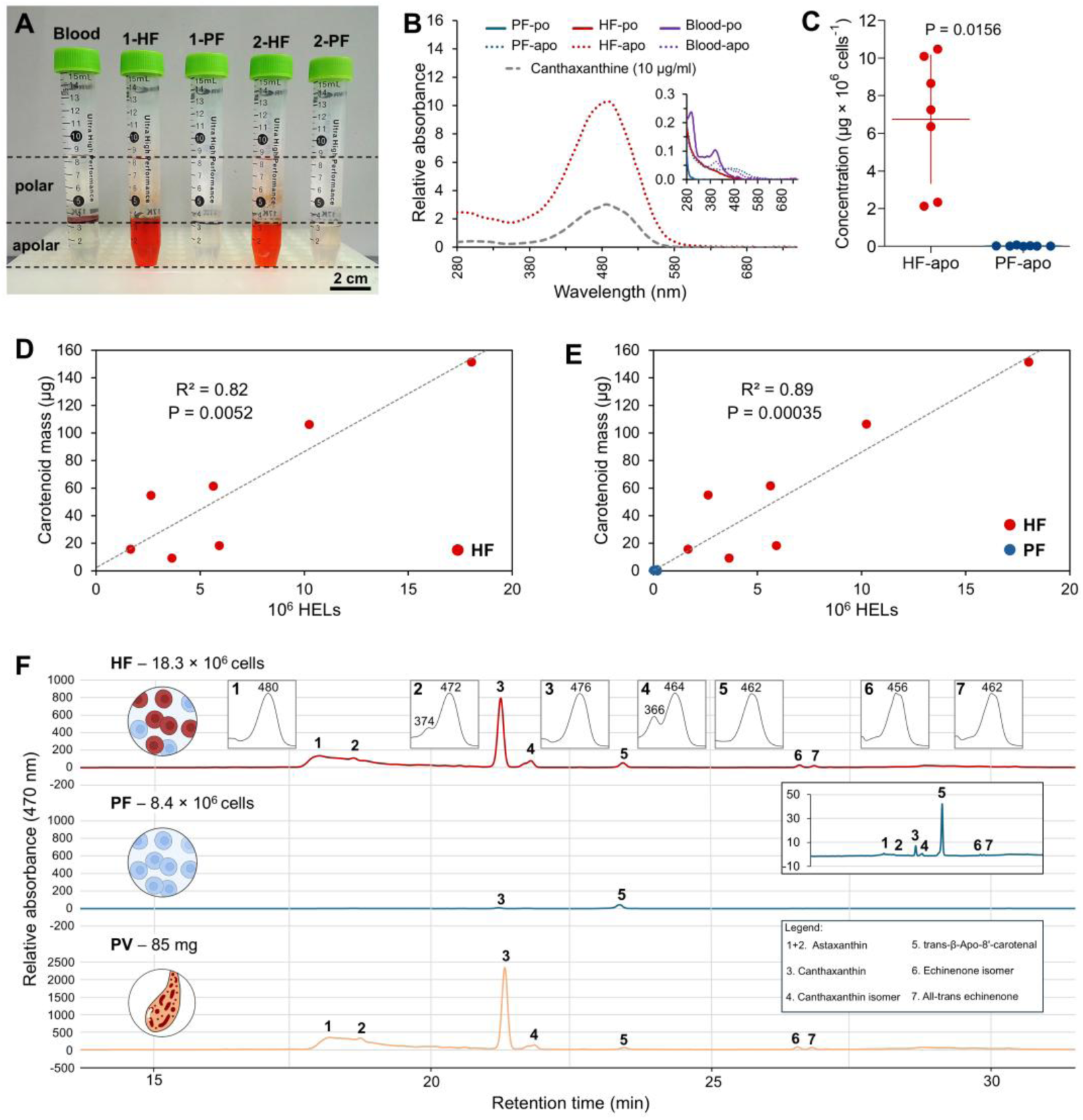
Pigment profiling of coelomocytes from the hydrovascular fluid (HF) and the perivisceral fluid (PF) of *Holothuria forskali*. **A.** Liquid-liquid extraction of pigment from HF and PF coelomocytes, as well as a bovine blood sample for haemoglobin positive control. The polar phase (methanol + water) surrounds the apolar phase (chloroform). Only the apolar phases of the HF samples show red pigmentation. **B.** Absorbance spectra of pigments extracted from the polar and apolar phases of HF and PF samples and from bovine blood (the apolar phase of the PF sample, polar phases and blood samples are shown in the zoomed graph, as their absorbance was too low to be distinguished in comparison with the spectrum of the apolar phase of HF). The spectra were compared to that of a canthaxanthin standard, which peaks at the same wavelength as the apolar phase extract of HF. **C.** Comparison of the pigment concentration between HF and PF samples calculated based on a calibration curve acquired based on a canthaxanthin commercial standard (λ = 490 nm; **Fig. S5A**). The masses of pigment are normalised by a million cells (all coelomocyte types included). Scale bars represent the SD, and the p-value is included on the graph (n = 7; Wilcoxon signed-rank test). **D.** Relation between carotenoid mass and the number of HELs in the HF samples. **E.** Relation between carotenoid mass and the number of HELs in the PF and HF samples. **F.** High-performance liquid chromatography (HPLC) carotenoid profile at 470 nm of HF, PF, and Polan vesicle samples in *H. forskali* (smaller graph shows absorbance spectra corresponding to each peak). Peaks were identified with commercial standards (**Fig. S5B**). Peaks 1+2 correspond to astaxanthin, peak 3 to canthaxanthin, peak 4 to a canthaxanthin isomer, peak 5 to the injected standard used for quantification (trans-β-Apo-8’-carotenal), peak 6 to an echinenone isomer and peak 7 to all-trans echinenone (Legend: apo – apolar phase; Rel. Abs – relative absorbance; HF – hydrovascular fluid; iso – isomer; PF – perivisceral fluid; po – polar phase; PV – Polian vesicle).

To determine the nature of the pigments in more detail, the polar and non-polar phases of each sample were analysed by spectrophotometry. The spectra corroborate the results obtained visually with a much higher absorbance in the apolar phase of the HF samples, showing a large relative absorbance peak at 490 nm corresponding to a red resulting colour (**Fig. 6B**). With regard the property of pigment (apolar and orange red colour), those measures were associated to the detection and identification of carotenoid pigments in High-Performance Liquid Chromatography (HPLC; see below). Canthaxanthin being the most abundant carotenoids, was used as a standard in spectrophotometry and shows a very similar spectrum to the HF apolar samples (**Fig. 6B**). Using a calibration curve (**Fig. S5A**), it was possible to calculate the pigment mass per sample and to standardise it by the number of cells in each pellet. This revealed a significant difference between the two fluids with 6.8 µg ± 3.4 µg per 10^6^ cells in HF compared to 2.6 ± 2.3 × 10^-2^ µg per 10^6^ cells in PF (Wilcoxon signed-rank test: p = 0.0156; W = -28; **Fig. 6C**). These concentrations appear consistent with the proportion of HELs in the two fluids, which was 77.3 ± 14.2% in HF and 0.8 ± 0.7% in PF. In this way, the carotenoid mass was correlated with the number of HELs only in the HF samples (**Fig. 6D**), and in the PF and HF samples (**Fig. 6E**), which revealed strong relationships with high coefficients of determination (R^2^) of 89.3% and 81.6%, respectively (Spearman’s test: S = 82.6; p = 3.5 × 10^-4^ in D; Pearson’s test: df = 5; t = 4.7; p = 5.2 × 10^-3^ in E).

In parallel with the quantification of carotenoids, HPLC analysis was used to identify the types of pigments present in the two different body fluids as well as in the Polian vesicle (PV) samples (rich in large red bodies; see section 7). It reveals the presence of different types of carotenoids, including, in decreasing order of abundance, canthaxanthin and its isomer, astaxanthin, and echinenone and its isomer (**Fig. 6F**). These could be identified thanks to injected standards (**Fig. S5B**) and quantified with trans-β-Apo-8’-carotenal (peak 5; **Fig. 6F**). **Table 2** shows the concentration and proportion of each type of pigment. The most abundant ones, namely canthaxanthin and astaxanthin, accounted for approximately 55% and 45%, respectively, in the various samples (including the isomer). Furthermore, the concentration of astaxanthin and canthaxanthin appears to be highly correlated in both fluids, as evidenced by a linear relationship with a high coefficient of determination (R^2^ = 99.0%: **Fig. S6**). The proportion of echinenone was less than 5% in the HF and Polian vesicle samples, with a slightly higher proportion in the PF at 7.76 ± 4.8% (including the isomer). Importantly, while the proportion of pigment was relatively constant between samples, their concentration was much higher in HF than in PF, corroborating the results obtained by spectrophotometry. However, it should be noted that the total carotenoid content calculated by HPLC is much higher than the one obtained by spectrophotometry (e.g., for HF samples: 28.7 ± 9.6 µg per 10^6^ cells by HPLC compared to 6.8 µg ± 3.4 µg per 10^6^ cells by spectrophotometry) which could be explained by different sensitivity of the two techniques. The Polian vesicle samples are also very rich in carotenoids (i.e., the carotenoid profile and the relative absorbance was similar to that of the HF samples; **Fig. 6F**). Their total carotenoid content was 0.3 ± 0.1 µg per µg of dry mass, thus representing approximately 30% of the total dry mass of the samples.

**Table 2.** Concentrations and proportions of carotenoids measured by high-performance liquid chromatography (HPLC) in the hydrovascular fluid (HF), the perivisceral fluid (PF) and the Polian vesicle (Pv) of *Holothuria forskali*. Peaks* correspond to the spectra shown in **Fig. 6F**. The concentration for body fluid samples (HF and PF) is calculated per million cells, including all coelomocyte types. The concentration for the Polian vesicle samples (^$^) is calculated based on the dry weight (“na” means not applicable). Results are presented as mean ± SD (n = 4).

| Peak* | Pigment | Concentration<br>($\mu\text{g} \times 10^6 \text{ cell}^{-1}$) | | Concentration<sup>§</sup><br>($\mu\text{g} \times \mu\text{g}^{-1}$) | Proportion (%) | | |
| --- | --- | --- | --- | --- | --- | --- | --- |
|  |  | HF | PF | PV | HF | PF | PV |
| 1 + 2 | Astaxanthin | 13.1 ± 8.4 | 0.6 ± 0.8 | 1.5 ± 1.1 $\times 10^{-2}$ | 42.1 ± 8.1 | 30.4 ± 9.8 | 43.4 ± 6.8 |
| 3 | Canthaxanthin | 13.2 ± 7.1 | 0.7 ± 0.7 | 1.6 ± 0.8 $\times 10^{-1}$ | 49.3 ± 8.3 | 48.8 ± 8.8 | 48.5 ± 6.1 |
| 4 | Canthaxanthin isomer | 1.5 ± 0.9 | 0.2 ± 0.2 | 1.8 ± 1.1 $\times 10^{-2}$ | 5.6 ± 0.9 | 13.1 ± 2.2 | 5.6 ± 1.3 |
| 6 | Echinenone isomer | 0.3 ± 0.2 | 0.04 ± 0.03 | 3.5 ± 2.7 $\times 10^{-3}$ | 1.1 ± 0.6 | 3.5 ± 2.8 | 1.1 ± 0.6 |
| 7 | All-trans-echinenone | 0.6 ± 0.4 | 0.06 ± 0.05 | 5.3 ± 4.7 $\times 10^{-3}$ | 1.9 ± 0.6 | 4.2 ± 2.0 | 1.4 ± 0.4 |
| All | Total carotenoid | 28.7 ± 9.6 | 1.5 ± 1.0 | 0.3 ± 0.1 | na | na | na |

### 7. Hemocyte-like cells are distributed across the different tissues and can form red bodies

As previously demonstrated, HELs are mainly observed in the hydrovascular system. However, within this system, HELs can take different forms and localise to different compartments. Moreover, HELs could be observed in other tissues under certain conditions, such as in dehiscent gonadal tubules (personal observation). This section aims to further describe HELs and their pathways throughout the body of *H. forskali* (**Fig. 7**). From a macroscopic to a microscopic point of view, the first evidence of the presence of HELs is large red bodies measuring from 200 to 2000 µm (**Fig. 7A-D**). These bodies may be attached to the inner wall of the Polian vesicle and buccal tentacle ampullae (**Fig. 7D**) or free in the HF (**Fig. 7C**). Round red aggregate appears to detach easily when the Polian vesicle is shaken or poured (**Fig. 7B**). Most of the red bodies were round, but some were irregular (**Fig. 7D**). Flattening them under a microscope slice revealed some intact HELs but the red pigmentation appears to be distributed throughout the aggregate matrix, with some even more pigmented areas forming reddish patches. In addition to the HELs, other cells are visible but have an empty appearance and do not correspond to the types observed in the fluid cell suspension. Moving on to a more microscopic view, the inner membrane of the Polian vesicle shows a large number of cells which appear to be marginated, *i.e.* attached to the inner membrane. Among these cells, many spherule cells can be recognised as well as HELs, which are present in two forms: some are isolated, and others form small aggregates composed solely of HELs. These small HEL aggregates were generally composed of 2 to 50 cells and generally did not exceed 100 µm in diameter. Zooming in further on an individual with a more reddish Polian vesicle reveals a higher density of HELs forming a “cell bed” on the inner membrane of the Polian vesicle, although a few spherule cells are still visible. In addition to the red bodies and the HEL aggregate, early aggregates were observed in the HF, as described in Section 2. Finally, the examination of the cells in the HF enabled us to distinguish different sizes of HELs and HEL aggregates (**Fig. 7J and K**). In SEM, HELs were observed forming highly compact HEL aggregates of different sizes (ranging from single cells – stage I to clusters of several dozens of cells – stage IV), with isolated cells being the most abundant (**Fig. 7J**), suggesting strong intercellular interactions. In light microscopy, different size groups could be distinguished: size I measuring 1.7 µm, size II measuring 2.8 µm, size III measuring 5.4 µm, size IV measuring 7 µm, size V measuring 10 µm, and size VI measuring 15 µm (**Fig. 7K**). The most abundant sizes were sizes I and II, with size III also frequent; the three sizes accounted for more than 90% of the free HELs. Sizes IV and V showed a granular appearance and were larger. Size VI could be attributed to a HEL aggregate as in Fig. 7J and was also more common than sizes IV and V. However, the fact that it is very similar to size V (**Fig. 7K**) suggests that this cellular element might be the result of large HEL fission, releasing size I and size II HELs.

**Fig. 7.**
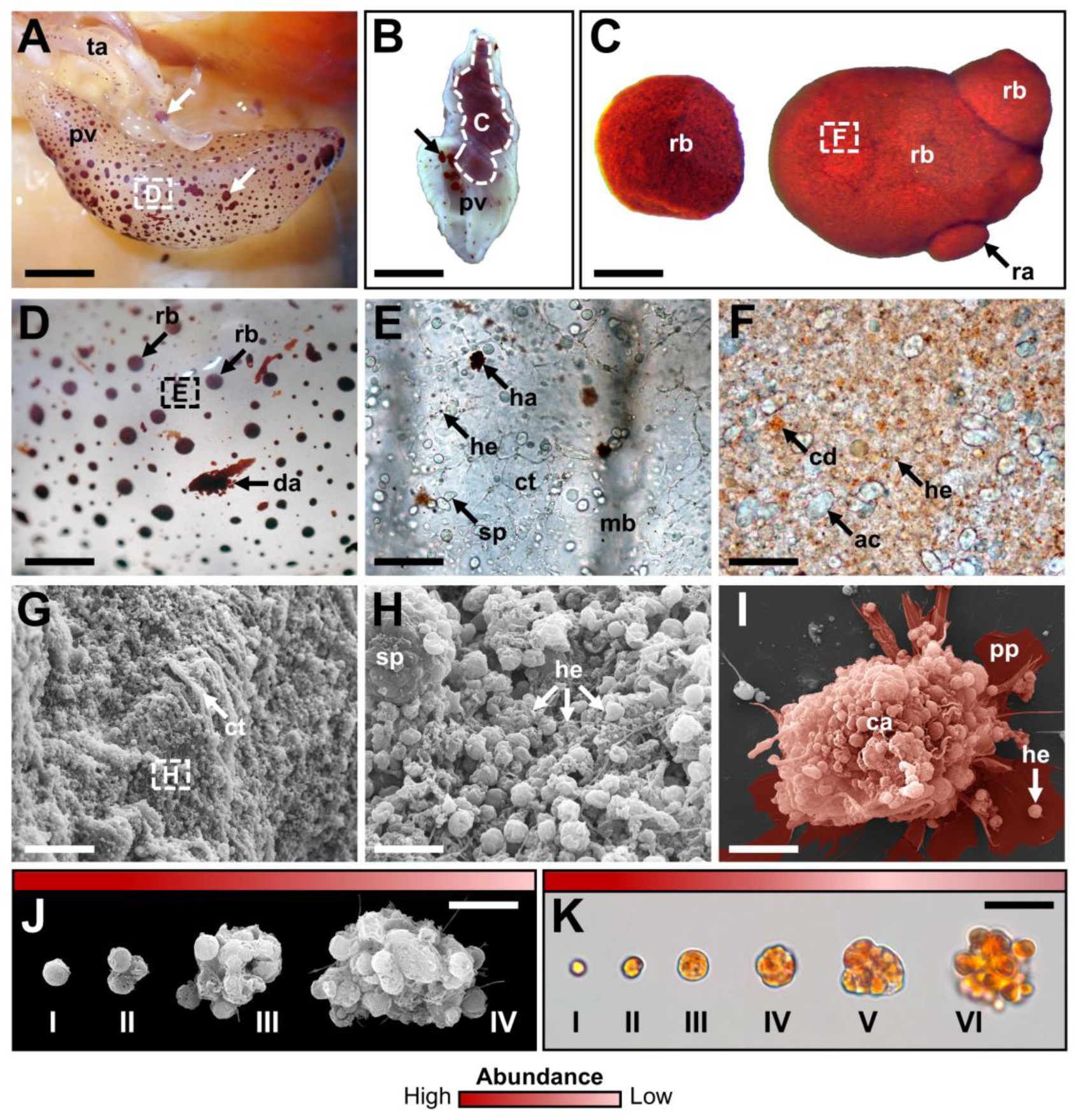
Distribution and morphological diversity of HELs. **A.** Polian vesicle and ampullae of the buccal tentacles showing marginated aggregates (indicated by white arrows). B. Poured Polian vesicle showing free red aggregates (indicated by black arrows). **C.** Red aggregates of different shapes are found in the hydrovascular fluid (HF) of the Polian vesicle. **D.** High magnification of marginated aggregates on the inner surface of the Polian vesicle; some disorganised outer aggregates are also visible. **E.** Inner surface of the Polian vesicle showing marginated HEL aggregates, HELs and spherulocytes. **F.** Cells constituting the red aggregates, including putative apoptotic coelomocytes and HELs; a few conspicuous coloured spots are visible, possibly corresponding to carotenoid deposition. **G.** SEM view of the inner surface of the Polian vesicle showing a large number of cells. **H.** Close-up (SEM view) on the inner surface of the Polian vesicle showing numerous marginated HELs and a large spherulocyte. **I.** Early aggregate stage II observed after incubation of the HF cells for 30 minutes on a slide; the colour indicates the cell aggregate and its pseudopodia, many HELs are conspicuous within the aggregate. **J.-K.** Different sizes of HELs and/or HEL aggregates, respectively, in SEM and light microscopy, illustrating the high adhesive properties of these cells as well as a potential fragmentation process of large HELs (their relative abundance is shown by a red scale; darker red means more abundant). Legend: ac – putative apoptotic cell; ca – coelomocyte aggregate; cd – putative carotenoid deposition; ct – connective tissue; da – disorganised aggregate; ha – HEL aggregate; he – HEL; pp – pseudopodia; mb – muscular band; pv – Polian vesicle; rb – red body; sp – spherulocyte; ta – tentacle ampulla; dotted contours refer to the subsequent images. Scale bars represent 950 mm in A; 110 mm in B; 340 µm in C; 45 mm in D; 75 µm in E; 100 µm in F; 25 µm in G; 5 µm in H; 11 µm in I; 6 µm in J; and 14 µm in K.3.7.

### 8. The autofluorescence signature of hemocyte-like cells allows their isolation by fluorescent-activated cell sorting (FACS)

When observing coelomocytes under a fluorescent microscope, it is possible to note that certain cell types present different autofluorescence levels at different excitation wavelengths. HELs, in particular, showed high autofluorescence at all the wavelengths tested, including dark violet 350 nm, blue 485 nm and green 589 nm, while at equivalent exposures, the other coelomocyte types were not visible (**Fig. 8A**). In order to obtain a more quantitative comparison of autofluorescence between the different populations, coelomocytes from both fluids were studied using spectral flow cytometry. With classical size (FSC) and granularity channels (SSC) (gating strategy in **Fig. S7**), 5 different populations were defined, including populations A, B and C in the HF and populations E and D in the PF (**Fig. 8B**; **Fig. S8** for full individual spectra). HELs were easily identified as population A by comparing the cell profile between fluids (as they are only present in the HF). In flow cytometry, this population corresponds to small granular cells. The comparison of autofluorescence spectra between the five populations (**Fig. 8C and D**) confirms that population A (corresponding to HELs) was the most autofluorescent population in all the channels tested and that this autofluorescence is higher than that of the other cell types. Furthermore, two other populations, namely population B of the HF and population D of the PF, showed slight autofluorescence in the dark violet (355nm) and violet (405nm) channels (**Fig. 8D**). They also have a higher granularity which, with their proportion, suggests that these cells might be spherule cells (*i.e.*, spherulocytes and Morula cells). The other two populations, population C of the HF and population E of the PF, were intermediate in size with small granularity. They were associated with cell types that do not have granules, notably phagocytes and SRCs.

**Fig. 8.**
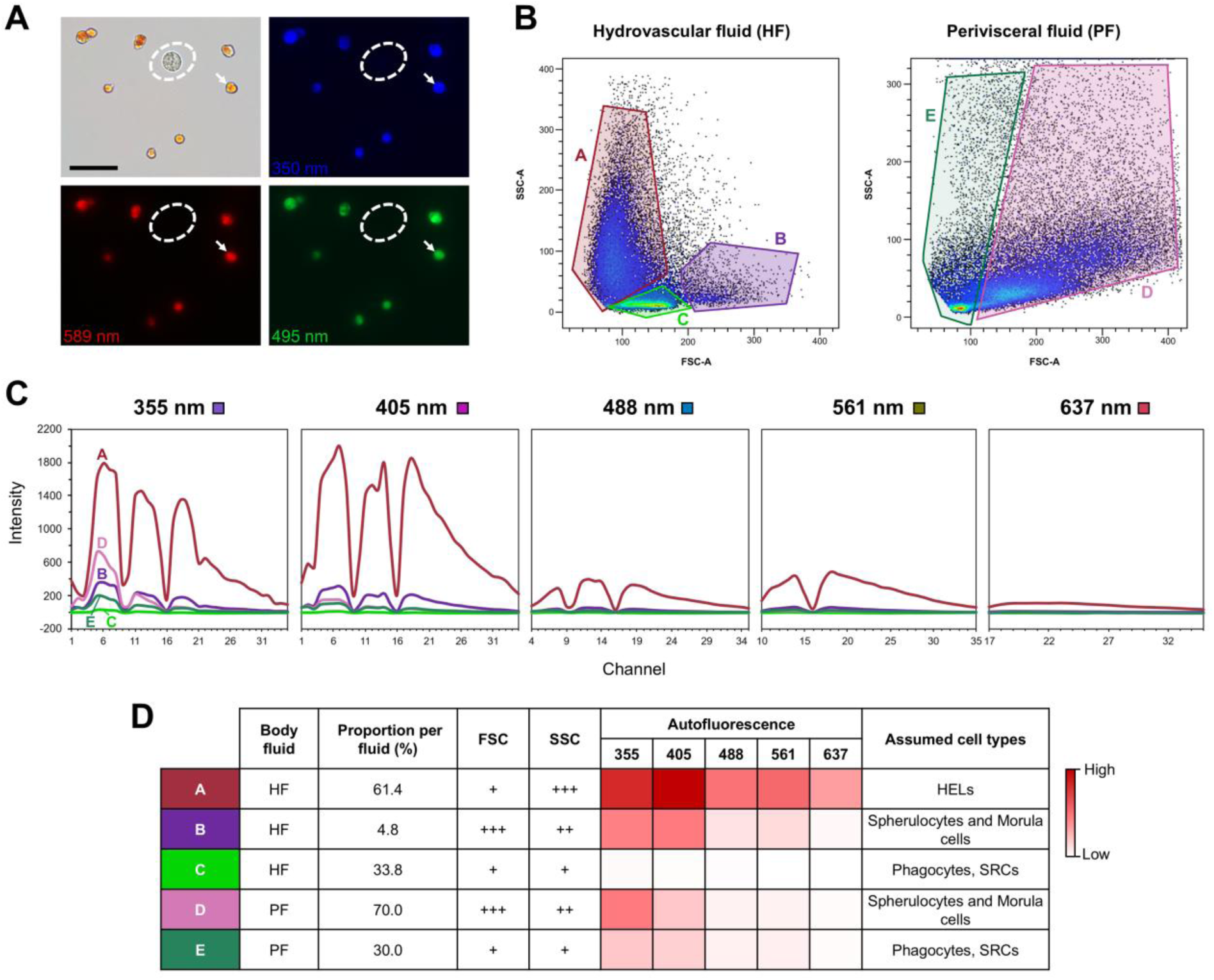
Autofluorescence of coelomocytes from perivisceral fluid (PF) and hydrovascular fluid (HF) revealed by spectral flow cytometry in *Holothuria forskali*. **A.** Coelomocytes from the HF observed by fluorescent microscopy at different excitation wavelengths (350, 495 and 589 nm; the dotted ellipse surrounds a spherulocyte, and the white arrow shows a hemocyte-like cell - HELs). **B.** Flow cytometry profile of HF and PF coelomocytes according to the cell size (forward scatter channel - FSC) and cell granularity (side scatter channel – SSC); the gating strategy is shown in **Fig. S7**. **C.** Autofluorescence spectra overlay for each population obtained by spectral flow cytometry; the graph shows the autofluorescence intensity for the different spectra, ranging from dark violet (355 nm) to red (637 nm). Individual autofluorescence spectra as well as resulting HF and PF total spectra are found in **Fig. S8**. **D.** Characteristics of each population: the corresponding body fluid; the proportion of the population (i.e., the number of events in the population out of the total number of events in the population of the fluid); FSC – size; SSC – granularity (+ means low; ++ means intermediate and +++ means high); the autofluorescence intensity in the different channels considered, represented by a colour scale (a darker colour means higher autofluorescence); cell type(s) are assigned to each population according to the different characteristics.

To confirm that HELs are carotenoid-carrying cells, we took advantage of the autofluorescence criteria established using spectral flow cytometry to develop a method for isolating HELs by FACS (**Fig. 9A**). Using the two fluorescent channels APC and Amcyan, it was possible to segregate the HEL population from the non- or less-autofluorescent populations (**Fig. 9A**). The fact that this population corresponds to HELs is confirmed when looking at a PF sample with the same channels, which does not show any fluorescent population. HELs were sorted from the HF sample with a HELs proportion of 85.1%. After the sorting, the purity of HELs was assessed by analysing the sorted sample with the same gating strategy, which resulted in a purity of 97.0%. In addition to this, the HEL purity was assessed by visual check under a microscope, which confirmed that no other cell types were present in the sample and that purified HELs by FACS still showed the same autofluorescent characteristics as freshly collected ones (**Fig. 9C**). A total of 2.1 × 10^6^ HELs were sorted to perform a targeted pigment analysis on this sample. Pigment analysis confirms that this sample has a very similar spectrum to canthaxanthin with a peak at 490 nm (**Fig. 9D**). By comparing the absorbance obtained with a canthaxanthin standard curve, it was possible to determine that the mass of carotenoid was 2.9 µg for this sample, i.e., 1.4 µg of carotenoid per 10^6^ HELs.

**Fig. 9.**
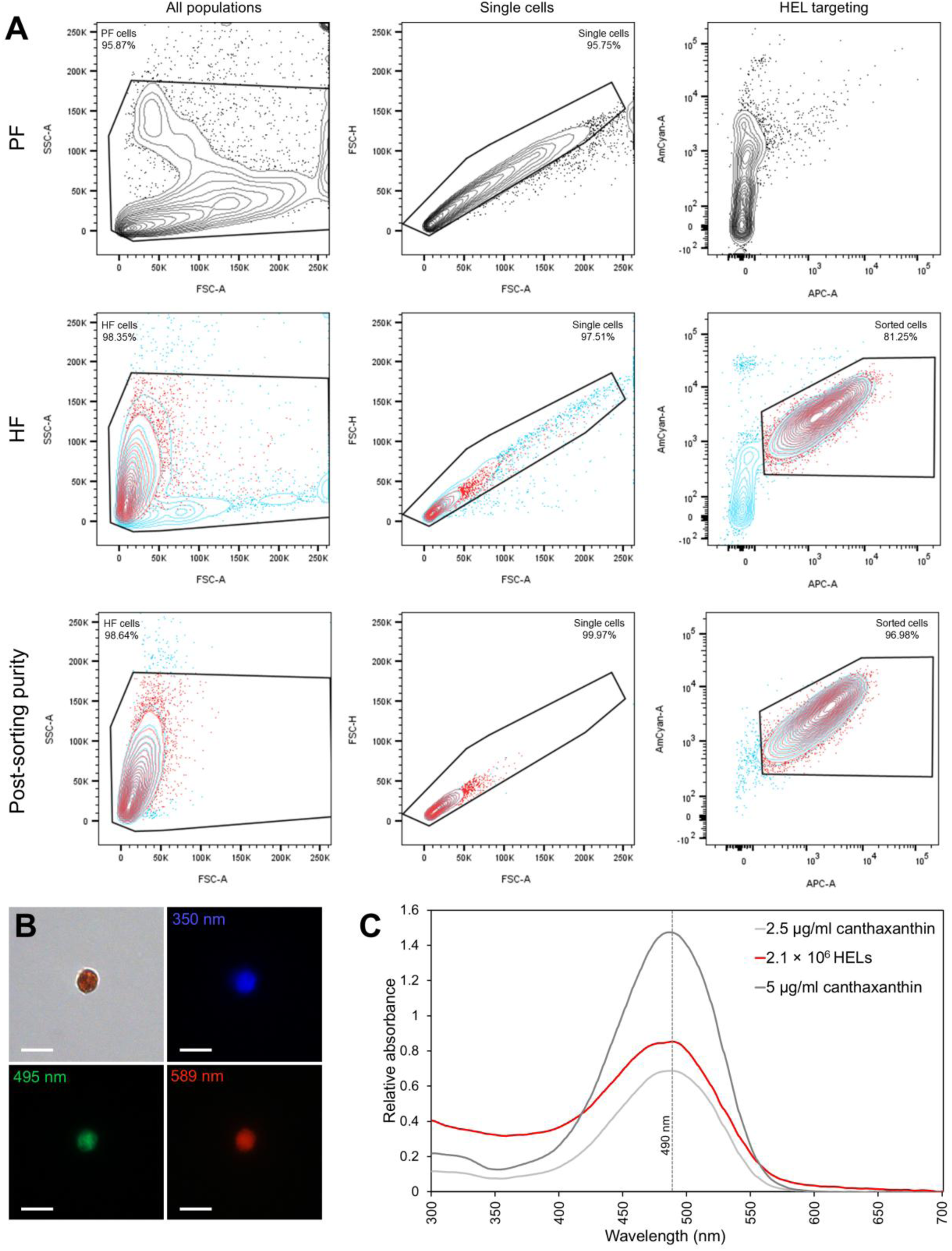
HELs isolated from the hydrovascular fluid of *Holothuria forskali* by Fluorescent-activated cell sorting (FACS). **A.** Sorting strategy based on autofluorescence of HELs. The HEL population is identifiable by comparing the HF and the PF samples using fluorescent channels (AmCyan and APC). The post-purity test confirms the sorting of a highly enriched HEL population. **B.** Autofluorescence of isolated HELs observed by fluorescent microscopy at different excitation wavelengths (λ = 350, 495 and 589 nm). The scale bar represents 10 µm. **C.** Comparison of the absorbance spectra between extracted pigment from the purified sample of HELs and the different dilutions of a canthaxanthin standard with known concentrations. The three spectra show a very similar pattern with a peak at 490 nm.

### 9. Hemocyte-like cells as a driver of differential expression between hydrovascular and perivisceral fluids

By hypothesising that each cell type has its own gene expression, the presence of HELs in HF and not in PF (or very few) should influence the differential expression between the two fluids. The first evidence of this can be observed in the differential expression analysis between the two fluids; while results of Section 3 demonstrate that HEL proportion increased in LPS-injected individuals, the differential expression was more marked between the two fluids when considering LPS individuals, suggesting a link between HEL proportion in the HF and differential gene expression. To further investigate this effect, the cell proportion and concentration were calculated for each sequenced sample (**Fig. 10A** and **10B**) and a differential expression analysis between the two fluids for each individual was carried out (based on fold change and Poisson distribution; **Fig. 10C**). Note some contaminating spermatozoa could also be find in coelomic fluids of male individuals. While they were not considered for the coelomocyte count in Section 3, for this analysis, they must have been taken into account as they also contain RNA that can dilute coelomocyte RNA. As found previously, while the concentration was highly variable between samples, LPS-injected individuals had a higher proportion of HELs in the HF at 82.3 ± 15.9% against 47 ± 30.8% in control individuals. Regarding differential expression analysis, the number of DEGs was variable between samples and seemed to correlate with the proportion of HELs in the HF (**Fig. 10C**). For example, individual 3 had the lowest number of DEGs between its two fluids at 3.7% (3,727 genes up-regulated in the HF and 696 genes up-regulated in the PF out of 118,025 genes) and also had the lowest proportion of HELs in its HF, namely 20.8%. The same analysis was done for genes that were expressed in only one fluid, namely the fluid-specific expressed genes (FSEGs), and it followed the same trend (**Fig. 10D**). Based on these observations, a significant relation was obtained between the proportion of HELs in the HF and the proportion of the DEG between PF and HF (Pearson’s correlation test: R^2^ = 86%; t = 5.0; df = 4; p = 0.0075; **Fig. 10E**), indicating that 86% of the variation in the number of DEGs between the two body fluids is explained by the proportion of HELs in the HF. Regarding the FSEGs, although the linear model had a lower coefficient of determination (R^2^ = 74%), the relation was also significant (Pearson’s correlation test: t = 3.6; df = 4; p = 0.029; **Fig. 10F**). Overall, these relations tend to demonstrate that most of the genes overexpressed in HF or only expressed in HF have a high probability of being overexpressed or only expressed in HELs.

**Fig. 10.**
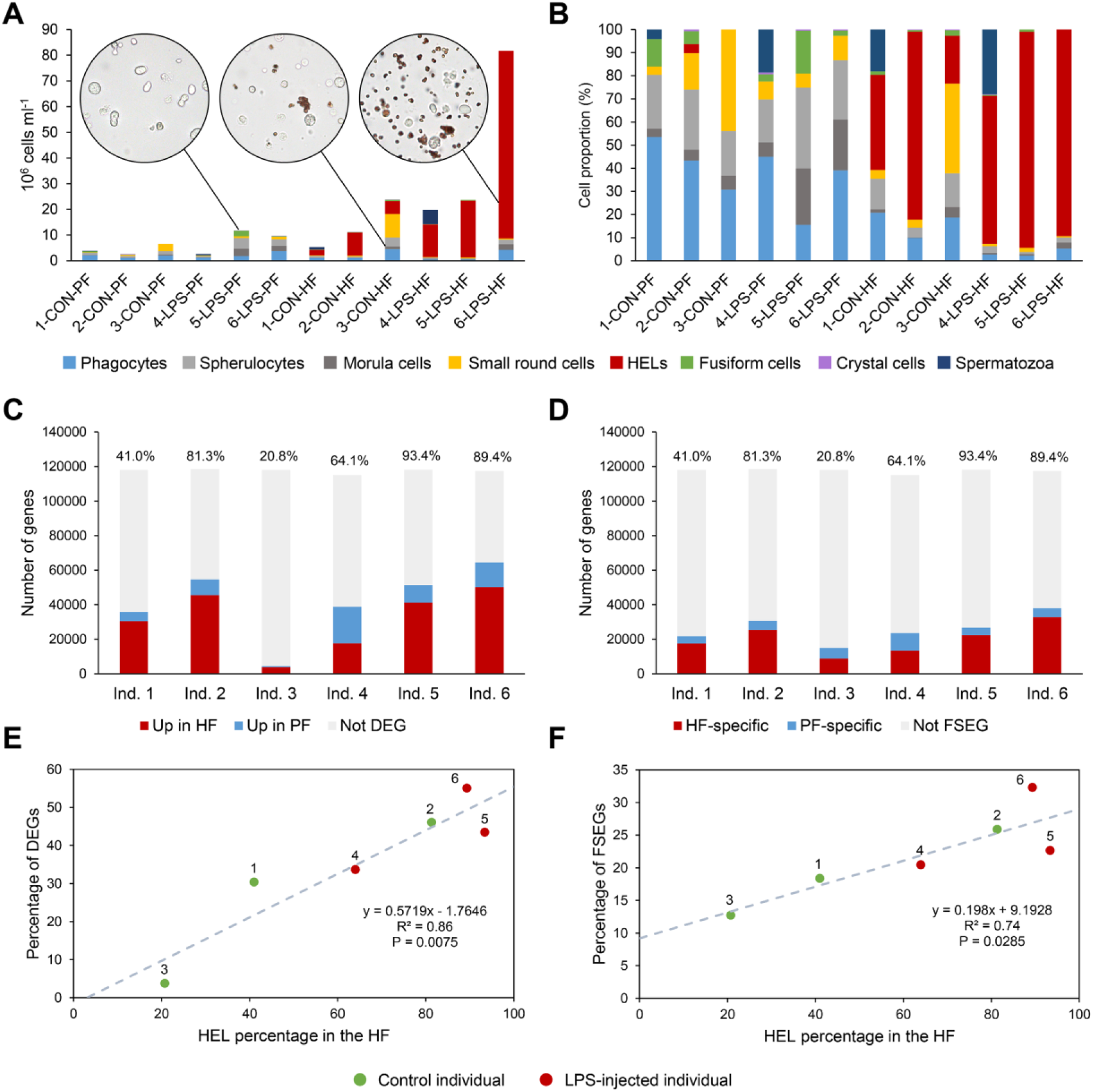
Relation between cell populations and differential gene expression between perivisceral fluid (PF) and hydrovascular fluid (HF) in *Holothuria forskali*. **A.** Cell population concentration in the different sequenced samples. Pictures of representative unprocessed fluids are included. **B.** Cell population proportion in the different sequenced samples. **C.** Number of differentially expressed genes (DEGs) in the comparison between HF and PF for each individual (based on the fold change and Poisson distribution). **D.** Number of fluid-specific expressed genes (FSEGs; i.e., only expressed in one of the two fluids). The percentage of HELs is indicated above the histograms in C and D, suggesting a correlation with the number of DEGs and FSEGs, respectively. **E.** and **F.** Relation between HELs proportion in the HF and proportion of DEGs in E or FSEGs in D (equation of the linear regression; coefficient of determination and p-value of the Pearson correlation test).

### 10. Proteomics validation of differential transcriptomics and search for candidate marker proteins of Hemocytes-like cells

To confirm the results obtained by differential transcriptomics and identify HEL-specific proteins, a proteomics comparison between the HF and the PF coelomocytes from the same individual was carried out by mass spectrometry (MS). The HF of this individual was highly enriched in HELs, accounting for 97% of the cells in this sample, constituting an almost pure population of HELs (**Fig. 11A**). The reason for such a high proportion of HELs in HF is unclear, but one possible explanation is that the individual used for this analysis was eviscerated, causing stress that may have triggered a specific response of HELs. Coelomocyte proportions in the PF fluid were similar to those found previously, with 49% of phagocytes followed by 31.5% of spherule cells (i.e., Morula cells and spherulocytes), 16.5% of SRC, and only 2.5% of HELs. A total of 1,073 proteins could be identified in both fluids, of which 361 were filtered out due to their peptide count being inferior to 2. Among the 712 remaining proteins, 618 were shared between the two body fluids, 20 were only expressed in the PF, and 74 were only expressed in the HF (**Fig. 11B**; the list of proteins is shown in **Table S8**). These proteins were annotated with the *de novo* transcriptome obtained from HF and PF coelomocytes (see section 4). While no protein had the same annotation (i.e., proteins annotated with the same unigene), one protein could be annotated by several genes, complicating the functional annotation of these proteins. To confirm the differential expression between the two body fluids, the fold change values obtained by mass-spectrometry were compared to the ones obtained by RNA-seq. For this analysis, only proteins that had only one matching gene in the list of DEGs between the two fluids were selected, resulting in ten proteins (see **Table S8** for the list of proteins and their annotation). A significant relation was found between the two techniques, validating, therefore, the RNA-seq differential expression analysis (**Fig. 11C**; R^2^ = 69.7%; Pearson correlation test: t = 4.3; df = 8; p = 2.6 × 10^-3^). To further investigate the function of HELs, proteins up-regulated in HF (i.e., enriched in HELs) were extracted from MS and RNA-seq analyses (i.e., translated from genes) to build functional protein networks against the proteome of *A. japonicus*. For the proteomics analysis, only proteins having a fold change > 5 were selected, giving a total of 133 proteins which, after extracting coding sequences from their annotation and aligning them against the *A. japonicus*’s proteome, resulted in 118 proteins (**Fig. 11D**). For the RNA-seq analysis, unigene with a fold change > 2 were selected. The same steps were followed based on translated peptide sequences, which resulted in 613 proteins in the final network (**Fig. 11E**; the result of the STRING mapping is shown in **Table S9**). For the two analyses, the network had significant protein interactions (string PPI p-value < 10^-16^). The same analysis was done for up-regulated proteins in the PF by RNA-seq and MS and the resulting networks had a lower number of proteins (108 and 52, respectively) with poorly significant PPI p-values (**Fig. S9**). Functional analyses were carried out on these networks using the “gene ontology” database to highlight common two biological processes that were conserved between MS and RNA-seq analyses, namely the “cell adhesion” process (FDR = 0.049 and 0.001, respectively) and the “Lipid metabolomic process” (FDR = 9.5 × 10^-8^ and 0.03, respectively). In addition, RNA-seq analysis revealed a local network cluster of 17 proteins corresponding to “Cytochrome P450, and steroid biosynthetic process” that was significantly enriched (FDR = 0.002; **Fig. 11E**). The ten most enriched biological processes for MS and RNA-seq are also shown in **Figure 11F** and **G**, respectively, to further characterise the global function of the protein networks. While MS analysis shows mainly lipid-related biological processes (e.g., “lipid catabolic process”, “cellular lipid catabolic process”, etc), RNA-seq analysis reveals mainly cytoskeleton-related processes (“cell adhesion”, “actin-filament based process”, “actin cytoskeleton organisation”, etc). Moreover, some processes are not the same between the two analyses, however, they can be seen as a part of the same biological functions, such as the “Steroid metabolic process” of the RNA-seq analysis, being considered as an overall lipid-related process. Furthermore, to highlight the most specific proteins in the HF sample, the fifteen proteins most differentially expressed in MS analysis were listed (**Fig. 11H**). The three most differentially expressed proteins were “putative development-specific protein LVN1.2 isoform X2” (log_2_FC = 13.5), “putative retinol dehydrogenase 12” (log_2_FC = 11.6) and “putative solute carrier family 15 member 4-like” (log_2_FC = 11.1). Regarding their peptide counts, they vary between 2 and 11, with “putative retinol dehydrogenase 12” having the highest, reflecting a more robust annotation for this protein (**Fig. 11H**). Finally, the annotation and expression of proteins from the local network cluster “Cytochrome P450 and steroid biosynthetic process” were analysed using a heat map visualisation for their interest in the carotenoid metabolism (**Fig. 11I**; Two proteins were filtered out by a t-test filter due to excessive variance). As expected, it shows a clear distinction between the two body fluids, with a higher expression in the HF. Moreover, proteins are divided into two groups, the highly expressed ones and the less expressed ones (**Fig. 11I**). Regarding the annotation, six proteins are notably annotated as “cytochrome P450”. Moreover, one protein is annotated as “putative retinol dehydrogenase 12” **Fig. 11H**), highlighting similar results between MS and RNA-seq analyses.

**Fig. 11.**
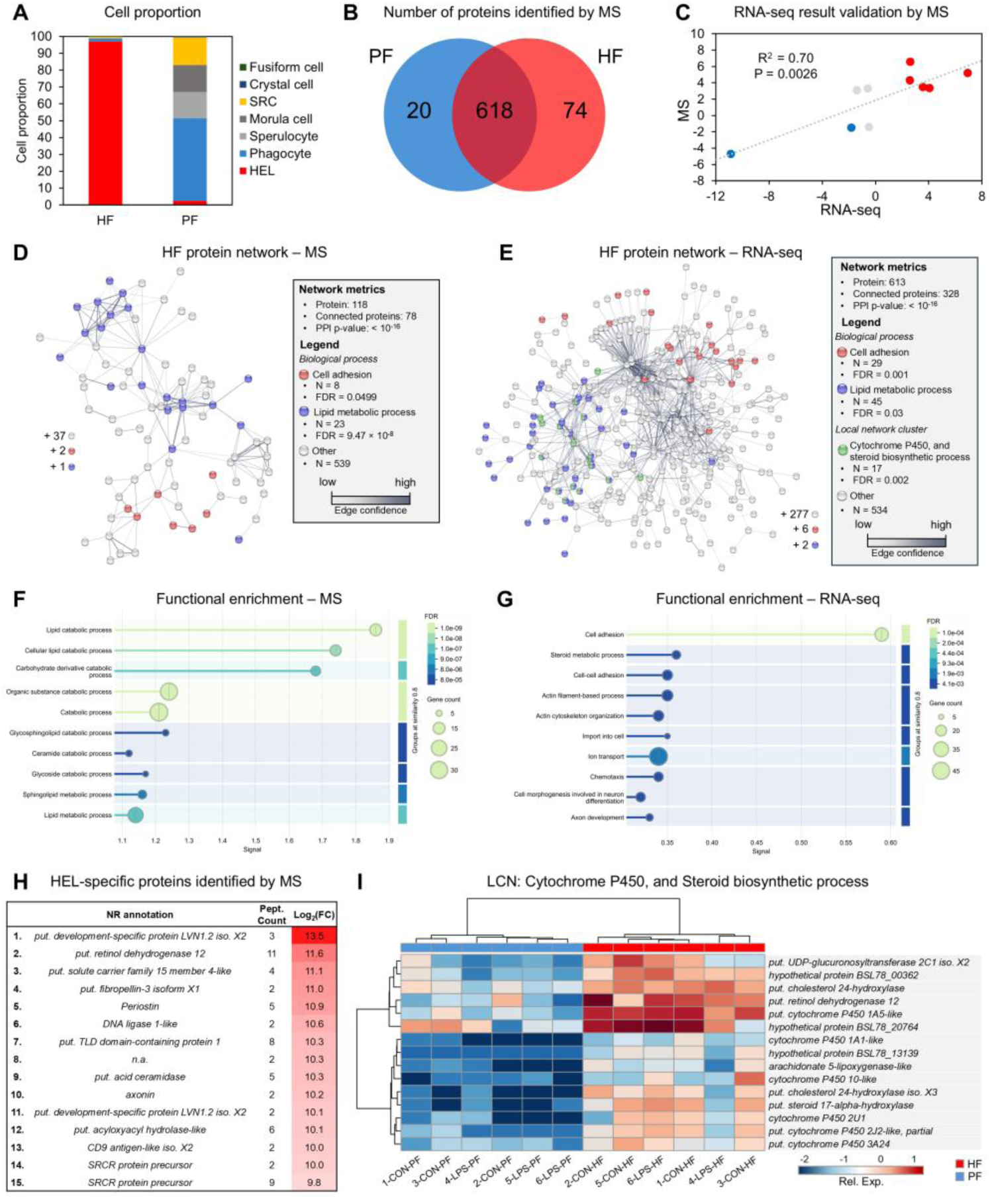
Specific expression of HELs and validation RNA sequencing analyses (RNA-seq) by mass spectrometry (MS). **A.** Proportion of cell types in hydrovascular fluid (HF) and perivisceral fluid (PF) samples analysed by MS. Note that the HF is constituted of 97% of HELs. **B.** Venn diagram of proteins identified by MS between PF and HF. The protein list and associated metrics are shown in **Table S8**. **C.** Relationship between the fold changes in RNA-seq and MS analyses between HF and PF for proteins annotated with a DEG from RNA-seq analysis (PF vs HF) and having only one gene annotation (Blue dots – down-regulated in the HF in the two analyses; red dots – up-regulated in PF in the two analyses; grey dots – conflicting results). The coefficient of determination (R^2^) and the p-value are indicated on the graph (Pearson correlation test). Protein annotations are shown in **Table S8**. **D.** and **E.** String protein network based on proteins with a fold change > 5 in MS analysis and proteins (i.e, translated genes) with a fold change > 2 in RNA-seq analysis, respectively (the legend includes protein network metrics and functional enrichment analysis, including two biological processes shared between MS and RNA-seq networks, and a local network cluster analysis for RNA-seq analysis; results of the string mapping is shown in **Table S9**). **F.** and **G.** The ten most enriched biological processes (based on the gene ontology database) of the protein networks in **D.** and **E.**, respectively. **H.** The fifteen most differentially expressed proteins between PF and HF according to MS analysis. For each protein, the annotation, the peptide count and the fold change (under log2 transformation) are indicated. **I.** A heat map of the differential expression between PF and HF of the proteins in the local network cluster ‘Cytochrome P450 and steroid biosynthesis process’ from the protein network in E, with their gene annotation (Nr annotation).

Parallel to these analyses, a search for genes coding for “globin” and “haemoglobin” was carried out in line with the haemoglobin-based pigmentation hypothesis. While four genes could be identified as coding for globin based on a Blast search against the transcriptome of PF and HF (**Fig. S10A**), none were differentially expressed between the two fluids (**Fig. S10B** and **C**). Moreover, no protein was annotated with one of these genes in the MS analysis, reflecting a low expression level. These results demonstrate that globins are not particularly expressed in HF and HELs.

### 11. Hemocyte-like cells do not produce ROS but express antioxidant enzymes

In order to characterise the function of the HELs in more detail, the redox balance was studied, assuming that the carotenoids contained in the HELs are antioxidant molecules, representing potentially an essential function of these cells. Firstly, a search was carried out for the expressed genes with an antioxidant function, namely superoxide dismutase (SOD), catalase (CAT) and glutathione peroxidase (GPx), among the DEGs between the two fluids. This resulted in one gene annotated as CAT being overexpressed in HF (PF versus HF: log_2_ FC = 3.43; FDR = 0.011) and one gene annotated as SOD overexpressed in PF (PF versus HF: log_2_ FC = -1.87; FDR = 0.041). However, by examining the entire transcriptome, it was possible to identify 8 genes coding for SOD, 5 coding for CAT and 20 coding for GPx expressed in the PF or HF. To study the expression of these genes as a functional entity, the total expression of the different genes was calculated (i.e., the sum of their FPKM value; **Fig. 12A**) and compared between the two body fluids (**Fig. 12B**). This shows that the total expression of the genes is significantly higher in HF (p = 0.0313), which is mainly due to the total expression of CAT, which is also significantly overexpressed in HF (p = 0.0163). On the other hand, the total expressions of SOD and GPx were similar between the two body fluids (P > 0.05). While only one CAT gene showed a significant differential expression between the two fluids, the other four genes encoding CAT appear to follow the same trend (i.e., in each individual, they are more expressed in the HF). Indeed, when examining their expression on a heat map, a relatively good clustering between the two body fluids can be noticed, although the 3-CON-HF sample clusters with 3-CON-PF (**Fig. 12C**). However, it is known from cell counting that this HF sample is the one containing the lowest proportion of HELs (**Fig. 12A**), suggesting a relationship between the proportion of HELs and CAT expression. To verify this, the total expression ratio PF/HF of CAT for each individual was correlated with the proportion of HEL in the HF. This results in a significant correlation (R^2=^ 88.1%; p = 0.0055), meaning that the higher the proportion of HEL in the HF, the lower the PF/HF ratio of CAT expression (**Fig. 12D**). This tends to show that HELs overexpress CAT genes compared to other cell types, but not other antioxidant genes such as SOD and GPx.

**Fig. 12.**
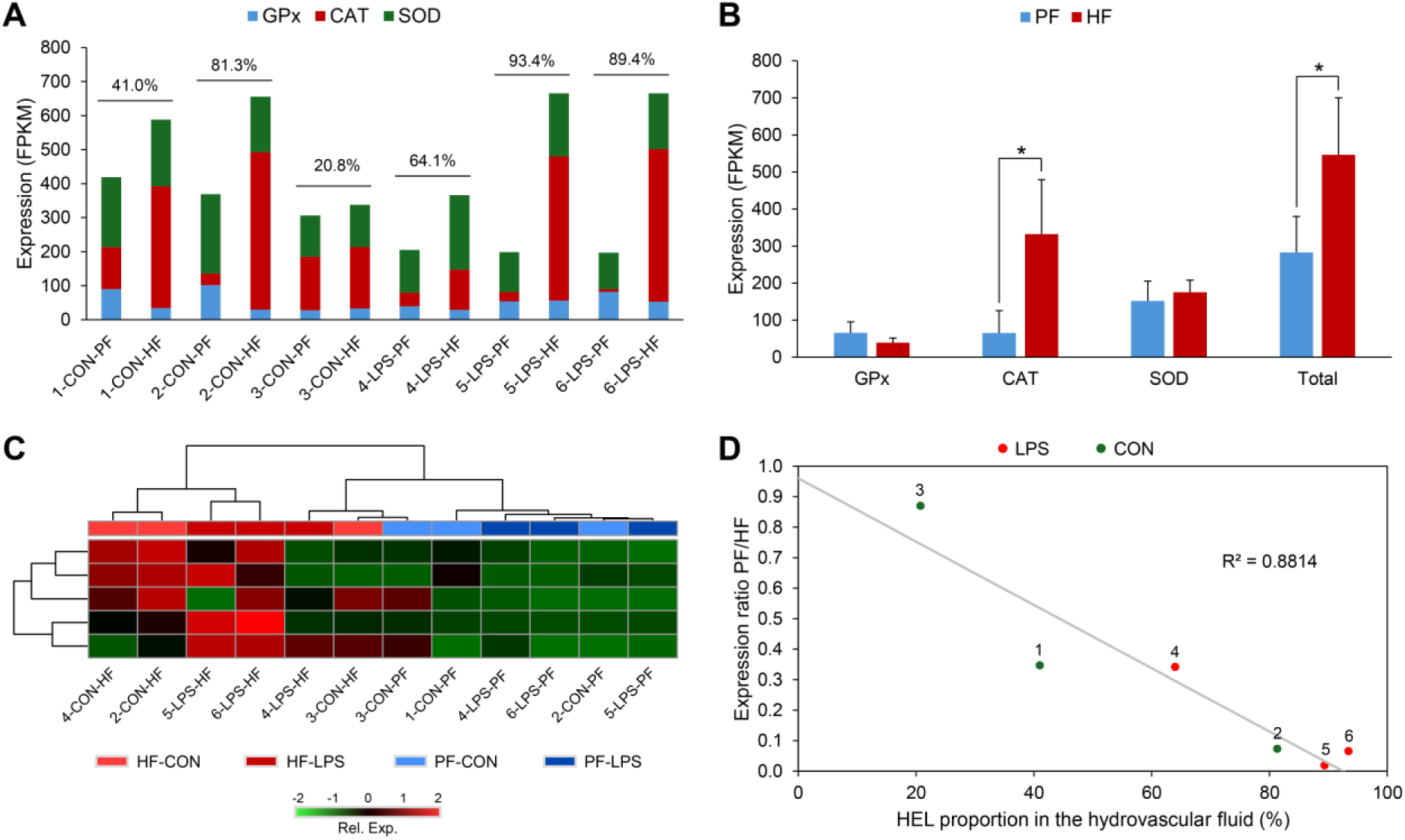
Expression of antioxidant genes in perivisceral fluid (PF) and hydrovascular fluid (HF) of *Holothuria forskali*. **A.** For each individual, the total expression of the three antioxidant gene families in PF and HF (i.e., the sum of the FPKM values of the genes annotated in the gene family). The proportion of hemocyte-like cells (HELs) is shown above the histograms of each individual, suggesting a correlation between catalase expression and the proportion of HELs. **B.** Comparison of total antioxidant gene expression between PF and HF, meaning the sum of expression of genes from the three gene families. Error bars represent the SD (n = 6; Wilcoxon signed rank test; * p-value < 0.05). **C.** Heatmap based on the expression of the five catalase genes expressed in coelomocytes. **D.** Relation of the PF/HF ratio of catalase total expression as a function of the proportion of HELs in the HF (Pearson correlation test). Legend: CAT – catalase; CON – control individual; HF – hydrovascular fluid; GPx – glutathione peroxidase; LPS – LPS-injected individual; PF-perivisceral fluid; Rel. Exp. – relative expression; SOD – superoxide dismutase.

In parallel with the analysis of antioxidant gene expression, ROS production between different types of coelomocytes was studied using the DCFH-DA – a ROS marker. First, ROS-positive cells were visualised using a fluorescence microscope (**Fig. 13A and B**). While the HELs showed high autofluorescence at 495 nm, this was no longer perceptible in the presence of other ROS-positive cells, being highly fluorescent (**Fig. 13A**). The DCFH-DA labelling was confirmed by examining the same samples without DCFH-DA (**Fig. S11**). By examining the different types of cells individually with the same fluorescence intensity, it was also possible to show that ROS production varies considerably between coelomocyte populations: phagocytes and large spherulocytes are strongly ROS-positive; small spherulocytes and SRCs are moderately ROS-positive; Morula cells, crystal cells and HELs are not ROS-positive (**Fig. 13B**). To confirm these results quantitatively, ROS production was studied by spectral flow cytometry using PF and HF cells with and without lipopolysaccharide exposure. This type of cytometry was chosen mainly to manage the high autofluorescence of coelomocytes, and in particular, HELs, which could lead to false-positive results (see gating strategy in the PF and the HF in **Fig. S12**, respectively). In terms of populations, the results showed a similar profile to that obtained previously (see section 8), except that the E population of the HF was divided into two subgroups, namely the E and Ebis populations because in this analysis, we noticed a higher density of highly granular cells (i.e., higher SSC for the Ebis population; **Fig. 13C**). It is known, from the comparison between the two fluids and the sorting experiment, that population A corresponds to HELs (see sections 8 and 9). By comparing the proportion of ROS between the different populations, it is clear that HELs show a lower proportion of positive ROS compared to the other populations (**Fig. 13C**). A statistical comparison was then conducted to compare the proportion of ROS-positive cells and ROS production (based on median fluorescence) between the different populations exposed or not exposed to LPS (**Fig. 13D**). Firstly, no clear difference could be found between the two conditions, neither in terms of the proportion of ROS-positive cells nor in terms of ROS production. Secondly, in the comparison of cell populations, significant differences were observed in the proportion of ROS-positive cells and ROS production (Friedman Χ^2^ = 18.6; df = 5; p = 0.0023 and Friedman Χ^2^ = 16.7; df = 5; p = 0.0051, respectively). For both measures, HELs have the lowest value with 8.4 ± 8.9% of ROS-positive cells and a median fluorescence of 246 ± 237. These are significantly different from all other populations, except for population E in the PF, corresponding to small cells that also had a relatively low level of ROS-positive cells and ROS production (25 ± 23% and 715 ± 888, respectively). In comparison, the most ROS-positive populations, namely population B in the HF and population D in the PF, have a proportion of ROS-positive cells of 87.9 ± 15.2% and 61 ± 28% and ROS production of 13.936 ± 12.813 and 8.828 ± 9.525, respectively. In addition, cell viability was investigated between the different conditions using propidium iodide staining. However, no difference was revealed, neither between the conditions nor between the cell populations (**Fig. S13**), with an overall low mortality proportion ranging from 1.2 ± 0.5% for population C to 14 ± 11% for population Ebis.

**Fig. 13.**
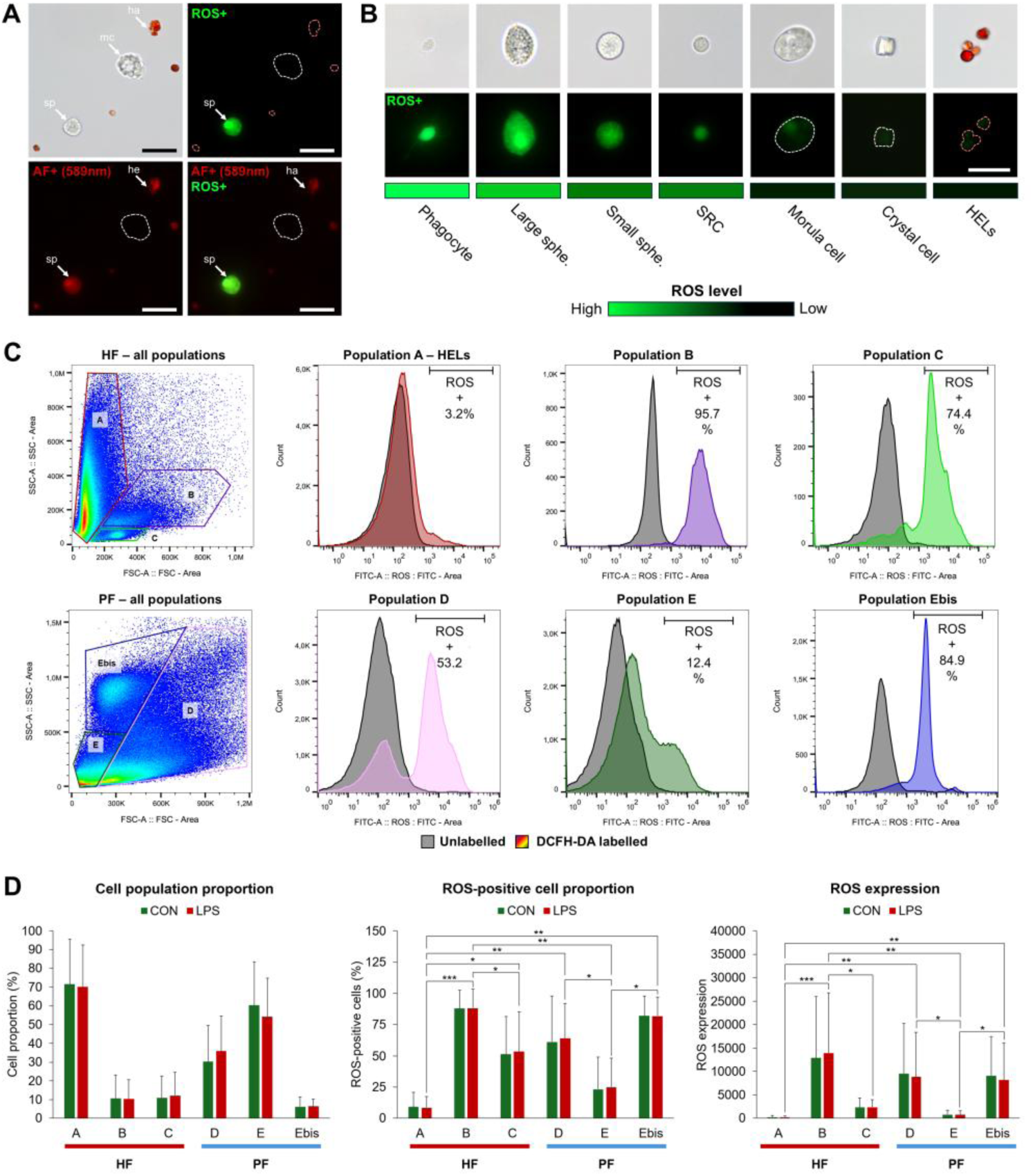
Study of the production of reactive oxygen species (ROS) by coelomocytes from hydrovascular (HF) and perivisceral fluids (PF) in *Holothuria forskali* using DCFH-DA labelling. **A.** DCFH-DA labelling of HF coelomocytes: while HELs are autofluorescent at 589 nm, the autofluorescence of HELs is no longer visible with the excitation wavelength of DCFH-DA (495 nm) when there are ROS-positive cells around (he – hemocyte-like cell; mc – Morula cell; sp – spherulocyte; a negative control is shown in **Fig. S11**). The scale bar represents 20 µm. **B.** DCFH-DA labelling of different types of coelomocytes photographed with the same intensity; the lighter green a cell is, the higher the level of ROS (the dotted lines in A and B indicate where the localisation of the cell is when they are not visible). Scale bars represent from left to right: 20 µm, 14 µm, 12 µm, 10 µm, 15 µm, 16 µm and 10 µm, respectively. **C.** ROS production quantified between different coelomocyte populations using spectral flow cytometry: the graph on the left shows the population gating of coelomocytes from HF and PF, histograms on the right show, for each population, the distribution of cells as a function of fluorescence intensity (black – no DCFH-DA labelling; coloured – same population with a DCFH-DA labelling; the proportion of ROS-positive cells is indicated on the graph based on the defined threshold).The gating strategy is shown in **Fig. S12**. **D.** Comparison of the proportion of ROS-positive cells and ROS expression between the different populations and between lipopolysaccharide-exposed and control samples based on the quantification method in C. The graph on the left shows the proportion of each population; the middle and right graphs show the proportion of ROS-positive cells and ROS expression (based on the median fluorescence intensity), respectively. Results are formulated as means with error bars representing the SD (n = 4 in LPS-exposed samples and n = 3 in control samples; Friedman test; p-value: * < 0.05; ** < 0.01; *** < 0.001).

## Discussion

### 1. Coelomocyte types are not equally distributed in the coelomic fluids

The wide diversity of coelomocyte morphotypes in sea cucumbers suggests a complex immune system in which each cell type fulfils distinct functions in immunity. However, very little information exists on these respective functions. With their distinctive coelomic compartment containing the hydrovascular fluid (HF) and the perivisceral fluid (PF), sea cucumbers offer a promising model for studying the distinctive function of these body fluids as reflected by their respective cellular composition. In *H. forskali*, 11 main morphotypes could be distinguished under light microscopy (including 5 subtypes and 2 uncertain types), with seven main cell types generally observed and included in cell counts **(Fig.1)**. These results confirm previous studies on other sea cucumber species (*e.g.*, Xing et al., 2008 for *A. japonicus*; Wambreuse et al., 2025 for *H. scabra*; Queiroz and Custódio, 2024 for the review) and notably the presence of a large number of small reddish cells – HELs – notably similar to the recent description of hemocytes in the order Holothuriida (Caulier et al., 2024). Our primary results demonstrated that coelomocyte types are not equally distributed between PF and HF **(Fig. 2)**, with a conspicuous difference in the almost exclusive presence of HELs in the HF, where they are generally the predominant type. In contrast, spherulocytes and crystal cells were more represented in the PF than in the HF. While differences in the concentration of cell types between the two fluids have rarely been investigated (e.g., in Li et al., 2013), certain common patterns are shared between species: crystal cells were similarly found to be more prevalent in the PF of *H. scabra* (Wambreuse et al., 2025), and the so-called hemocytes were found to be restricted to the HF in many species including *Allothyone mexicana*, *Bohadschia argus*, *Cucumaria frondosa*, *Eupentacta quinquesemita, Holothuria atra*, *H. scabra*, *H. forskali* and *Sclerodactyla briareus* (Caulier et al., 2024, 2020; Manwell and Baker, 1963; Roberts, 1980; Roberts et al., 1984). These differences in the cellular composition of the two fluids, mainly represented by the remarkably high concentration of HELs in the HF, suggest divergent functions associated with their respective cellular composition.

### 2. Coelomocytes of the two body fluids are immunocompetent

The activity and behaviour of the cells were followed during immune response activation to study the function associated with the different cell types. Although fluid extraction does not truly represent immunological stress, we noticed that when no anticoagulant solution was added, the cells initiated amoeboid-like movements and formed aggregates (**Videos S2-S4**). In this case, the immune activation could be triggered by the recognition of damage-associated molecular patterns released during dissection, such as intracellular proteins or extracellular nuclear DNA (Schaefer, 2014). As these cellular behaviours were characteristic of some cell types, such an approach can be informative about the function of different cell types. According to our observations, the process begins with the activation of phagocytes that adhere to the slide and extend their pseudopodia to recruit other types of cells. This recruitment is thought to be induced by the release of humoral factors such as cytokines and agglutinins, which we have identified as being widely expressed in coelomocytes (**Fig. 4**). These could activate the mobility of other cells and attract them by chemotaxis. Spherule cells, in particular, showed a significant degree of mobility through a characteristic amoeboid movement based on the formation of spherical protrusions at the apex of the cell, followed by an undulation, giving them a figure-eight shape (**Fig. S1B**; **Video S1**). This particular mobility is reminiscent of the bleb-driven mobility, a particular amoeboid movement based on an increase in the hydrostatic pressure inside the cell and distinct from pseudopod-based mobility, which is thought to confer to the cells a “swimming ability” (Paluch et al., 2016). This ability could be useful for a fast circulating cell recruitment to the site of infection. When they encountered a newly formed aggregate, some spherule cells could carry out sudden cell lysis, releasing numerous granules (**Video S1**). Previous reports suggested that spherulocytes or Morula cells could perform similar cell lysis to release granules and amorphous material based on morphological analysis during the formation of brown bodies following the injection of foreign material (Dybas and Fankboner, 1986; Jans et al., 1995). The released granules could also explain the observation of minute corpuscles in several studies (Queiroz and Custódio, 2024). Furthermore, our results revealed that early aggregates, by becoming increasingly compact and merging, can move autonomously on the slide, as observed by Taguchi et al. (2016), who proposed that this "crawling ability" could be useful for the aggregate to migrate to the sites of injury. In HF, we were able to confirm that HELs also participate in this process. Although they are not mobile, they appear to be highly interactive when they encounter another cell or a cell aggregate. The aggregation process is known to be an essential immune mechanism in echinoderms that aims to encapsulate foreign matter and pathogens (Caulier et al., 2020; Majeske et al., 2013; Taguchi et al., 2016) and is thought to be responsible for the formation of coloured bodies in the different echinoderm classes (Jobson et al., 2022). In *H. forskali*, we were able to easily observe large red bodies in the Polian vesicle and buccal tentacles, similarly to the description of these red bodies in *C. frondosa* (Caulier et al., 2020). Overall, these results confirm that aggregation is an important immune mechanism in sea cucumbers and provide new information on the first stages of this phenomenon involving the HELs.

Immunostimulation with lipopolysaccharides indicated that only the concentration of HELs in the HF was significantly increased one day after the injections, which suggests that this type of cell has a specific function in the immune response (**Fig. 3**). Changes in the coelomocyte population have already been studied in *H. glaberrima* and *H. scabra* following injections of various Pathogen-Associated Molecular Patterns (PAMPs), including lipopolysaccharides (Ramírez-Gómez et al., 2010; Wambreuse et al., 2025). Although these species show different patterns in the modifications of the cell populations, both displayed a clear response. Furthermore, the timing of the immune response is important to take into account and could have prevented the detection of changes in other cell populations in our study. In *C. frondosa*, for example, the concentration of coelomocytes was monitored at different time points after various stresses, including the presence of a predator or an injured conspecific (Hamel et al., 2021). In this study, the peaks of phagocytes and Morula cells preceded the peak of hemocytes and began to decrease three hours after the stress. Therefore, we cannot exclude that an increase in the concentration of other cells occurred earlier and that only a peak in hemocytes was visible at that time. Moreover, the high responsiveness of these cells to many non-pathogenic stressors (Jobson et al., 2021), including handling (Caulier et al., 2020), makes it difficult to obtain a real negative control. Although an acclimation period of a least two weeks was respected before any experiment, stress related to the detention of individuals might have influenced the concentration of some cell type responding to non-specific stress (i.e., not necessarily pathogenic). In any case, our result showed that HELs increased after one day, suggesting, in combination with the study of Hamel et al. (2021), that HELs act later than other cells in the immune response.

The immune response to LPS injection was also assessed at the level of gene expression between the two body fluids (**Fig. 4**). Both show a clear transcriptional response, with many physiological pathways related to the immune system that were differentially expressed and shared between them, including “NOD-like receptor signalling pathways” or “Toll and Imd signalling pathways”, which are important pathogen recognition pathways (Chen et al., 2021, Chiaramonte and Russo, 2015; Sun et al., 2013). Other pathways involved in immune mechanism regulation include apoptosis, necroptosis, the “IL-17 signalling pathway”, or the “NF-kappa B signalling pathway”. These pathways are consistent with previous studies investigating the gene expression of coelomocytes following exposure to PAMPs, notably in *H. leucospilota* and *H. scabra,* which showed a notably high expression of cytokines and NOD-like receptor genes (Wambreuse et al., 2025; Wu et al., 2020). In contrast to these pathways, some were not enriched in the same way between the two body fluids, including the “PI3K-Akt signalling pathway” and “RIG-I-like receptor signalling pathway” that were respectively more differentially expressed in the HF and the PF during the immune response (**Fig. 5**). “PI3K-Akt signalling pathway” is notably involved in cell proliferation and epithelial-mesenchymal transition (Xu et al., 2015). During an immune response, it was proposed that hemocytes, initially marginated to the inner wall of hydrovascular tissues, could detach to enter the cell suspension (Caulier et al., 2020). This is consistent with our observation of a large number of HELs on the inner membrane of the Polian vesicle (**Fig. 7**) and their increase in concentration following the immunological stress that only occurred in the HF. Differential expression of this pathway could, therefore, reflect the demargination process during the immune response (Caulier et al., 2020; Xu et al., 2015). “RIG-I-like receptor signalling pathway” is a pathogen recognition receptor pathway that has an important function in innate immunity and can notably cooperate with Toll-like receptor signalling (Loo and Gale, 2011). Overall, gene expression analyses following the injection of LPS demonstrated that coelomocytes of both fluids are highly responsive cells expressing a large diversity of immune genes (**Fig. 4**), including numerous homologues of vertebrate immune genes (Hibino et al., 2006; La Paglia et al., 2025).

### 3. Unexpected characteristics of Hemocyte-like cells

To study the function of the HELs in more detail, we decided to take advantage of the significant enrichment of these cells in the HF to compare them with the coelomocytes of the PF. Surprisingly, we discovered that the HF coelomocytes contained an enormous quantity of carotenoids that correlated with the number of HELs in the samples analysed (**Fig. 6**). Furthermore, by performing the same analysis on FACS-purified HELs, we unequivocally demonstrated that HELs were the pigment-carrying cells (**Fig. 9**). Reddish-pigmented cells of sea cucumbers have historically been considered and designated as "hemocytes" because of their haemoglobin content (Crescitelli, 1945; Kindred, 1924). Consequently, it has been suggested that their function is to oxygenate tissues by participating in the transport of oxygen (Baker and Terwilliger, 1993; Smith, 1981). This hypothesis on the function of hemocytes has been widely accepted for over a century and has been posited as a universal characteristic of hemocytes in sea cucumber species. In order to characterise this newly described cell type in more detail, various analyses were carried out and are discussed below.

First of all, our results showed that the HELs were located mostly in the HF, where they were generally observed in large red bodies or formed small aggregates of about ten cells that were generally marginated at the inner membrane of the Polian vesicle (**Fig. 7**). While most HELs are small, we observed some that were larger and granular in appearance. Furthermore, given that some HELs were minute (less than 2 µm), we hypothesised that s ome HELs could instead result from the release of small anucleate corpuscles by larger ones. This hypothesis is consistent with previous observations of hemocytes forming cytoplasmic protuberances that detached as small spheroids before the entire cell underwent cytolysis (Fontaine and Lambert, 1973; Hetzel, 1963). It should be noted that Hetzel (1963) also mentioned that some of these vesicles contained refractive yellow granules, suggesting a possible role in the release of pigments or metabolites. This process of cytolysis is also reminiscent of platelet production in vertebrates, which results from the cellular lysis of megakaryocytes (Patel, 2005). Although further research is needed to confirm this fragmentation ability, this potential mechanism of HEL fragmentation might enhance the organism ability to promptly respond to physiological needs and facilitate the distribution of these corpuscles across tissues or within cellular aggregates to carry out various biological processes.

Another characteristic of the HELs was their strong autofluorescence (**Fig. 8**). This autofluorescence was attributed to the high concentration of carotenoids inside the cell, as it has been shown that astaxanthin-producing green algae also exhibit a strong autofluorescence (Ota et al., 2018) that was notably used in flow cytometry analyses (Chen et al., 2017). In the present study, we also took advantage of this autofluorescence using spectral flow cytometry to determine the best strategy for isolating these cells by FACS (**Fig. 9**). FACS has also been used to sort the red spherule cells in sea urchins according to their autofluorescence, which has made it possible to better characterise the pigment contained in these cells (Hira et al., 2020; Smith et al., 2019). While sea urchin red spherule cell pigments are rather naphthoquinone (Brasseur et al., 2018), the strategy for isolating pigmented cells based on autofluorescence has now also proven to be effective in sea cucumbers with carotenoids, which offers good prospects for the use of this technique in other echinoderm species in the future.

### 4. Carotenoid diversity and acquisition in sea cucumbers

HPLC analyses have shown that the carotenoids contained in HELs are mainly composed of canthaxanthin (∼55%) and astaxanthin (∼45%) and, to a lower proportion, echinenone (<5%) (**Fig. 6**). Astaxanthin and canthaxanthin are keto-carotenoids found in a wide variety of aquatic organisms, including shrimps, crabs, crayfishes and fishes (Czeczuga, 1975; Czeczuga-Semeniuk and Czeczuga, 1999; Maoka, 2011; Shahidi and Synowiecki, 1991). In sea cucumbers, a wide variety of carotenoids have been identified, mainly in the integument, gonads or other viscera (e.g., Bandaranayake and Rocher, 1999). These pigments include astaxanthin, canthaxanthin and echinenone, in accordance with our results, as well as β-carotene, phoenicoxanthin and cucumariaxanthin, which were not detected in *H. forskali* (Bandaranayake and Rocher, 1999; Matsuno and Tsushima, 1995; Morrison et al., 2024). Recently, David et al. (2023) studied the diversity of carotenoids in the gonad of *H. forskali*, showing a very similar composition in the same types of carotenoids but with a much lower concentration compared to the Polian vesicle sample (which contained a large amount of HELs). This suggests that the carotenoid profile within a species is relatively constant between the different tissues. As for interspecies differences, different proportions of carotenoids are generally observed. For example, Matsuno and Tsushima (1995) identified three new carotenoids, namely cucumariaxanthines A, B and C, which were specific to the order Dendrochirotida, suggesting a possible taxonomic specificity in their distribution. This difference in carotenoid composition could thus explain why certain coloured coelomocytes, described as hemocytes, may have a different colouration. For example, it has been shown that the hemocytes in *H. scabra* are brownish (Caulier et al., 2020; Wambreuse et al., 2025), which was initially attributed to lower oxygen concentrations in the body fluid of this tropical species, resulting in a reduced form of haemoglobin (Caulier et al., 2024). Following our results, it could reasonably be suggested that this colour is due instead to the presence of different carotenoid types. Further research is underway to identify the nature of the HEL pigmentation in various species of sea cucumbers belonging to different taxonomic groups and with different ecologies to better determine which factors shape this pigmentation.

Our results indicate that the carotenoid content of HELs varies between 1.4 and 30 µg per 10^6^ cells (**Fig. 6 and 9**). This range is not far from that reported for the green algae *Haematococcus pluvialis*, which can reach higher values, such as 80 µg per 10^6^ cells, but are larger cells (Orosa, 2005). Although these values cannot be compared to those of metazoan tissues, generally measured as percentage of the mass, a comparison of the results obtained with the Polian vesicle samples reveals a much higher concentration than those found in salmon or krill, known to have a high carotenoid content (Tan et al., 2024). Metazoans, with a few exceptions, cannot synthesise their carotenoids *de novo* and must acquire them through their diet (Maoka, 2011). The transformation and storage of such a quantity of carotenoids can be costly for the metabolism and takes a certain amount of time to accumulate them (Tan et al., 2024). Consequently, the possibility of storing these carotenoids in specialised cells for long-term use could constitute a significant advantage. In echinoderms, carotenoids were notably found in different species and organs of sea stars (Asteroidea) (e.g., Lourtie et al., 2024; Williams et al., 2025), where they are supposed to help cope with environmental stresses (Williams et al., 2025), however, it is still unclear if they are acquired through their diet (Lourtie et al., 2024). In many holothuroid species, including *H. forskali,* a high concentration of carotenoids was found in gonads, which was attributed to a protective effect for the gametes and a potential source of energy for the embryos (Bandaranayake and Rocher, 1999; David et al., 2023). However, the mechanism of such an accumulation of carotenoids in the gonads remains unclear. David et al. (2023) hypothesised that the acquisition of carotenoids could occur during gonadal maturation through diet. However, they could not find any difference in carotenoid content throughout the year, which raised questions about the source and allocation of these carotenoids. They concluded that they could originate from the reabsorption of carotenoids in the spent gonadal tubules (after spawning). During our experiments, large quantities of HELs and coelomocyte aggregates were observed in the dehiscent gonadal tubules (unpublished results), which suggests that these circulating cells could play a central role in the transport and recycling of carotenoids. Furthermore, the study of the composition of the diet in the intestine of *H. forskali* reveals mainly the presence of fucoxanthin, which would come from brown algae (David et al., 2020). This type of carotenoid is not a direct precursor of keto-carotenoids such as canthaxanthin and astaxanthin (Maoka, 2011). However, a recent study testing different rearing methods for *H. forskali*’s juveniles in co-culture with oysters and abalones revealed the presence of β-carotene and, to a lesser proportion, canthaxanthin (David et al., 2024). These pigments are precursors of astaxanthin, which means that under certain conditions (Maoka, 2011), *H. forskali* could directly accumulate these precursors. Although it remains unclear whether *H. forskali* can acquire carotenoids throughout its life, it can be reasonably assumed that the HELs could regulate the use of carotenoids by acting as long-term storage inside the hydrovascular system. It is also important to note that, in contrast to other carotenoid-rich organs such as gonads or digestive tract, the Polian vesicle(s), and the hydrovascular system in general, are not expelled during self-evisceration (Li et al., 2018), allowing to keep a carotenoid stock even during the regeneration process.

### 5. Differential expression between the two body fluids provides insight into the function of HELs

Our results showed that gene expression between the two body fluids tends to be more divergent under immunostimulation conditions than under immunoquiescent conditions (**Fig. 5 and 10**), which coincides with the increase in the proportion of HELs that only occurs in the HF of immunostimulated individuals (**Fig. 3**). Thereby, many pathways are not shared in the comparison of PF versus HF between immunostimulated and immunoquiescent condition analyses. Furthermore, it is interesting to note that a large proportion of these pathways were not related to the immune system, but rather to the endocrine and digestive systems, with a higher overall number of up-regulated genes in the HF, suggesting that this fluid is also involved in the transport of nutrients and regulation of physiological processes through hormonal control. By counting the proportion of coelomocyte types in the sequenced samples, it was possible to correlate differential gene expression with the proportion of HELs in the HF, meaning that a large part of these functions can be attributed to HELs (**Fig. 10**). This approach, combining cell counting and comparative transcriptomics, proved to be very informative in terms of cell function. Moreover, thanks to this, we were able to select samples where the concentration of spermatozoa was reasonable in comparison with the proportion of coelomocytes, which would not have been possible without checking the cellular composition of our samples before extracting the RNA. Such gamete contamination might explain why some differences can be observed in the expression of immune genes in coelomocytes between males and females (e.g., this study **Fig. S4**; Pérez-Portela and Leiva, 2022). For these reasons, we strongly encourage determining the cellular proportions when considering omic analyses of sea cucumber coelomocytes.

On the basis of the genes and proteins that were upregulated in HF, we were able to identify two broad biological functions characteristic of HELs, namely "cell adhesion" and "lipid metabolic processes" (**Fig. 11**). The processes in cell adhesion encompass proteins involved in the interaction with the extracellular matrix and interaction with other cells (Ince et al., 2019; Meager, 1999). This process corresponds to the characteristics of HELs, which are often observed in small aggregates or marginated at the wall of the Polian vesicle (Caulier et al., 2020; Ince et al., 2019), probably involving proteins such as integrins, cadherins and laminins that have been identified as overexpressed in HF (**Table S9**). As for the "lipid metabolic processes", they can reasonably be attributed to the metabolism of carotenoids, which are lipidic molecules synthesised from tetraterpene precursors (Stra et al., 2023). We have been notably able to identify candidate proteins involved in carotenoid metabolism, particularly in the pathway "Cytochrome P450, and steroid biosynthetic process". In this metabolism, six proteins have been annotated as Cytochromes P450 and are overexpressed in the HF (**Fig. 11**). These proteins are highly conserved enzymes with a function in the biosynthesis and detoxification (Stegeman, 2000). However, it has been shown that they can have a bifunctional activity by also playing the role of carotenoid-ketolase, which transforms “conventional” carotenoids such as β-carotene into ketocarotenoids such as canthaxanthin and astaxanthin. This property was first demonstrated in birds, in which the expression of a gene belonging to the cytochrome P450 family, namely CYP2J19, was identified as responsible for the red-beaked phenotype (Mundy et al., 2016). Later, a similar carotenoid-ketolase function of cytochrome P450 genes was reported in several lineages, including spider mites, shrimps and scallops (Wang et al., 2022; Weaver et al., 2020; Wybouw et al., 2019) but, to our knowledge, not in echinoderms. It has been suggested that the acquisition of this bifunctional activity of cytochrome P450 would result from the evolutionary convergence of these enzymes under different types of selective pressure. Interestingly, disruption in the expression of cytochromes P450 can lead to yellow-brown phenotypes, as illustrated by spider mites (Wybouw et al., 2019). Given that some species of sea cucumber have brown HELs, such as *H. scabra*, it could be hypothesised that these species have lost or never acquired the bifunctional Cytochrome P450 enzyme. However, further research is needed to determine whether this difference in pigmentation is due to genetic variation or different ecological parameters, such as different food type resources. In addition to Cytochrome P450, we have been able to identify genes coding for "retinol dehydrogenase 12" and "UDP-glucuronosyltransferase", which are also thought to play a role in carotenoid metabolism. Retinol is a derivative of vitamin A, synthesised from the β-carotene precursor similar to canthaxanthin and astaxanthin (Lidén and Eriksson, 2006; Maoka, 2011). UDP-glucuronosyltransferase is an enzyme also involved in detoxification, in particular by catalysing the transfer of glucuronic acid to small lipophilic molecules (Zheng et al., 2016). It has recently been shown that it is differentially expressed in the integument between different colour morphotypes of *A. japonicus*, suggesting that this gene may also have a role in regulating integument pigmentation through the carotenoid metabolism (Liu et al., 2024). More broadly, Cytochrome P450 metabolisms have been reported in several studies investigating colour phenotypes in sea cucumbers without making a direct link with the carotenoid-ketolase activity highlighted in other lineages (e.g., Han et al., 2025; Liu et al., 2024). These results suggest, therefore, that this metabolic pathway plays a key role in sea cucumber pigment processing.

In addition to these enzymes, we listed the fifteen most enriched proteins in HELs by mass spectrometry (**Fig. 11**). Among them, two were annotated as “putative developmental specific protein LVN1.2 isoform X2”. Surprisingly, this protein has been reported as a marker of endodermal tissues in sea urchin embryos (Wessel et al., 1989) and has also been overexpressed in the intestine following heat stress in sea cucumbers (Xu et al., 2016). While further research is ongoing to identify the origin of HELs, this expression suggests that these cells might have an endodermal origin, unlike other types of coelomocytes (Li et al., 2019). Overall, it should be emphasised that the proteins discussed here represent only a minor proportion of the proteins identified in these pigmented cells, but that this dataset could help to identify this cell type in other studies and other species, particularly with regard to single-cell RNA datasets in sea cucumbers.

### 6. Function of Hemocyte-like cells in the immune response regulation through the redox balance

Beyond their function in pigmentation, it has been shown that carotenoids play several roles in photoprotection, reproduction and immunity (Lamare and Hoffman, 2004; Tan et al., 2020; Tsushima and Matsuno, 1997; Williams et al., 2025). These are notably enabled by their distinctive structural composition, incorporating a hydroxyl group and a carboxyl group, which give them high antioxidant properties (Brotosudarmo et al., 2020). Among these molecules, astaxanthin has been identified as the carotenoid with the most potent antioxidant properties (Galasso et al., 2017; Miki, 1991). In addition, carotenoids have also been identified as antimicrobial molecules (Jeyachandran et al., 2020; Karpiński et al., 2021). Therefore, HELs, with their high concentration of carotenoids and astaxanthin in particular, could be considered highly antioxidant and antimicrobial immune cells.

In the same vein, we have also demonstrated that HELs contain fewer reactive oxygen species (ROS) than other types of coelomocytes (**Fig. 13**). ROS are highly reactive compounds that can be released during the immune response to neutralise pathogens in a process called respiratory burst (Biller and Takahashi, 2018). However, an excessive concentration of ROS can also be harmful to the host’s tissues, hence the importance of regulating the redox balance during the immune response (Tan et al., 2020). In addition to their high concentration of carotenoids, we have shown that HELs also overexpress catalase (**Fig. 12**), an enzyme that reduces hydrogen peroxide into water and oxygen (Tan et al., 2020). However, surprisingly, the superoxide dismutase, an enzyme that reduces superoxide into hydrogen peroxide (Biller and Takahashi, 2018), is not overexpressed in HELs. We believe that carotenoids could play this role instead of superoxide dismutase, rendering the overexpression of this enzyme useless. Such competitive activity between carotenoids and antioxidant enzymes has been studied in several invertebrates (Babin et al., 2015; Dhinaut et al., 2017; Gostyukhina et al., 2013), particularly in *Gammarus pulex*, in which the uptake of carotenoids decreased the concentration of ROS and the expression of superoxide dismutase, but increased the expression of catalase (Babin et al., 2015).

In contrast to HELs, our results showed that certain populations of coelomocytes, including phagocytes and spherulocytes, were strongly positive for ROS (**Fig. 13**). Although this level of ROS was not higher in the cells exposed to LPS, we believe that harvesting the fluids and maintaining the cells in short-term culture may have led to ROS production in the control samples as well, thereby masking any signal of the immunostimulation. Beyond their immune function, ROS have also been described as key mediators of regeneration processes in metazoans (Vullien et al., 2025). Moreover, it has been shown that coelomocytes are recruited to the site of injury during wound healing or intestinal regeneration (Medina-Feliciano et al., 2025; San Miguel-Ruiz and García-Arrarás, 2007). Lacouth et al. (2024) even reported that among coelomocytes, phagocytes and spherulocytes were the first cell types to be recruited to the site of the lesion. Based on our results, we believe that these cell types could constitute a first line of ROS-producing cells aimed at activating regeneration processes such as cell proliferation at the lesion site. Other types of cells containing a lower quantity of ROS, such as Morula cells and HELs, could ultimately be recruited to balance this effect. Furthermore, it can be hypothesised that the high carotenoid content of sea cucumbers could be an adaptation to their extraordinary regenerative capacities, balancing the need for ROS that could be regulated in the tissues by coelomocytes.

Given that hemocytes participate in the formation of large red bodies, previous studies have suggested that the haemoglobin contained in these cells could release ROS in the aggregate to neutralise pathogens (Caulier et al., 2020). Although we could confirm the involvement of HELs in the formation of red bodies in *H. forskali*, our results tend to demonstrate the opposite. We have therefore proposed a new model explaining the function of HELs, consisting of a highly antioxidant immune regulator (**Fig. 14**). During a bacterial infection or an injection of lipopolysaccharides, phagocytes and spherulocytes begin to aggregate, potentially producing ROS within the newly formed aggregate to neutralise the pathogenic agent (**Fig. 14A-C**). In a second phase, demargination and potential production of HELs occur in the tissues of the hydrovascular system, with possible fragmentation of the larger HELs to release small spheroids and promote their distribution (**Fig. 14D**). Two possibilities are then proposed: these cells could be recruited at the site of the infection or, more unlikely, the early aggregates could migrate in the hydrovascular system (**Fig. 14E**). Thanks to their high adhesive properties, HELs could form a coating around the early aggregate which, given their high antioxidant properties, would constitute an antioxidant shell. In this way, HELs could restrict ROS production to the aggregate by carotenoids and catalase expression, preventing reactive compounds from damaging the host’s tissues (**Fig. 14G**). The resulting coloured aggregates would finally merge to form large coloured bodies (i.e., larger and more compact aggregates) that might be excreted from the body through the cloaca as proposed by Caulier et al., 2020.

**Fig. 14.**
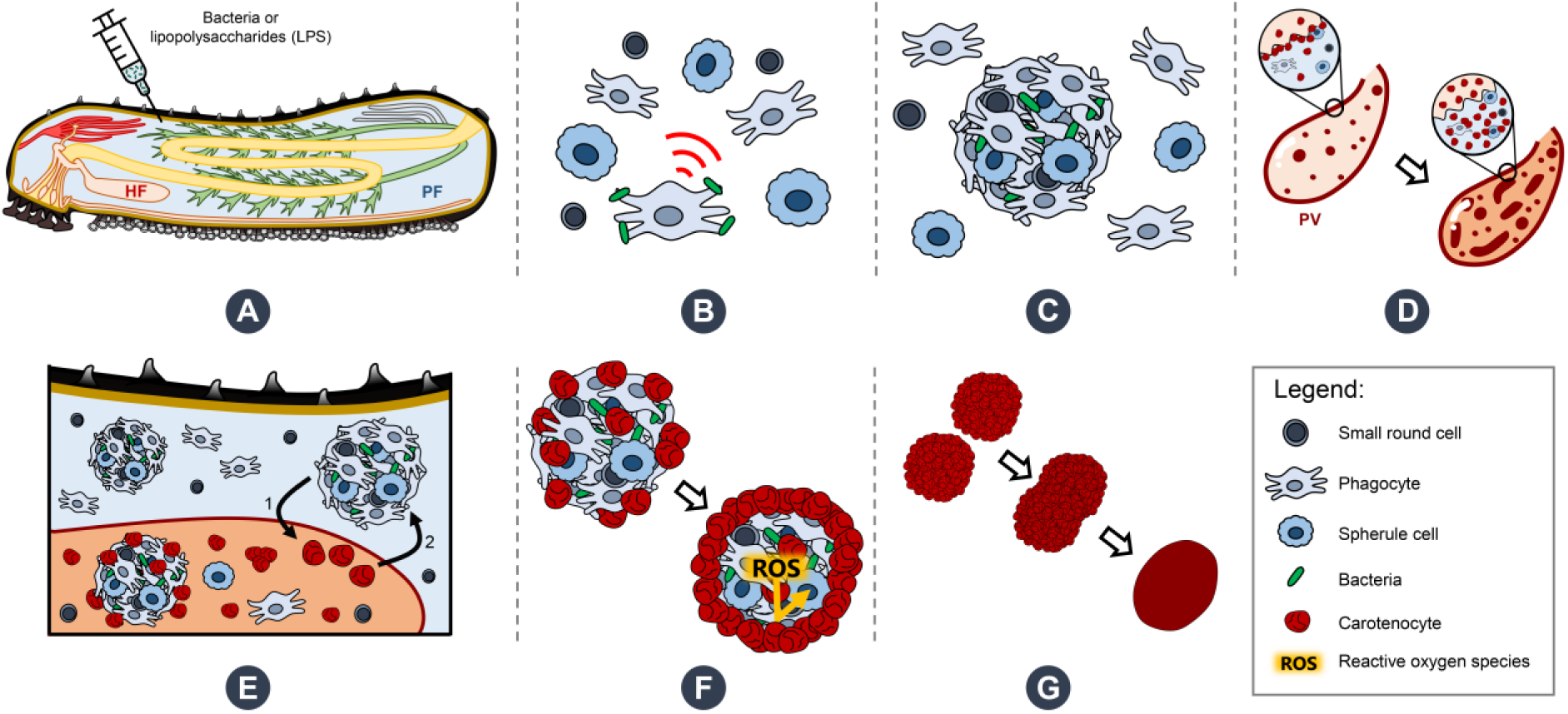
Proposed model illustrating the function of hemocyte-like cells (HELs) – renamed carotenocytes according to our results – in the immune response of *Holothuria forskali* and potentially other sea cucumber species. **A**. The organism is immunostimulated either by bacterial exposure or lipopolysaccharide injections. **B**. The immune response is initiated by phagocytes, which start the aggregation process in perivisceral and hydrovascular fluids (PF and HF, respectively). **C**. Formation of early aggregates occurs through the recruitment of other coelomocytes, leading to the neutralisation of pathogens. **D**. Releasing of marginated HELs, with potential production and fragmentation of HELs in the hydrovascular fluid. **E.** HELs encounter the aggregates via two possible pathways: either HELs migrate toward the site of infection, or the aggregates migrate toward the hydrovascular system after the pathogen entrapment. **F**. Aggregates are covered by a layer of HELs, which form an antioxidant shell to mitigate the production of reactive oxygen species (ROS) within the aggregate. **G**. The red aggregates gather and progressively compact to form large red bodies that are stored in the Polian vesicle (PV) for possible recycling of carotenoids and/or eventual excretion.

### 7. Hemocyte-like cells are sea cucumber pigmented coelomocytes – carotenocytes

Finally, with all the evidence that HELs do not contain haemoglobin but carotenoids, we have proposed to rename these cells “carotenocytes”. Although further research is needed to assess the distribution of these cells in sea cucumbers, we believe that a large proportion of the hemocytes **–** HELs that have been described over the past century do not contain haemoglobin but carotenoids. However, we do not totally exclude the potential presence of true haemoglobin-containing cells in sea cucumbers, particularly in certain burrowing species living in low-oxygen environments (e.g., *Paracaudina chiliensis*; Baker and Terwilliger, 1993). The existence of carotenoid-carrying cells has previously been documented in other marine animals, including fish and crustaceans, in which they constitute erythrophores (carotenoid chromatophores) serving the external colouration (Fingerman, 1965), but no example of circulating carotenoid-carrying cells in a metazoan was found in the literature. Moreover, Xing et al. (2023) recently discovered different pigmented cells in sea cucumbers, namely the quinocytes and melanocytes in the integument of *A. japonicus*, but to our knowledge, our study is the first to describe pigmented coelomocytes in sea cucumbers (by not considering respiratory pigments). This discovery, therefore, opens up a new paradigm on pigmented coelomocytes in echinoderms, which were previously thought to be only present in sea urchins and sea stars (Queiroz and Custódio, 2025; Smith et al., 2018; Wahltinez et al., 2023) and suspected in brittle stars (Wambreuse et al., 2024). Furthermore, it is interesting to note that, although sea cucumber carotenocytes and sea urchin red spherule cells contain different pigments, they appear to share certain functions, including in the antimicrobial response and the ability to scavenge ROS (Coates et al., 2018; **Fig. 14**). It could, therefore, be hypothesised that sea cucumbers and sea urchins have independently acquired similar functional cell types through similar immune selection pressure but by two different molecular systems. While further research is underway to determine the significance of the distribution of carotenocytes in sea cucumbers and confirm their function in immune regulation, carotenoid storage and the regeneration process, we have evidence that such cell types are broadly distributed in sea cucumbers and believe that they play a central role in the sea cucumber homeostasis.

## Material and method

### 1. Coelomocyte collection and morphological characterisation

#### 1.1. Collection and maintenance of organisms

Adult specimens (42) of *Holothuria forskali* Delle Chiaje, 1824 were obtained by the collection service of the Roscoff Biological Station, from where they were initially collected by scuba diving in Morlaix Bay (Brittany, France) just before their shipment. As soon as they were delivered to the University of Mons (Belgium), the specimens were kept in a closed-circuit tank containing 400 litres of filtered seawater, with a substrate made of small pebbles, at a temperature varying between 14° and 17°c throughout the year and salinity of between 33 and 35 psu. An artificial circadian rhythm was recreated using neon lighting set at a constant exposure time from 8 am to 8 pm. The specimens were fed once a week with a mix of dried algae in agar-agar-based gel. Before any experiments, the specimens were acclimatised for at least two weeks in the tanks.

#### 1.2. Coelomocyte Harvesting from the two body fluids

For each specimen, coelomocytes were collected from the two body fluids of interest: the perivisceral fluid (PF) from the general cavity and the hydrovascular fluid (HF) from the Polian vesicle. To perform the PF collection, a longitudinal incision was first made on the bivium, from the anterior to the posterior part of the animal using a scalpel to open the integument between two radial canals (to avoid any contamination with HF), allowing PF to be harvested in a 15 ml tube. The incision was then extended anteriorly to access the Polian vesicle, which was then placed over another 15 ml tube to pierce it and harvest the dropping HF. Body fluids were systematically placed on ice before subsequent analyses to avoid coelomocyte aggregation/death.

#### 1.3. Establishment of cell concentration and proportion

Typically, 20 µl of each body fluid was pipetted directly in collected HF and PF cand mixed at an equivalent volume with calcium- and magnesium-free artificial seawater containing EDTA (CMFSW + EDTA: 460 mM NaCl; 10.7 mM KCl; 7 mM Na_2_SO_4_; 2.4 mM NaHCO_3_; 20 mM HEPES; 70 mM EDTA; pH = 7.4) to avoid cell aggregation (Smith 2019). Cells were then counted using a Neubauer hemacytometer within an hour post-body fluid collection. To do this, 10 µl were placed on the hemacytometer, and the 16 squares corresponding to a total volume of 0.1 mm³ were photographed under the microscope (Axio imager A1, Zeiss). Cells were counted manually using ImageJ software V1.54f, and their concentrations per milliliter were calculated using the following formula:

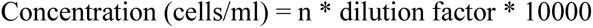

Once concentrations were obtained, the proportion could also be calculated following the formula:

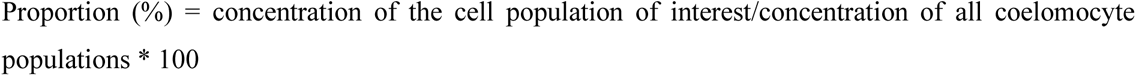

Note that if spermatozoa are present (see Caulier et al., 2024), they are not assimilated into a coelomocyte population (i.e., not counted). Cell populations were identified based on previous studies on holothuroid coelomocytes (Chia and Xing, 1996; Queiroz and Custódio, 2024). To highlight significant differences in the concentration and proportion of cell populations between the two body fluids, a Wilcoxon signed rank test was performed in R V.4.4.2 (α = 0.05).

#### 1.4. Microscopic analysis of coelomocytes and macroscopic pictures

Depending on the needs, cells were observed using different microscopy techniques, including light microscopy, fluorescence microscopy and scanning electron microscopy (SEM). Light microscopy and fluorescence microscopy were generally performed on fresh samples using an epifluorescence microscope (Axio imager A1, Zeiss) with or without an activated laser, depending on the type of microscopy desired. Three fluorescent filters could be used, yielding three different excitation wavelengths: 350 nm (corresponding to violet), 495 nm (corresponding to cyan), and 589 nm (corresponding to green). If necessary, CMFSW with 70 mM EDTA could be used when cell aggregates had to be avoided (especially used for cell counting and flow cytometry). The SEM protocol was taken from Wambreuse et al. (2025), which was initially adapted from Smith et al. (2019). Briefly, this protocol includes an incubation phase where cells can settle on the slide in a humid chamber for 30 minutes before being fixed with a glutaraldehyde solution, dehydrated with successive baths of ethanol, chemically dried with successive baths of hexamethyldisilane and coated with a thin layer of a mixture of palladium and gold (60:40%). Samples were then observed under a scanning electron microscope (JSM-7200F, Jeol). For the tissue samples, the same protocol was used, but the tissues were immersed directly in the fixation solution containing glutaraldehyde. Finally, some macroscopic pictures were taken to show certain parts of interest in the anatomy of the organisms; these were produced using a Leica M28 binocular camera or an Olympus TG-6 digital camera.

### 2. Monitoring of coelomocyte activity by time-lapse imaging

PF and HF were collected as described above and placed directly on ice. Twenty-five minutes later, 100 µl of the fluid to be analysed was deposited on a slide already mounted on the microscope and without a cover glass. After five minutes on the glass for the cells to settle, time-lapse images were captured at 30-second intervals for 30 minutes at 200x magnification (Axio imager A1 microscope, Zeiss). The image sequences obtained were then loaded into ImageJ software V1.54f to build the videos. To quantify cell aggregation, the pixel threshold was automatically modified to obtain a binary value for each pixel. After adjusting the resolution to the actual pixel size, ”particle analyses” were carried out; the first targeting particles with a surface area between 4 and 50 µm^2^, and the second targeting particles with a surface area greater than 4 µm^2^. The output of these analyses includes the number of particles on each image and the mean particle number. These two analyses were used to count isolated HELs and the mean particle size (including cells and aggregates) over time, respectively. Finally, a linear least squares regression was applied to these time series, and a Mann-Kendall test was run in R V4.4.2. to check for a relation over time (α = 0.05).

## 3. Immunostimulation using lipopolysaccharide injection

In order to study the response of coelomocytes from both fluids to immunological stress, lipopolysaccharide (LPS) injections were performed 24 hours prior to harvesting the body fluids. These injections consisted of 200 µL of sterile CMFSW without EDTA, containing 5 mg/ml of LPS from *Escherichia coli* O111:B4 (L2630; Merck). As a control, injections of CMFSW without LPS were used to avoid injection stress bias. In total, seven individuals received an LPS injection, and six received a control injection. These injections were carried out using 1 ml syringes with 23 g needles in the right anterior part of the animal. Immediately afterwards, the inoculated individuals were isolated in their tanks until the following day. Coelomocytes from both body fluids were collected as previously explained (see section 1.2). Importantly, during the dissections, the sex of the individuals was systematically noted to avoid misinterpretation of results due to an unequal sex distribution between the conditions (identified based on the gonadal aspect according to Tuwo and Conand, 1992). For each fluid sample, no more than 2 ml of fluid was used for the RNA extraction, and 20 µl was systematically retained for cell counting. The concentration and proportion of each coelomocyte population were calculated as above (see section 1.3). To reveal significant differences between control-injected and LPS-injected individuals in terms of concentration and proportion, a Mann-Whitney U test was performed in R V4.4.2 (α = 0.05).

### 4. Transcriptomics analysis using RNA-sequencing

#### 4.1. RNA extraction, library preparation and sequencing

For RNA extraction, freshly collected body fluids were centrifuged at 500 × g and 4°c for 5 minutes to pellet cells. The volume used for RNA extractions was noted and, combined with cell counts, allowed us to know exactly how many cells were in the pellet and the proportions of the different cell populations. RNA extractions were carried out using the Qiagen RNeasy mini kit according to the manufacturer’s instructions. Other samples were also prepared for RNA-sequencing to be used in other projects; these consisted of a stone canal and podia collected from other individuals, and their RNA extraction was performed using a Trizol kit according to the manufacturer’s instructions (Merck; T9424). The concentration and purity of extracted RNA were determined using a nanodrop spectrophotometer (Denovix DS11), and the RNA integrity value (RIN) was assessed using the Agilent 2100 bioanalyser (Agilent RNA 6000 Nano kit). Only the three samples per condition showing the best RNA quality of HF and PF were retained for RNA-seq (note that HF and PF extracts were selected from the same individuals to allow intra-individual comparisons). Preparation of the cDNA libraries and sequencing were carried out by the Beijing Genomics Institute (BGI, Hong Kong). Briefly, the cDNA libraries were assembled as follows: mRNAs were isolated from total RNA using the oligo(dT) method; purified mRNAs were fragmented and reverse transcribed into the first cDNA strand, prior to synthesis of the second cDNA strand; Double-stranded cDNA fragments were end-repaired, 3’-adenylated and ligated to sequencing adapters; cDNA fragments of appropriate size were selected and enriched by PCR; PCR products were heat-denatured and single-stranded DNA was cyclized by oligo splint and DNA ligase. The libraries were then sequenced on the BGISEQ-500 platform.

#### 4.2. Raw data filtering, de novo assembly and gene expression level

Before assembly, raw data were filtered to eliminate adapter-polluted reads, reads containing more than 5% unknown bases, and low-quality reads (i.e., reads comprising more than 40% of bases with a quality value below 20). As no reference genome exists for *H. forskali*, the transcriptome was assembled *de novo* using Trinity software (V2.5.1). The obtained transcripts were then grouped using Tgicl software (V2.5.1) to eliminate redundancies and obtain the final sequences referred to as unisgenes. The unigenes can either form clusters comprising several unigenes with more than 70 % overlapping or singletons (i.e., single unigenes). To obtain the expression level of each unigene (or cluster), reads were mapped onto the transcriptome using Bowtie2 (V.2.2.5) and the unigene expression level was calculated using RSEM (V.1.2.12). The result is expressed as "fragments per kilobase of the transcript, per million mapped reads" (FPKM). As the sequence length is a proxy of the assembly quality, the size distribution of unigenes was represented.

#### 4.3. Functional annotation of unigenes

To assess the completeness of the assembled transcriptomes, BUSCO annotation was performed for each sequenced library and the merged transcriptome using the tool BUSCO in the Galaxy server (https://usegalaxy.eu; V5.4.6) against the metazoan_odb10 dataset (954 BUSCOs). The BUSCO metric attempts to provide a quantitative assessment of the completeness of genomics data by classifying orthologs into the following four categories: complete and single-copy, complete and duplicated, fragmented, or missing BUSCOs (Simão et al., 2015). Moreover, to obtain an initial indication of unigene function, the sequence of each unigene was aligned with several protein databases, including NCBI NT, NCBI NR, GO – Gene Ontology, KOG – EuKaryotic Orthologous Groups, KEGG – Kyoto Encyclopedia of Genes and Genomes, SwissProt and InterPro using Blast (V2.2.23), Diamond (V0.8.31), Blast2GO (V2.5.0) and InterProScan5 (V5.11-51.0). The unigene annotation provides an E-value that quantifies the degree of similarity with the annotation: only annotations with an E value < 10^-5^ were taken into account.

#### 4.4. Differential expression analysis

Differential expression analyses were performed to answer two main questions: i) “What are the differentially expressed genes (DEGs) between control and LPS-injected individuals?” and ii) “What are the DEGs between HF and PF?” The first question was assessed for PFs (PF analysis; n = 3), HFs (HF analysis; n = 3) and all fluids together (merged fluid analysis; n = 6). The second question was assessed for control individuals (CON analysis; n = 3), LPS-injected individuals (LPS analysis; n = 3) and all individuals together regardless of the stress condition (merged condition analysis; n = 6). In addition, a differential expression analysis was performed between male and female samples to reveal a potential sex-specific expression (n = 4 in males and 8 in females). Differential expression analyses were performed using the DESeq2 package. The output of these analyses consists mainly of two metrics, a fold change value (FC) and a false discovery rate (FDR; corresponding to a Wald statistical test p-value adjusted following the Benjamini–Hochberg procedure). Only unigenes having a |log_2_(FC) value| ≥ 1 and an FDR ≤ 0.05 were considered as significantly differentially expressed.

Heat maps were carried out using the package “Pheatmap” to illustrate the result based on normalised read count. Principal component analyses (PCA) were also generated to illustrate the heterogeneity in gene expression between different samples (based on log-transformed FPKM values) in the MetaboAnalyst (V6.0) server (https://www.metaboanalyst.ca/). In addition, Venn diagrams were performed to highlight DEGs shared between the differential expression analyses. They were produced using the web tool “Venn” on the server https://bioinformatics.psb.ugent.be/webtools/Venn/.

#### 4.5. Functional enrichment analysis

Functional enrichment analyses were performed using KEGG annotation to reveal the most differentially expressed metabolic pathways. We chose to present, for each analysis, the twenty most enriched pathways and to group all these pathways in a table to compare the most enriched pathways between the different analyses (i.e., analyses comparing stress conditions and body fluids). For each pathway, various measures are provided: the pathway ranking (from 1 to 20), the q value represented by a colour scale (the lower the q value, the greater the differential expression of the pathway), the rich factor representing the percentage ratio between the number of annotated DEGs over the number of all annotated unigenes, the number of annotated DEGs in the pathways and the ratio between up- and down-regulated unigenes within these DEGs.

### 5. Pigment identification and quantification by High-Performance Liquid Chromatography and spectrophotometry

To obtain a first identification of the nature of the coelomocyte pigments, pigments were extracted using liquid-liquid phase extraction. This was conducted on lyophilised cell pellets from HF and PF of seven individuals of *H. forskali* to compare the distribution and amount of pigment between the two fluids. To estimate the number of cells in the pellet, the concentration of the fluid, as well as the volume, was determined prior to the centrifugation (see section 1.3). In addition to holothuroid body fluids, two samples of bovine blood (*Bos taurus*), acquired from the conventional meat industry, were included in the analysis to serve as a positive control for potential haemoglobin, following previous hypotheses that hemocyte pigmentation is due to haemoglobin (Baker and Terwilliger, 1993; Crescitelli, 1945). Body fluids were centrifuged at 500 g and 4°c for 5 minutes to isolate the cells from the supernatant. Rapidly after the centrifugation, the different samples and positive control were flash-frozen in liquid nitrogen before being freeze-dried under a vacuum at -60°C (using CHRIST Alpha 1-2 LDplus freeze-dryer) for 24 hours. Dried samples were stored and protected from light at 6°c until their processing. In order to determine the pigment affinity, several solvents were used to dissolve the lyophilised samples, including methanol, chloroform and distilled water (volume ratio of 2:3:2.7) and separate pigments in different phases. First, 2 ml of ice-cold methanol (HPLC grade) was added, followed by 700 µl of ice-cold distilled water, before vortexing the samples for 1 minute. Next, the cell membranes were disrupted using an ultrasonic homogeniser (U50 control, IKA Labortechnik) set to cycle 1 and 50% amplitude for 45 seconds. Once completed, 1.5 ml of ice-cold chloroform (HPLC grade) was added, and the samples were vortexed again for 1 minute before being agitated on ice at 300 rpm for 10 minutes. After adding 1.5 ml of ice-cold chloroform and 2 ml of ice-cold distilled water, the samples were centrifuged at 1500 x g for 30 minutes at 4°c (Amplitude 20x, cycle 1). The resulting polar (water + methanol) and apolar (chloroform) phases were separated: 2 ml of each phase was collected in separate tubes and then centrifuged at 10,000 x g for 5 minutes to ensure that all debris was removed. The supernatants were then collected, and the absorption spectrum, from 220 to 750 nm of excitation wavelength, was obtained for each sample using a nanodrop spectrophotometer (Denovix DS-11) with quartz cuvettes. Note that some samples of HF had to be diluted for the measurement to avoid the saturation of the device. After confirming that the red pigmentation could be assigned to carotenoids (see HPLC results; **Fig. 6**), a quantification of the mass of carotenoids per million cells in the apolar phase was performed using a pure canthaxanthin standard (canthaxanthin_trans; Merck n°11775) of known concentration to obtain a calibration curve. This was done by measuring the absorbance of the standard at increasing dilutions in chloroform (maximum absorbance at 490 nm in chloroform; **Fig. S3A**). This allows us to quantify the mass of the carotenoid for each sample. As the dry masses of the HF and PF pellets were insufficient, the abundance of pigments was normalised according to the number of cells in the pellet and expressed in µg per million cells. In order to highlight a putative significant difference between the two body fluids, a Wilcoxon signed-rank test was carried out in R V4.4.2 (n = 7; α = 0.05). Moreover, knowing the number of HELs in each sample, a Pearson correlation test was performed to see if the mass of carotenoids in each sample is correlated to the number of HELs (α = 0.05). This was done considering two body fluids (HF and PF; n = 14) and only considering HF samples (n = 7). The normality of data was determined using the Shapiro-Wilk normality test in R V4.4.2 (α = 0.05).

In order to obtain a better identification of carotenoid pigments, High-Performance Liquid Chromatography (HPLC) was performed, using samples of HF, PF, and Polian vesicles (containing large red coelomocyte aggregates as in **Fig. 7A-C**), which were collected from four individuals. Body fluid and Polian vesicle samples were prepared as explained in the spectrophotometry protocol. Once dried, the mass of each sample was measured using a high-precision balance (Sartorius Extend). For the extraction, samples were dissolved in a 1.5 ml extraction solution consisting of 95% methanol and 2% ammonium acetate with an additional 100 µl of trans-β-Apo-8’-carotenal (commercial standard, Roth ®) that was added to serve as an internal standard for mass quantification. Samples were then homogenised using beads at 30Hz for 10 minutes with the Mixer Mill MM 400 (Retsch) and filtered with 0.2 µm mesh filters. Pigments were finally identified using an Agilent 1260 Infinity HPLC system following the methodology developed by Brotas and Plante-Cuny (2003) and adapted in David et al. (2023). The relative abundance of pigments in each sample was determined by measuring the areas under the peaks (AUP) of the curve at a wavelength of 470 nm to target the carotenoid pigments (David et al., 2023). The AUP of different pigments were standardised with the AUP of the known mass of the commercial standard (trans-β-Apo-8’-carotenal). The abundance of pigments in Polian vesicles was normalised according to their mass, while the abundance of pigments in the body fluid samples was converted into µg per million cells. To identify the chemical nature of the pigments, the resulting HPLC peaks were compared with peaks of commercial carotenoid compounds such as astaxanthin, canthaxanthin, and all-trans-echinenone (DHI Laboratory Products) that were also passed through HPLC (**Fig. S5B**).

### 6. Analysis of coelomocyte autofluorescence using Spectral Flow Cytometry

In order to quantify the autofluorescence of cells, samples of PF and HF were analysed by spectral flow cytometry. HF and PF fluids were collected as previously described and directly mixed with an equivalent volume of CMFSW+EDTA 70mM to avoid clotting. Every step was carried out on ice to prevent cell deterioration. Samples were filtered with a cell strainer CellTrics equipped with a 100 µm mesh (Sysmex), and 10 µl was pipetted to establish the cell concentration. The samples were then centrifuged for 5 minutes and 4°c at 500 × g to resuspend the pellet in the flow cytometry buffer consisting of (3 × PBS buffer containing 20 mM HEPES and 50 mM EDTA; pH = 7.4; inspired by Smith et al. (2019) to obtain a final concentration of 10^7^ cells per ml. The Sony ID7000™ Spectral Cell Analyser, equipped with 5 lasers (355 nm, 405 nm, 488 nm, 561 nm, and 637 nm), was used for data acquisition. Gating strategy as well as individual spectra for each cell population are shown in **Fig. S7** and **Fig. S8**, respectively.

### 7. FACS Hemocyte-like cell pigment analysis

In order to isolate HELs from HF, HF and PF from one individual were sampled and prepared similarly as described for spectral flow cytometry (see section 6). The two fluids were then analysed using the BD FACS Aria^TM^ III cell sorter. The HEL population was easily identified by comparing the profile of the cells in the two body fluids. This population was sorted on the basis of autofluorescence criteria, previously determined using the Sony ID7000™ Spectral Cell Analyser. Once sorted, the purity of the sample was calculated by analysing the sample again, using the same parameters. Their presence and purity were also assessed visually under the microscope (Axio imager A1, Zeiss). Having established that the sample was sufficiently pure, cells were centrifuged at 500 × g and 4°c for 5 minutes and the pellet was stored at -80°c, protected from light, until the pigment analysis. For pigment extraction, the pellet was prepared as for spectrophotometry analysis in section 5 and resuspended in 1 ml of chloroform before a brief sonication to release the pigment. After a brief centrifugation at 10,000g and 4°c for 10 minutes to pellet cell debris, the sample was analysed by spectrophotometry (as in section 5). To estimate the amount of carotenoid per cell, the absorbance obtained was compared to a commercial canthaxanthin standard (canthaxanthin_trans; Merck; 11775) using a calibration curve (see **Fig. S5A**). Knowing the initial number of HELs in the sample (number of events counted by the flow cytometer), the final quantity of carotenoid was calculated in µg per million HELs.

### 8. Hemocyte-like cell gene and protein expression

#### 8.1. Relation between gene expression and HEL proportion

To test the relation between the proportion of HELs and the differential expression between PF and HF, a differential expression analysis was carried out between the HF and the PF for each individual. This analysis was conducted using the PossionDis algorithm, which is based the detection of DEGs on the fold change and Poisson distribution according to Audic and Claverie (1997). Only unigenes with a value of |log2(FC)| ≥ 1 and an FDR ≤ 0.05 were considered to be significantly differentially expressed. The proportion of DEGs obtained for each individual (i.e., the number of DEGs out of the total number of genes in the respective transcriptomes) was then correlated to the proportion of HELs in the HF, determined before RNA extraction by cell counting (see section 1.3). In addition to the DEGs, the fluid-specific expressed genes (FSEGs) proportion was tested for a putative relation with the proportion of HELs in HF. These genes had a null expression in one of the two fluids and were identified based on Venn diagram analyses. To statistically test the relations, Pearson correlation tests were performed (α = 0.05) after checking the normality of data using the Shapiro-Wilk normality test in R V.4.2.2 (α = 0.05).

#### 8.2. Proteomics analysis by mass spectrometry and validation of transcriptomics differential analysis

To validate the transcriptomics results and obtain the proteome of a highly enriched population of HELs, mass-spectrometry was carried out on the PF and the HF of one individual. Cells from the two body fluids were collected and counted as explained previously (see section 1.3). This individual in particular was selected because it had a very high concentration of HELs in its HF (> 97%). After centrifugating for 5 minutes at 500 × g and 4°c, cell pellets were isolated and flash-frozen. To perform the protein extraction, pellets were resuspended in a 5% acetic acid solution and 8M urea and incubated for 1 hour. Then, ultrasonication was performed three times for 10 sec (amplitude 20x, cycle 1) before centrifuging the samples at 18,000 × g for 15 minutes to eliminate cell debris. The protein concentration in the supernatant was assessed using a Bradford assay with Bovine Gamma Globulin as a standard, and concentrations were adjusted to 50µg/µl for downstream processing. To reduce the proteins, a solution containing dithioerythritol (DTE) was added to obtain a final concentration of 12.5 mM before incubating for 25 minutes at 56°c under agitation. The samples were then alkylated for 30 minutes in the dark by adding an iodoacetamide (IAA) solution at a final concentration of 25 mM. Proteins were precipitated by adding 4 volumes of ice-cold acetone, incubating for 4 hours at -20°c and centrifuging at 4°c for 20 minutes at 18,000 × g. After discarding the supernatant, the pellet was resuspended in a solution at 50 mM NH_4_HCO_3_ containing 2 µg/20 µl of trypsin (Promega) and incubated overnight at 37°c in order to lyse proteins into peptides. The lysis was stopped by adding 5µl of formic acid 0.5% to the 20µl of solution. Samples were centrifuged a last time for 15 minutes at 11,000 × g and 4°c. The peptide concentration was established using the Pierce^TM^ Quantitative Colourimetric Peptide Assay kit, and the supernatant was mixed with the loading buffer containing pepcalmix to obtain a final peptide concentration of 4µg/10µl before and 100 fmol/10µl before the injection.

Protein identification and quantification were performed following a label-free strategy on a UHPLC HRMS platform (Eksigent 2D ultra, AB SCIEX, TripleTOF™ 6600). Peptides (4µg) were separated on a 15 cm C18 column (YMC-Triart 12nm, S – 3 µm, 150 x 0.3 mm ID, 1/32”) using a linear acetonitrile (ACN) gradient [3-80% (v/v), in 75 min] in water containing 0.1% formic acid (v/v) at a flow rate of 5 µL min-1. Spectra were acquired in data-independent (DIA, SWATH) acquisition modes. For SWATH analysis, 100 incremental steps were defined as windows of variable m/z values over a 400/1250 m/z mass range. The MS/MS working time for each window was 7 ms, leading to a duty cycle of 2.65s per cycle. DIA analysis for identification and quantification was done using DIA-NN™ software (V1.9.2, for academic use) using the FASTA file format of the coelomocyte proteome of *H. forskali* and annotated with Uniprot standards. This proteome was acquired based on the coding sequence prediction using the TransDecoder software (V3.0.1) from the merged coelomocyte transcriptome. Proteins with a peptide count < 2 were filtered out due to their unreliable annotation.

To validate the transcriptomics differential expression analysis between HF and PF, the expression of proteins annotated with a DEG was compared to the expression of the corresponding DEGs. As one protein could be annotated by several genes, only proteins annotated with one gene were considered to ensure a real correspondence between the protein and the gene expression. A fold change value of PF versus HF intensity was calculated for these proteins to correlate it with the fold change value of transcriptomics analysis (under log_2_(fold change)). To test this correlation, a Pearson test was performed (α = 0.05) after checking the normality of the data using a Shapiro-Wilk test on R V4.2.2 (α = 0.05).

#### 8.3. Search for marker proteins for Hemocyte-like cells

To investigate further the function of HELs, a targeted analysis was carried out on candidate marker proteins. To do this, genes and proteins up-regulated in the HF (highly enriched in HELs) were extracted to conduct functional analysis. For transcriptomics analysis, genes having a fold change value > 2 were selected, whereas for proteomics analysis, proteins with a fold change value > 5 were selected, considering the need for a more stringent selection due to the lack of replicates. For this analysis, proteins annotated by several genes were considered, but only if the first three annotations were consistent with each other (i.e., the corresponding unigene having the same Nr annotation or coding for proteins of the same family, representing 98.7% of the identified proteins with a peptide count > 2). The coding peptide sequences were obtained from the transcriptome using the TransDecoder software (V3.0.1). The resulting proteins were aligned against the proteome of the species *Apostichopus japonicus* for protein-protein network analysis using STRING V12.0. The interactome significance was assessed using the PPI p-value in STRING. Moreover, the protein networks obtained by MS and RNA-seq analyses were compared by looking at the significantly enriched biological processes (from the Gene Ontology database) shared between the two analyses (FDR < 0.05). In addition, a string local network analysis was used to highlight functional protein clusters of interest (FDR < 0.05). In this case, the expression and annotation of proteins identified in the cluster were further analysed using a heat map visualisation carried out in Metaboanalyst V6.0 (log-normalised FPKM after t-test filtering). The annotation displayed corresponds to the gene Nr annotations, but the corresponding STRING annotation is shown in **Table S8**. Finally, to emphasise the most specific proteins in HELs, the fifteen proteins having the highest fold change value in the MS analysis were listed with their annotation and peptide count as an annotation proxy (i.e., the higher the peptide count is, the more reliable the gene annotation is).

As a complementary analysis, a search was carried out for the expression of globin, in line with the hypothesis that HEL pigmentation is due to haemoglobin (Baker and Terwilliger, 1993; Crescitelli, 1945). To do this, a local tBlastn was performed using Bioedit V7.7.1 on the whole transcriptome (E-value < 10^-5^) based on four existing peptide sequences coding for globin in echinoderms reported as intracellular in coelomocytes (Christensen et al., 2015; see **Fig. S10A**). Obtained unigenes were blasted again using Blastx and were only considered if the first hit matched the annotation of “globin”. The resulting unigenes were finally searched in the DEG list between PF and HF (i.e., |log_2_(FC) value| ≥ 1 and an FDR ≤ 0.05), and their expression was visualised using a heatmap in MetaboAnalyst V6.0 (based on FPKM log-normalised value). A search for proteins annotated with these genes was also carried out in the MS protein list to compare their expression between PF and HF samples.

### 9. Redox analysis

To investigate the antioxidant properties of HELs, we (i) looked at the expression of genes involved in reactive oxygen species (ROS) reduction and (ii) analysed ROS production among the different coelomocyte populations by flow cytometry and fluorescent microscopy using a ROS marker.

#### 9.1. Expression of antioxidant gene

To check if antioxidant genes could be overexpressed in HELs, we searched in the transcriptome genes known to have key functions in antioxidant response, including genes annotated as “superoxide dismutase” (SOD), “catalase” (CAT) and “glutathione peroxidase” (GPx) (Vulien et al., 2025; Billier and Takahashi, 2013). We first checked if some of these genes were significantly differentially expressed between PF and HF based on differential transcriptomics analysis (see section 4.4). Then, in order to study the expression of these antioxidant genes as a functional entity, the expression of all of these genes that were identified in the transcriptome was summed (sum of FPKM values) within each gene family (i.e., for SOD, CAT and GPx) as well as for their total (i.e., sum of FPKM value of the genes from the three families). These were compared between PF and HF samples using a Wilcoxon signed-rank test in R V4.2.2 (α = 0.05; n = 6). To further visualise the gene expression of the gene of interest (i.e., antioxidant genes), a heatmap was carried out based on FPKM values in the MetaboAnalyst (V6.0) server (log(FPKM)). Finally, a relation was tested between the ratio of PF expression out of HF expression as a function of HELs proportion in the HF for each individual to determine whether the proportion of HELs influences the antioxidant gene expression. This relation was tested using a Pearson correlation test (α = 0.05) after checking the normality of data using a Shapiro-Wilk test R V4.2.2 (α = 0.05).

#### 9.2. ROS production analysis

To estimate the ROS production in the different coelomocyte populations, 2′,7′-Dichlorodihydrofluorescein diacetate (DCFH-DA) was used as a marker of intracellular ROS (Merck D6883). DCFH-DA is a cell-permeable non-fluorescent probe that forms fluorescent DCF when oxidised by intracellular ROS, having an excitation wavelength of 485 nm and an emission range between 500 and 600 nm (Reiners et al. 2017). ROS production was compared in coelomocyte populations from the two body fluids exposed or not to lipopolysaccharides. Four individuals were used for this experiment. Their fluids were collected and mixed directly with CMFASW+EDTA 70 mM. Each fluid was then separated into two tubes: one for the LPS condition and one for the control condition. The cells were then counted, pelleted using centrifugation for 5 minutes at 500 × g and 4°c and resuspended in coelomocyte culture medium (CCM; 0.5 M NaCl; 5 mM MgCl_2_; 1 mM EGTA; 20 mM HEPES; pH 7.2) to obtain a concentration of 5 million cells per ml. The CCM of treated samples contained a final concentration of 20 µg per ml of lipopolysaccharides from *Escherichia coli* O111:B4 (Merck L2630). After incubation for one hour, each sample was separated into three tubes corresponding to the three labels: a sample without labelling, a sample with 1 µg/ml of propidium iodide (PI – a marker of cell death; Merck P4170) and a sample with 5 µM of DCFH-DA (Final concentration). The DCFH-DA was first added to the corresponding samples. Samples were incubated again for 30 minutes, and the cells were washed and resuspended in the flow cytometry buffer (3 × PBS buffer containing 20 mM HEPES and 50 mM EDTA; see section 6). PI was added to the samples 5 minutes before the analysis. The different samples corresponding to different fluids, conditions and labelling were analysed using the Sony ID7000™ Spectral Cell Analyser. We used the Weighted Least Squares Method (WLSM) algorithm, which is implemented in the ID7000 system. Each fluorochrome was visualised on a dot plot alongside the other fluorochromes to assess potential unmixing inaccuracies. Moreover, the Sony ID7000™ Spectral Cell Analyser provides a tool called Autofluorescence Finder (AF), which allows for the identification and characterisation of autofluorescence signatures within complex multicolour samples. Once autofluorescence spectra are identified, they are processed and mathematically separated as independent fluorescence parameters (AF). Data for analysis were gated on live cells, and manual adjustments were made when necessary, before loading the result on FlowJo V10.10.0 for cell analysis. Cell populations were gated following the same strategy as in section 6. For each population, ROS-positive cell and dead cell proportions were calculated based on discrimination with unlabelled samples, using the FITC and PI channels, respectively. Moreover, ROS production was calculated using the median fluorescence. To highlight significant differences between populations, a Friedman statistical test was performed in R V4.2.2. with the Dunn test as a post-hoc test (only considering LPS-treated samples; n = 4; α = 0.05). Moreover, differences between LPS and CON samples were tested using the Mann-Whitney U test in R V4.2.2 (n = 4 in the LPS condition and n = 3 in the control condition). Finally, to confirm the results obtained by flow cytometry, samples were observed under an epifluorescent microscope using a FITC filter (excitation wavelength = 495 nm; Zeiss Axio 1). The level of ROS positivity was qualitatively compared between the coelomocyte populations by comparing them using the same fluorescence intensity with the same parameters.

## Acknowledgements

First of all, the authors would like to thank the collection service of the Roscoff Biological Station for supplying the biological material necessary for this study. They are also grateful to the project collaborators who welcomed them into their institutes: Prof. Nadia Ameziane and Prof. Cédric Hubas of the Concarneau Marine Station for providing access to the Mass Spectrometry Technical Platform (PtSMB) and Prof. Sandra Ormenese of the Flow Cytometry Platform of the GIGA Institute. They would also like to thank everyone who occasionally helped them during the laboratory work or for analyses, with special thanks to Guillaume Corbisier and Emilie Duthoo for their availability and Leandro Smacchia for his precious bioinformatic advice. They are also grateful to Prof. Patrick Flammang, Prof. Annie Mercier, Dr Jean-François Hamel and Dr Vanessa Tagliatti for the interesting discussions on the results of this study and their constant support. Finally, they would like to extend their warmest thanks to the laboratory technicians, Nathan Puozzo and Antoine Batigny, without whom this research would not have been possible.

## Author contributions

**Noé Wambreuse**: Conceptualisation, Methodology, Validation, Formal analysis, Investigation, Data curation, Writing – Original Draft, Visualisation, Supervision; **Estelle Bossiroy**: Conceptualisation, Methodology, Validation, Formal analysis, Investigation, Data curation, Writing – Original Draft, Visualisation; **Frank David**: Conceptualisation, Methodology, Investigation, Resources, Writing – Review & Editing; **Celine Vanwinge**: Methodology, Formal analysis, Investigation, Resources, Writing – Review & Editing, Validation; **Igor Eeckhaut**: Conceptualisation, Writing – Review & Editing, Supervision, Project administration, Funding acquisition; **Laurence Fievez**: Writing – Review & Editing, Supervision; Writing – Review & Editing; **Fabrice Bureau**: Funding acquisition, Project administration; **Sylvain Gabriele**: Resources, Writing – Review & Editing; **Tania Karasiewicz**: Methodology, Investigation, Validation, Writing – Review & Editing; Software; **Cyril Mascolo**: Methodology, Investigation, Validation, Writing – Review & Editing; **Ruddy Wattiez**: Resources, Writing – Review & Editing; **Guillaume Caulier**: Conceptualisation, Methodology, Investigation, Writing – Review & Editing, Supervision, Project administration, Funding acquisition; **Jérôme Delroisse**: Conceptualisation, Methodology, Software, Data curation, Investigation, Writing – Review & Editing, Supervision, Project administration, Funding acquisition.

## Supplementary files

Supplementary materials can be found in dedicated files or via the following figshare link: https://doi.org/10.6084/m9.figshare.c.7839038

## Videos

- **Video S1.** Time-lapse imaging of a spherule cell performing cell lysis upon contact with an early aggregate.
- **Video S2.** Stage I early aggregate formation
- **Video S3.** Stage II early aggregate formation
- **Video S4.** Stage III early aggregate formation
- **Video S5.** Formation of early aggregates in the hydrovascular fluid with the participation of hemocyte-like cells

## Tables

- **Table S1.** Statistical analysis – cell concentration and proportion difference between PF and HF.
- **Table S2.** Statistical analysis – cell proportion and concentration after a lipopolysaccharide immunostimulation.
- **Table S3.** Metrics of the *de novo* RNA-sequencing in *Holothuria forskali*.
- **Table S4.**List of differentially expressed genes (DEGs) between control and lipopolysaccharide-injected individuals
- **Table S5.** KEGG pathway enrichment analysis of the differential expression analysis between control (CON) and LPS-injected (LPS) individuals.
- **Table. S6.** List of differentially expressed genes (DEGs) between perivisceral fluid (PF) and hydrovascular fluid (HF).
- **Table S7.** KEGG pathway enrichment analysis of the differential expression analysis between perivisceral fluid (PF) and hydrovascular fluid (HF).
- **Table S8.** Proteins identified by mass spectrometry (MS) and validation of the RNA-sequencing differential expression analysis between the two fluids.
- **Table S9.** Results of the STRING Mapping on the proteome of *Apostichopus japonicus*.

## Figures

- **Fig. S1.** Spherule cell-specific behaviours and mobility.
- **Fig. S2.** Automated cell analysis using the Particle Analysis tool in ImageJ software.
- **Fig. S3.** Transcriptome assembly and general annotation in *Holothuria forskali*.
- **Fig. S4.** Differential expression analysis of coelomocytes between males and females in *Holothuria forskali*.
- **Fig. S5.** Identification of pigment based on commercial standards.
- **Fig. S6.** Linear relationship between astaxanthin and canthaxanthin concentrations.
- **Fig. S7.** Spectral flow cytometry gating strategy.
- **Fig. S8.** Autofluorescence spectra of coelomocytes revealed by spectral flow cytometry.
- **Fig. S9.** String protein networks based on genes overexpressed in the perivisceral fluid (PF).
- **Fig. S10.** Globin expression in coelomocytes of perivisceral and hydrovascular fluids.
- **Fig. S11.** Validation of DCFH-DA labelling for reactive oxygen species (ROS) detection.
- **Fig. S12.** Flow cytometry gating strategy of ROS production analysis.
- **Fig. S13.** Estimation of cell mortality using propidium iodide labelling and flow cytometry.

## Data availability

The SRA files from transcriptomics analysis were submitted to NCBI and are available under the BioProject ID PRJNA1259574. TSA files (including the merged transcriptome and translated proteome) and gene annotation, expression files are publicly available on the figshare repository following this link: https://doi.org/10.6084/m9.figshare.c.7839038. Additional raw and partially processed data regarding the results of this study will be provided under reasonable request.

## References

Audic S, Claverie J-M. 1997. The Significance of Digital Gene Expression Profiles. Genome Res 7:986– 995. doi:10.1101/gr.7.10.986

Babin A, Saciat C, Teixeira M, Troussard J-P, Motreuil S, Moreau J, Moret Y. 2015. Limiting immunopathology: Interaction between carotenoids and enzymatic antioxidant defences. Developmental & Comparative Immunology 49:278–281. doi:10.1016/j.dci.2014.12.007

Baker SM, Terwilliger NB. 1993. Haemoglobin Structure and Function in the Rat-Tailed Sea Cucumber, *Paracaudina chilensis*. The Biological Bulletin 185:115–122. doi:10.2307/1542135

Bandaranayake WM, Rocher AD. 1999. Role of secondary metabolites and pigments in the epidermal tissues, ripe ovaries, viscera, gut contents and diet of the sea cucumber *Holothuria atra*. Marine Biology 133:163–169. doi:10.1007/s002270050455

Biller JD, Takahashi LS. 2018. Oxidative stress and fish immune system: phagocytosis and leukocyte respiratory burst activity. An Acad Bras Ciênc 90:3403–3414. doi:10.1590/0001-3765201820170730

Boolootian RA, Giese AC. 1958. Coelomic corpuscles of echinoderms. The Biological Bulletin 115:53–63. doi:10.2307/1539092

Brasseur L, Caulier G, Flammang P, Gerbaux P, Eeckhaut I. 2018. Mapping of Spinochromes in the Body of Three Tropical Shallow Water Sea Urchins. Natural Product Communications 13:1934578X1801301222. doi:10.1177/1934578X1801301222

Brotas V, Plante-Cuny M-R. 2003. The use of HPLC pigment analysis to study microphytobenthos communities. Acta Oecologica 24:S109–S115. doi:10.1016/S1146-609X(03)00013-4

Brotosudarmo THP, Limantara L, Setiyono E, Heriyanto. 2020. Structures of Astaxanthin and Their Consequences for Therapeutic Application. International Journal of Food Science 1–16. doi:10.1155/2020/2156582

Caulier G, Hamel J-F, Mercier A. 2020. From Coelomocytes to Colored Aggregates: Cellular Components and Processes Involved in the Immune Response of the Holothuroid *Cucumaria frondosa*. The Biological Bulletin 239:95–114. doi:10.1086/710355

Caulier G, Jobson S, Wambreuse N, Borrello L, Delroisse J, Eeckhaut I, Mercier A, Hamel J-F. 2024. Chapter 24 - Vibratile cells and hemocytes in sea cucumbers—Clarifications and new paradigms In: Mercier A, Hamel J-F, Suhrbier AD, Pearce CM, editors. The World of Sea Cucumbers. Academic Press. pp. 403–412. doi:10.1016/B978-0-323-95377-1.00024-2

Chen J, Wei D, Pohnert G. 2017. Rapid Estimation of Astaxanthin and the Carotenoid-to-Chlorophyll Ratio in the Green Microalga *Chromochloris zofingiensis* Using Flow Cytometry. Marine Drugs 15:231. doi:10.3390/md15070231

Chen K, Zhang S, Shao Y, Guo M, Zhang W, Li C. 2021. A unique NLRC4 receptor from echinoderms mediates Vibrio phagocytosis via rearrangement of the cytoskeleton and polymerization of F-actin. PLoS Pathog 17:e1010145. doi:10.1371/journal.ppat.1010145

Chia FS, Xing J. 1996. Echinoderm coelomocytes. Zoological studies 35:231–254.

Chiaramonte M, Russo R. 2015. The echinoderm innate humoral immune response. Italian Journal of Zoology 82:300–308. doi:10.1080/11250003.2015.1061615

Christensen AB, Herman JL, Elphick MR, Kober KM, Janies D, Linchangco G, Semmens DC, Bailly X, Vinogradov SN, Hoogewijs D. 2015. Phylogeny of Echinoderm Haemoglobins. PLoS ONE 10:e0129668. doi:10.1371/journal.pone.0129668

Clements CS, Pratte ZA, Stewart FJ, Hay ME. 2024. Removal of detritivore sea cucumbers from reefs increases coral disease. Nat Commun 15:1338. doi:10.1038/s41467-024-45730-0

Coates CJ, McCulloch C, Betts J, Whalley T. 2018. Echinochrome A Release by Red Spherule Cells Is an Iron-Withholding Strategy of Sea Urchin Innate Immunity. J Innate Immun 10:119–130. doi:10.1159/000484722

Crescitelli F. 1945. A note on the absorption spectra of the blood of *Eudistylia gigantea* and of the pigment in the red corpuscles of *Cucumaria miniata* and *Molpadia intermedia*. The Biological Bulletin 88:30–36. doi:10.2307/1538168

Czeczuga B. 1975. Carotenoids in fish IV. Salmonidae and Thymallidae from Polish waters. Hydrobiologia 46:223–239. doi:10.1007/BF00043142

Czeczuga-Semeniuk, Czeczuga. 1999. Comparative studies of carotenoids in four species of crayfish. Crustac 72:693–700. doi:10.1163/156854099503735

David F, Herault G, Ameziane N, Meziane T, Badou A, Hubas C. 2023. Sex-specific seasonal variations in the fatty acid and carotenoid composition of sea cucumber gonads and implications for aquaculture. Mar Biol 170:47. doi:10.1007/s00227-023-04198-0

David F, Hubas C, Laguerre H, Badou A, Herault G, Bordelet T, Ameziane N. 2020. Food sources, digestive efficiency and resource allocation in the sea cucumber *Holothuria forskali* (Echinodermata: Holothuroidea): Insights from pigments and fatty acids. Aquacult Nutr 26:1568–1583. doi:10.1111/anu.13103

David F, Raymond G, Grys J, Ameziane N, Sadoul B. 2024. Survival, Growth, and Food Resources of Juvenile Sea Cucumbers *Holothuria forskali* (Echinodermata, Holothuroidea) in Co-Culture with Shellfish in Brittany (France). Aquaculture Nutrition :7098440. doi:10.1155/2024/7098440

Deridoux A, Heydari S, Gorb SN, Kanso EA, Flammang P, Gabriele S. 2025. Tube feet dynamics drive adaptation in sea star locomotion. doi:10.1101/2025.04.22.649740

Dhinaut J, Balourdet A, Teixeira M, Chogne M, Moret Y. 2017. A dietary carotenoid reduces immunopathology and enhances longevity through an immune depressive effect in an insect model. Sci Rep 7:12429. doi:10.1038/s41598-017-12769-7

Dong Y, Sun H, Zhou Z, Yang A, Chen Z, Guan X, Gao S, Wang B, Jiang B, Jiang J. 2014. Expression Analysis of Immune Related Genes Identified from the Coelomocytes of Sea Cucumber (*Apostichopus japonicus*) in Response to LPS Challenge. IJMS 15:19472–19486. doi:10.3390/ijms151119472

Dybas L, Fankboner PV. 1986. Holothurian survival strategies: Mechanisms for the maintenance of a bacteriostatic environment in the coelomic cavity of the sea cucumber, *Parastichopus californicus*. Developmental & Comparative Immunology 10:311–330. doi:10.1016/0145-305X(86)90022-4

Fingerman M. 1965. Chromatophores. Physiological Reviews. doi:10.1152/physrev.1965.45.2.296

Flammang P, Michel A, Van Cauwenberge A, Alexandre H, Jangoux M. 1998. A Study of the Temporary Adhesion of the Podia in the Sea Star Asterias Rubens (Echinodermata, Asteroidea) through their Footprints. Journal of Experimental Biology 201:2383–2395. doi:10.1242/jeb.201.16.2383

Fontaine AR, Hall BD. 1981. The haemocyte of the holothurian *Eupentacta quinquesemita* : ultrastructure and maturation. Can J Zool 59:1884–1891. doi:10.1139/z81-256

Fontaine AR, Lambert P. 1973. The fine structure of the haemocyte of the holothurian, *Cucumaria miniata* (Brandt). Can J Zool 51:323–332. doi:10.1139/z73-046

Galasso C, Corinaldesi C, Sansone C. 2017. Carotenoids from Marine Organisms: Biological Functions and Industrial Applications. Antioxidants 6:96. doi:10.3390/antiox6040096

Gostyukhina OL, Soldatov AA, Golovina IV, Borodina AV. 2013. Content of carotenoids and the state of tissue antioxidant enzymatic complex in bivalve mollusc *Anadara inaequivalvis* Br. J Evol Biochem Phys 49:309–315. doi:10.1134/S0022093013030055

Gross PS, Al-Sharif WZ, Clow LA, Smith LC. 1999. Echinoderm immunity and the evolution of the complement system. Developmental & Comparative Immunology 23:429–442. doi:10.1016/S0145-305X(99)00022-1

Hamel J-F, Eeckhaut I, Conand C, Sun J, Caulier G, Mercier A. 2022. Global knowledge on the commercial sea cucumber *Holothuria scabra*. Advances in Marine Biology. Elsevier. pp. 1–286. doi:10.1016/bs.amb.2022.04.001

Hamel J-F, Jobson S, Caulier G, Mercier A. 2021. Evidence of anticipatory immune and hormonal responses to predation risk in an echinoderm. Sci Rep 11:10691. doi:10.1038/s41598-021-89805-0

Han L, Hao P, Xiao H, Li W, Fan Y, Tian W, Tian Y, Wang L, Chang Y, Ding J. 2025. Molecular Mechanism of Body Color Change in the Ecological Seedling Breeding Model of *Apostichopus Japonicus*. doi:10.2139/ssrn.5082025

Han Q, Keesing JK, Liu D. 2016. A Review of Sea Cucumber Aquaculture, Ranching, and Stock Enhancement in China. Reviews in Fisheries Science & Aquaculture 24:326–341. doi:10.1080/23308249.2016.1193472

Hetzel HR. 1963. Studies on holothurian coelomocytes. I. A survey of coelomocyte types. The Biological Bulletin 125:289–301. doi:10.2307/1539404

Hibino T, Loza-Coll M, Messier C, Majeske AJ, Cohen AH, Terwilliger DP, Buckley KM, Brockton V, Nair SV, Berney K, Fugmann SD, Anderson MK, Pancer Z, Cameron RA, Smith LC, Rast JP. 2006. The immune gene repertoire encoded in the purple sea urchin genome. Developmental Biology 300:349–365. doi:10.1016/j.ydbio.2006.08.065

Hira J, Wolfson D, Andersen AJC, Haug T, Stensvåg K. 2020. Autofluorescence mediated red spherulocyte sorting provides insights into the source of spinochromes in sea urchins. Sci Rep 10:1149. doi:10.1038/s41598-019-57387-7

Hossain A, Dave D, Shahidi F. 2022. Antioxidant Potential of Sea Cucumbers and Their Beneficial Effects on Human Health. Marine Drugs 20:521. doi:10.3390/md20080521

Ince LM, Weber J, Scheiermann C. 2019. Control of Leukocyte Trafficking by Stress-Associated Hormones. Front Immunol 9:3143. doi:10.3389/fimmu.2018.03143

Janakiram N, Mohammed A, Rao C. 2015. Sea Cucumbers Metabolites as Potent Anti-Cancer Agents. Marine Drugs 13:2909–2923. doi:10.3390/md13052909

Jans D, Dubois P, Jangoux M. 1995. Defensive mechanisms of holothuroids (Echinodermata): Formation, role, and fate of intracoelomic brown bodies in the sea cucumber *Holothuria tubulosa*. Cell and Tissue Research 283:99–106. doi:10.1007/s004410050517

Jeyachandran S, Kiyun P, Ihn-Sil K, Baskaralingam V. 2020. Identification and characterization of bioactive pigment carotenoids from shrimps and their biofilm inhibition. J Food Process Preserv 44. doi:10.1111/jfpp.14728

Jobson S, Hamel J-F, Hughes T, Mercier A. 2021. Cellular, Hormonal, and Behavioral Responses of the Holothuroid Cucumaria frondosa to Environmental Stressors. Front Mar Sci 8:695753. doi:10.3389/fmars.2021.695753

Jobson S, Hamel J-F, Mercier A. 2022. Rainbow bodies: Revisiting the diversity of coelomocyte aggregates and their synthesis in echinoderms. Fish & Shellfish Immunology 122:352–365. doi:10.1016/j.fsi.2022.02.009

Karpiński TM, Ożarowski M, Alam R, Łochyńska M, Stasiewicz M. 2021. What Do We Know about Antimicrobial Activity of Astaxanthin and Fucoxanthin? Marine Drugs 20:36. doi:10.3390/md20010036

Kindred JE. 1924. The cellular elements in the perivisceral fluid of echinoderms. The Biological Bulletin 46:228–251. doi:10.2307/1536725

La Paglia L, Mauro M, Arizza V, Urso A, Luparello C, Vazzana M, Vizzini A. 2025. Bioinformatics analyses of the proteome of *Holothuria tubulosa* coelomic fluid and the first evidence of primary cilium in coelomocyte cells. Frontiers in Immunology 16. doi:10.3389/fimmu.2025.1539751

Lacouth P, Majer A, Arizza V, Vazzana M, Mauro M, Custódio MR, Queiroz V. 2024. Physiological responses of *Holothuria grisea* during a wound healing event: An integrated approach combining tissue, cellular and humoral evidence. Comparative Biochemistry and Physiology Part A: Molecular & Integrative Physiology 296:111695. doi:10.1016/j.cbpa.2024.111695

Lamare MD, Hoffman J. 2004. Natural variation of carotenoids in the eggs and gonads of the echinoid genus, *Strongylocentrotus*: implications for their role in ultraviolet radiation photoprotection. Journal of Experimental Marine Biology and Ecology 312:215–233. doi:10.1016/j.jembe.2004.02.016

Li Q, Qi R, Wang Y, Ye S, Qiao G, Li H. 2013. Comparison of cells free in coelomic and water-vascular system of sea cucumber, *Apostichopus japonicus*. Fish & Shellfish Immunology 35:1654–1657. doi:10.1016/j.fsi.2013.07.020

Li Q, Ren Y, Liang C, Qiao G, Wang Y, Ye S, Li R. 2018. Regeneration of coelomocytes after evisceration in the sea cucumber, *Apostichopus japonicus*. Fish & Shellfish Immunology 76:266–271. doi:10.1016/j.fsi.2018.03.013

Li Q, Ren Y, Luan L, Zhang J, Qiao G, Wang Y, Ye S, Li R. 2019. Localization and characterization of hematopoietic tissues in adult sea cucumber, *Apostichopus japonicus*. Fish & Shellfish Immunology 84:1–7. doi:10.1016/j.fsi.2018.09.058

Lidén M, Eriksson U. 2006. Understanding Retinol Metabolism: Structure and Function of Retinol Dehydrogenases. Journal of Biological Chemistry 281:13001–13004. doi:10.1074/jbc.R500027200

Liu B, Xing L, Liu S, Sun L, Su F, Cui W, Jiang C. 2024. Transcriptome analysis of purple and green *Apostichopus japonicus* reared under different breeding environments. Front Mar Sci 11. doi:10.3389/fmars.2024.1334761

Loo Y-M, Gale M. 2011. Immune Signalling by RIG-I-like Receptors. Immunity 34:680–692. doi:10.1016/j.immuni.2011.05.003

Lourtie A, Mussoi L, Caulier G, Isorez M, Mahavory H, Tolodraza T, Engels G, David F, Eeckhaut I, Mallefet J. 2024. Exploring the mimetic pigmentation of symbiotic shrimps associated with echinoderms. Symbiosis 94:107–127. doi:10.1007/s13199-024-01018-x

MacDonald CLE, Stead SM, Slater MJ. 2013. Consumption and remediation of European Seabass (*Dicentrarchus labrax*) waste by the sea cucumber *Holothuria forskali*. Aquacult Int 21:1279– 1290. doi:10.1007/s10499-013-9629-6

Majeske AJ, Bayne CJ, Smith LC. 2013. Aggregation of Sea Urchin Phagocytes Is Augmented In Vitro by Lipopolysaccharide. PLoS ONE 8:e61419. doi:10.1371/journal.pone.0061419

Manwell C, Baker CMA. 1963. A sibling species of sea cucumber discovered by starch gel electrophoresis. Comparative Biochemistry and Physiology 10:39–53. doi:10.1016/0010-406X(63)90101-4

Maoka T. 2011. Carotenoids in Marine Animals. Marine Drugs 9:278–293. doi:10.3390/md9020278

Matsuno T, Tsushima M. 1995. Comparative biochemical studies of carotenoids in sea cucumbers. Comparative Biochemistry and Physiology Part B: Biochemistry and Molecular Biology 111:597–605. doi:10.1016/0305-0491(95)00028-7

Meager A. 1999. Cytokine regulation of cellular adhesion molecule expression in inflammation. Cytokine & Growth Factor Reviews 10:27–39. doi:10.1016/S1359-6101(98)00024-0

Medina-Feliciano JG, Valentín-Tirado G, Luna-Martínez K, Beltran-Rivera A, Miranda-Negrón Y, García-Arrarás JE. 2025. Single-cell RNA sequencing of the holothurian regenerating intestine reveals the pluripotency of the coelomic epithelium. doi:10.7554/eLife.100796.2

Miki W. 1991. Biological functions and activities of animal carotenoids. Pure and Applied Chemistry 63:141–146. doi:10.1351/pac199163010141

Morrison RA, Hamel J-F, Sun J, Mercier A. 2024. Comparative analysis of phenotypes in the sea cucumber *Cucumaria frondosa* from the Arctic and the NW Atlantic. Arctic Science 10:69–86. doi:10.1139/as-2023-0025

Mundy NI, Stapley J, Bennison C, Tucker R, Twyman H, Kim K-W, Burke T, Birkhead TR, Andersson S, Slate J. 2016. Red Carotenoid Coloration in the Zebra Finch Is Controlled by a Cytochrome P450 Gene Cluster. Current Biology 26:1435–1440. doi:10.1016/j.cub.2016.04.047

Nair SV, Del Valle H, Gross PS, Terwilliger DP, Smith LC. 2005. Macroarray analysis of coelomocyte gene expression in response to LPS in the sea urchin. Identification of unexpected immune diversity in an invertebrate. Physiological Genomics 22:33–47. doi:10.1152/physiolgenomics.00052.2005

Orosa M. 2005. Analysis and enhancement of astaxanthin accumulation in *Haematococcus pluvialis*. Bioresource Technology 96:373–378. doi:10.1016/j.biortech.2004.04.006

Ota S, Morita A, Ohnuki S, Hirata A, Sekida S, Okuda K, Ohya Y, Kawano S. 2018. Carotenoid dynamics and lipid droplet containing astaxanthin in response to light in the green alga *Haematococcus pluvialis*. Sci Rep 8:5617. doi:10.1038/s41598-018-23854-w

Paluch EK, Aspalter IM, Sixt M. 2016. Focal Adhesion–Independent Cell Migration. Annu Rev Cell Dev Biol 32:469–490. doi:10.1146/annurev-cellbio-111315-125341

Pangestuti R, Arifin Z. 2018. Medicinal and health benefit effects of functional sea cucumbers. Journal of Traditional and Complementary Medicine 8:341–351. doi:10.1016/j.jtcme.2017.06.007

Patel SR. 2005. The biogenesis of platelets from megakaryocyte proplatelets. Journal of Clinical Investigation 115:3348–3354. doi:10.1172/JCI26891

Pérez-Portela R, Leiva C. 2022. Sex-Specific Transcriptomics Differences in the Immune Cells of a Key Atlantic-Mediterranean Sea Urchin. Front Mar Sci 9:908387. doi:10.3389/fmars.2022.908387

Perillo M, Sepe RM, Paganos P, Toscano A, Annunziata R. 2024. Sea cucumbers: an emerging system in evo-devo. EvoDevo 15:3. doi:10.1186/s13227-023-00220-0

Potts WTW. 2003. The Physiological Function of the Coelom in Starfish Larvae and Its Evolutionary Implications. Physiological and Biochemical Zoology 76:771–775. doi:10.1086/381463

Purcell S, Conand C, Uthicke S, Byrne M. 2016. Ecological Roles of Exploited Sea Cucumbers. Oceanography and Marine Biology: An Annual Review.

Purcell S, Lovatelli A, Gonzalez-Wanguemert M, Solis-Marin F, Samyn Y, Conand C. 2023. Commercially important sea cucumbers of the world.

Queiroz V, Custódio MR. 2025. Diversity of coelomocytes in the class Echinoidea (Echinodermata): A comparative study based on morphological evidence. Comparative Immunology Reports 8:200212. doi:10.1016/j.cirep.2025.200212

Queiroz V, Custódio MR. 2024. Diversity of coelomocytes in the class Holothuroidea. The World of Sea Cucumbers. Elsevier. pp. 377–401. doi:10.1016/B978-0-323-95377-1.00011-4

Ramírez-Gómez F, Aponte-Rivera F, Méndez-Castaner L, García-Arrarás JE. 2010. Changes in holothurian coelomocyte populations following immune stimulation with different molecular patterns. Fish & Shellfish Immunology 29:175–185. doi:10.1016/j.fsi.2010.03.013

Roberts MS. 1980. A comparative survey of some holothurian haemoglobins (PhD Thesis). University of Oregon theses, Dept. of Biology, MS, 1980.

Roberts MS, Terwilliger RC, Terwilliger NB. 1984. Comparison of sea cucumber haemoglobin structures. Comparative Biochemistry and Physiology Part B: Comparative Biochemistry 77:237–243. doi:10.1016/0305-0491(84)90250-5

San Miguel-Ruiz JE, García-Arrarás JE. 2007. Common cellular events occur during wound healing and organ regeneration in the sea cucumber *Holothuria glaberrima*. BMC Dev Biol 7:115. doi:10.1186/1471-213X-7-115

Santos R, Dias S, Pinteus S, Silva J, Alves C, Tecelão C, Pedrosa R, Pombo A. 2016. Sea cucumber *Holothuria forskali*, a new resource for aquaculture? Reproductive biology and nutraceutical approach. Aquac Res 47:2307–2323. doi:10.1111/are.12683

Schaefer L. 2014. Complexity of Danger: The Diverse Nature of Damage-associated Molecular Patterns. Journal of Biological Chemistry 289:35237–35245. doi:10.1074/jbc.R114.619304

Schillaci D, Cusimano M, Cunsolo V, Saletti R, Russo D, Vazzana M, Vitale M, Arizza V. 2013. Immune mediators of sea-cucumber *Holothuria tubulosa* (Echinodermata) as source of novel antimicrobial and anti-staphylococcal biofilm agents. AMB Express 3:35. doi:10.1186/2191-0855-3-35

Shahidi F, Synowiecki J. 1991. Isolation and characterization of nutrients and value-added products from snow crab (Chionoecetes opilio) and shrimp (*Pandalus borealis*) processing discards. J Agric Food Chem 39:1527–1532. doi:10.1021/jf00008a032

Simão FA, Waterhouse RM, Ioannidis P, Kriventseva EV, Zdobnov EM. 2015. BUSCO: assessing genome assembly and annotation completeness with single-copy orthologs. Bioinformatics 31:3210–3212. doi:10.1093/bioinformatics/btv351

Smith LC, Arizza V, Barela Hudgell MA, Barone G, Bodnar AG, Buckley KM, Cunsolo V, Dheilly NM, Franchi N, Fugmann SD, Furukawa R, Garcia-Arraras J, Henson JH, Hibino T, Irons ZH, Li C, Lun CM, Majeske AJ, Oren M, Pagliara P, Pinsino A, Raftos DA, Rast JP, Samasa B, Schillaci D, Schrankel CS, Stabili L, Stensväg K, Sutton E. 2018. Echinodermata: The Complex Immune System in Echinoderms In: Cooper EL, editor. Advances in Comparative Immunology. Cham: Springer International Publishing. pp. 409–501. doi:10.1007/978-3-319-76768-0_13

Smith LC, Hawley TS, Henson JH, Majeske AJ, Oren M, Rosental B. 2019. Methods for collection, handling, and analysis of sea urchin coelomocytes. Methods in Cell Biology. Elsevier. pp. 357–389. doi:10.1016/bs.mcb.2018.11.009

Smith VJ. 1981. The Echinoderms Invertebrate Blood Cells. Academic Press. pp. 513–562.

Stra A, Almarwaey LO, Alagoz Y, Moreno JC, Al-Babili S. 2023. Carotenoid metabolism: New insights and synthetic approaches. Front Plant Sci 13:1072061. doi:10.3389/fpls.2022.1072061

Strathmann RR. 1975. Limitations on Diversity of Forms: Branching of Ambulacral Systems of Echinoderms. The American Naturalist 109:177–190. doi:10.1086/282985

Stegeman JJ. 2000. Cytochrome P450 gene diversity and function in marine animals: past, present, and future. Marine Environmental Research 50:61–62. doi:10.1016/S0141-1136(00)00136-7

Sun H, Zhou Z, Dong Y, Yang A, Jiang B, Gao S, Chen Z, Guan X, Wang B, Wang X. 2013. Identification and expression analysis of two Toll-like receptor genes from sea cucumber (*Apostichopus japonicus*). Fish & Shellfish Immunology 34:147–158. doi:10.1016/j.fsi.2012.10.014

Taguchi M, Tsutsui S, Nakamura O. 2016. Differential count and time-course analysis of the cellular composition of coelomocyte aggregate of the Japanese sea cucumber *Apostichopus japonicus*. Fish & Shellfish Immunology 58:203–209. doi:10.1016/j.fsi.2016.06.060

Tan K, Zhang H, Lim L-S, Ma H, Li S, Zheng H. 2020. Roles of Carotenoids in Invertebrate Immunology. Front Immunol 10:3041. doi:10.3389/fimmu.2019.03041

Tan K, Zhang H, Zheng H. 2024. Carotenoid content and composition: A special focus on commercially important fish and shellfish. Critical Reviews in Food Science and Nutrition 64:544–561. doi:10.1080/10408398.2022.2106937

Tsushima M, Fujiwara Y, Matsuno T. 1996. Novel marine di-Z-carotenoids: cucumariaxanthins A, B, and C from the sea cucumber *Cucumaria japonica*. J Nat Prod 59:30–34. doi:10.1021/np960022s

Tuwo A, Conand C. 1992. Reproductive biology of the holothurian *Holothuria forskali* (Echinodermata). J Mar Biol Ass 72:745–758. doi:10.1017/S0025315400060021

Vullien A, Amiel A, Baduel L, Diken D, Renaud C, Vervoort M, Röttinger E, Gazave E. 2024. The rich evolutionary history of the ROS metabolic arsenal shapes its mechanistic plasticity at the onset of metazoan regeneration. doi:10.1101/2024.07.25.605162

Wahltinez SJ, Byrne M, Stacy NI. 2023. Coelomic fluid of asteroid echinoderms: Current knowledge and future perspectives on its utility for disease and mortality investigations. Vet Pathol 60:547–559. doi:10.1177/03009858231176563

Wambreuse N, Caulier G, Eeckhaut I, Borrello L, Bureau F, Fievez L, Delroisse J. 2025. Morpho-functional characterisation of cœlomocytes in the aquacultivated sea cucumber *Holothuria scabra*: From cell diversity to transcriptomics immune response. Fish & Shellfish Immunology 158:110144. doi:10.1016/j.fsi.2025.110144

Wambreuse N, Coubris C, Mallefet J, Delroisse J, Caulier G, Duchatelet L. 2024. Insights into ophiuroid coelomocytes: diversity and potential function in coelenterazine transfer in the bioluminescent brittle star *Amphiura filiformis*. Cahiers de Biologie Marine 65:357–369. doi:10.21411/CBM.A.FBA7A5E7

Wang S, Wang H, Zhao L, Zhang Y, Li T, Liu S, Shi J, Lian S, Hu J, Bao Z, Hu X. 2022. Identification of genes associated with carotenoids accumulation in scallop (*Patinopecten yessoensis*). Aquaculture 550:737850. doi:10.1016/j.aquaculture.2021.737850

Weaver RJ, Gonzalez BK, Santos SR, Havird JC. 2020. Red Coloration in an Anchialine Shrimp: Carotenoids, Genetic Variation, and Candidate Genes. The Biological Bulletin 238:119–130. doi:10.1086/708625

Wessel GM, Goldberg L, Lennarz WJ, Klein WH. 1989. Gastrulation in the sea urchin is accompanied by the accumulation of an endoderm-specific mRNA. Developmental Biology 136:526–536. doi:10.1016/0012-1606(89)90278-9

Williams ST, Heyworth SM, Kano Y, Roberts NW, Carter HF, Cheney KL. 2025. The blue advantage: a novel blue carotenoprotein pigment in the tropical seastar *Linckia laevigata* is an antioxidant defence against extreme environmental stress. Mar Biol 172:31. doi:10.1007/s00227-025-04595-7

Wu X, Chen T, Huo D, Yu Z, Ruan Y, Cheng C, Jiang X, Ren C. 2020. Transcriptomics analysis of sea cucumber (*Holothuria leucospilota*) coelomocytes revealed the echinoderm cytokine response during immune challenge. BMC Genomics 21:306. doi:10.1186/s12864-020-6698-6

Wybouw N, Kurlovs AH, Greenhalgh R, Bryon A, Kosterlitz O, Manabe Y, Osakabe M, Vontas J, Clark RM, Van Leeuwen T. 2019. Convergent evolution of cytochrome P450s underlies independent origins of keto-carotenoid pigmentation in animals. Proc R Soc B 286:20191039. doi:10.1098/rspb.2019.1039

Xiao K, Zhang S, Li C. 2022. The complement system and complement-like factors in sea cucumber. Developmental & Comparative Immunology 136:104511. doi:10.1016/j.dci.2022.104511

Xing J, Leung M-F, Chia F-S. 1998. Quantitative Analysis of Phagocytosis by Amebocytes of a Sea Cucumber, *Holothuria leucospilota*. Invertebrate Biology 117:67. doi:10.2307/3226853

Xing K, Yang H, Chen M. 2008. Morphological and ultrastructural characterization of the coelomocytes in *Apostichopus japonicus*. Aquat Biol 2:85–92. doi:10.3354/ab00038

Xing L, Wang L, Liu S, Sun L, Wessel GM, Yang H. 2023. Single-Cell Transcriptome and Pigment Biochemistry Analysis Reveals the Potential for the High Nutritional and Medicinal Value of Purple Sea Cucumbers. IJMS 24:12213. doi:10.3390/ijms241512213

Xu W, Yang Z, Lu N. 2015. A new role for the PI3K/Akt signalling pathway in the epithelial-mesenchymal transition. Cell Adhesion & Migration 9:317–324. doi:10.1080/19336918.2015.1016686

Xu D, Sun L, Liu S, Zhang L, Yang H. 2016. Understanding the Heat Shock Response in the Sea Cucumber *Apostichopus japonicus*, Using iTRAQ-Based Proteomics. IJMS 17:150. doi:10.3390/ijms17020150

Yu K, Zhao X, Xiang Y, Li C. 2023. Phenotypic and functional characterization of two coelomocyte subsets in *Apostichopus japonicus*. Fish & Shellfish Immunology 132:108453. doi:10.1016/j.fsi.2022.108453

Zheng Y, Min J, Kim D, Park J, Choi S-W, Lee E, Na K, Bae S. 2016. In Vitro Inhibition of Human UDP-Glucuronosyl-Transferase (UGT) Isoforms by Astaxanthin, β-Cryptoxanthin, Canthaxanthin, Lutein, and Zeaxanthin: Prediction of in Vivo Dietary Supplement-Drug Interactions. Molecules 21:1052. doi:10.3390/molecules21081052

