## Supplementary material for "Carotenoid-based immune response in sea cucumbers relies on newly identified coelomocytes – the carotenocytes": Fig. S1

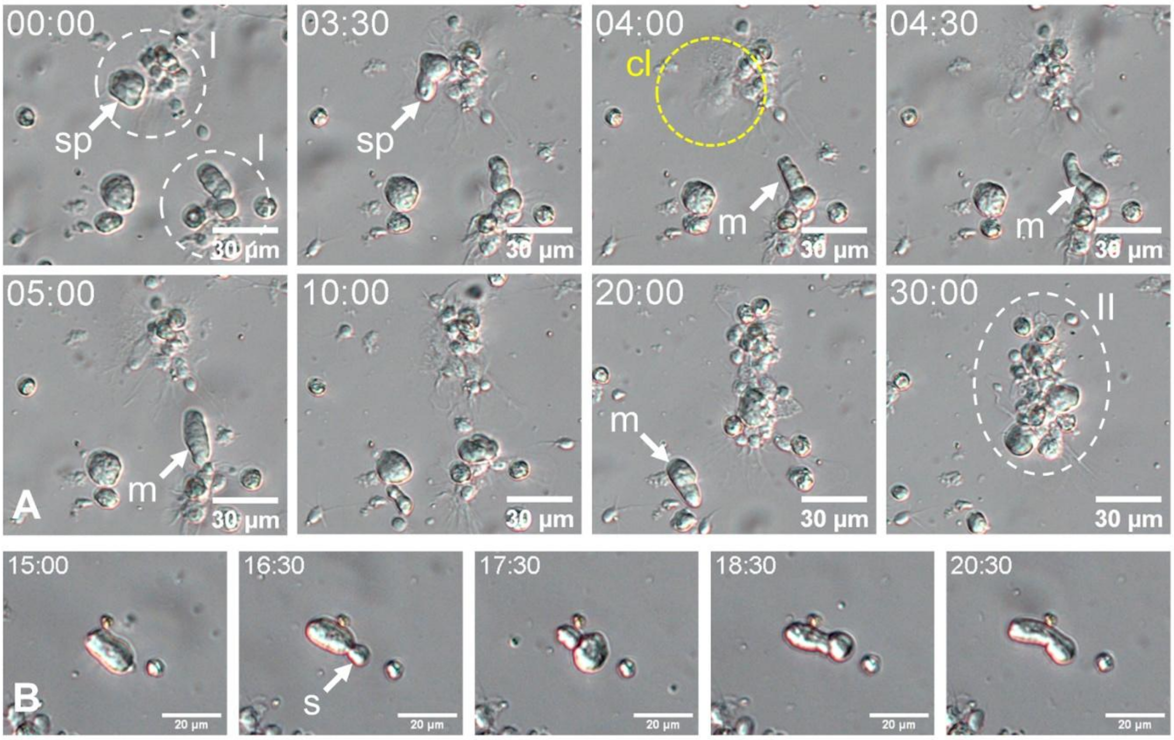


**Fig. S1.** Spherule cell-specific behaviours and mobility. A. Time-lapse imaging (same as **Video S1**) showing a spherule cell achieving lysis in contact with an aggregate. Some spherule cells also display high mobility. B. Particular movement of a small spherulocyte displaying a bleb-driven-like mobility starting with a small spheroid at the apex of the cell, followed by an undulation throughout the cell membrane (time frames: min:sec). Legend: cl - cell lysis; I - stage I aggregate; II - stage II aggregate; m - mobility; s - spheroid; sp - spherule cell.
