## Supplementary material for "Carotenoid-based immune response in sea cucumbers relies on newly identified coelomocytes – the carotenocytes": Fig. S2

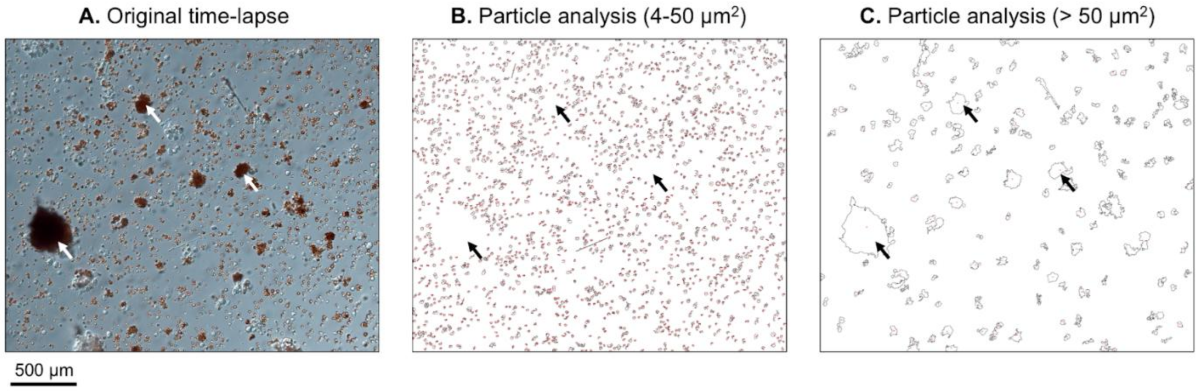


**Fig. S2.** Automated cell analysis using the Particle Analysis tool in ImageJ software. A. Original time-lapse converted to video in ImageJ. B. After applying a threshold filter, particle analysis was performed on an area between 4 and 50 $\mu m^2$ to target hemocyte-type cells. Each number corresponds to a cell. C. Particle analysis on an area greater than 50 $\mu m^2$, mainly targeting cell aggregates. Arrows indicate large aggregates in which no small cells (i.e., particles < 50 $\mu m^2$) are visible.
