## Supplementary material for "Carotenoid-based immune response in sea cucumbers relies on newly identified coelomocytes – the carotenocytes": Fig. S3

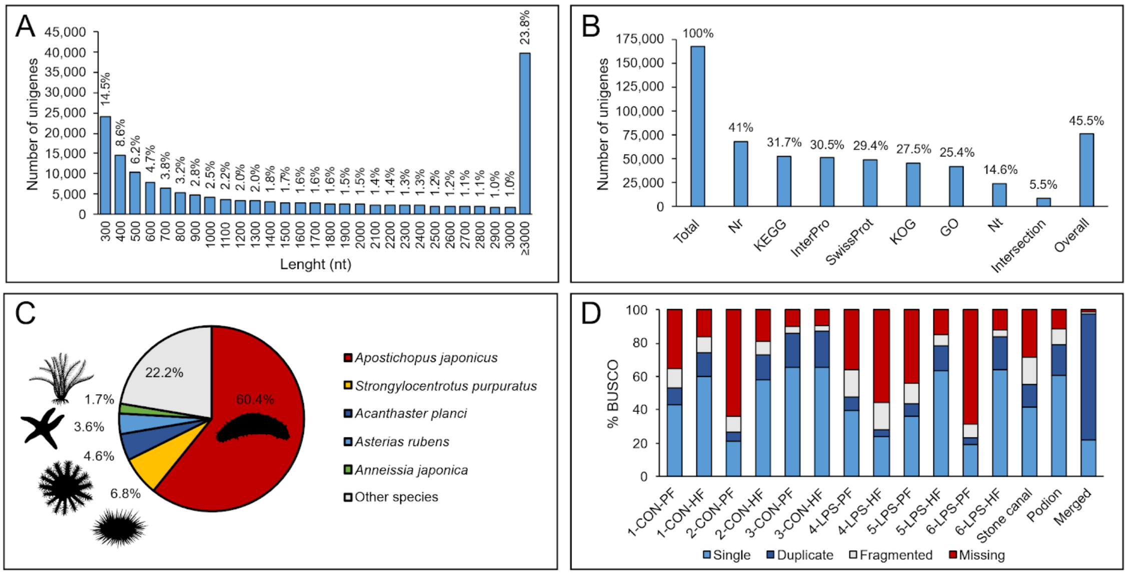


**Fig. S3.** Transcriptome assembly and general annotation in *Holothuria forskali*. A. Length distribution of unigenes (corresponding percentages are embedded in the graph). B. General annotations of unigenes against seven functional databases (match percentages are embedded in the graph). C. Species distribution for the Nr annotation (match percentages are embedded in the graph; species icons were modified from PhyloPic (www.phylopic.org). D. BUSCO assessment graph (CON – control individual; HF – hydrovascular fluid; LPS – LPS-injected individual; PF – perivisceral fluid).
