## Supplementary material for "Carotenoid-based immune response in sea cucumbers relies on newly identified coelomocytes – the carotenocytes": Fig. S4

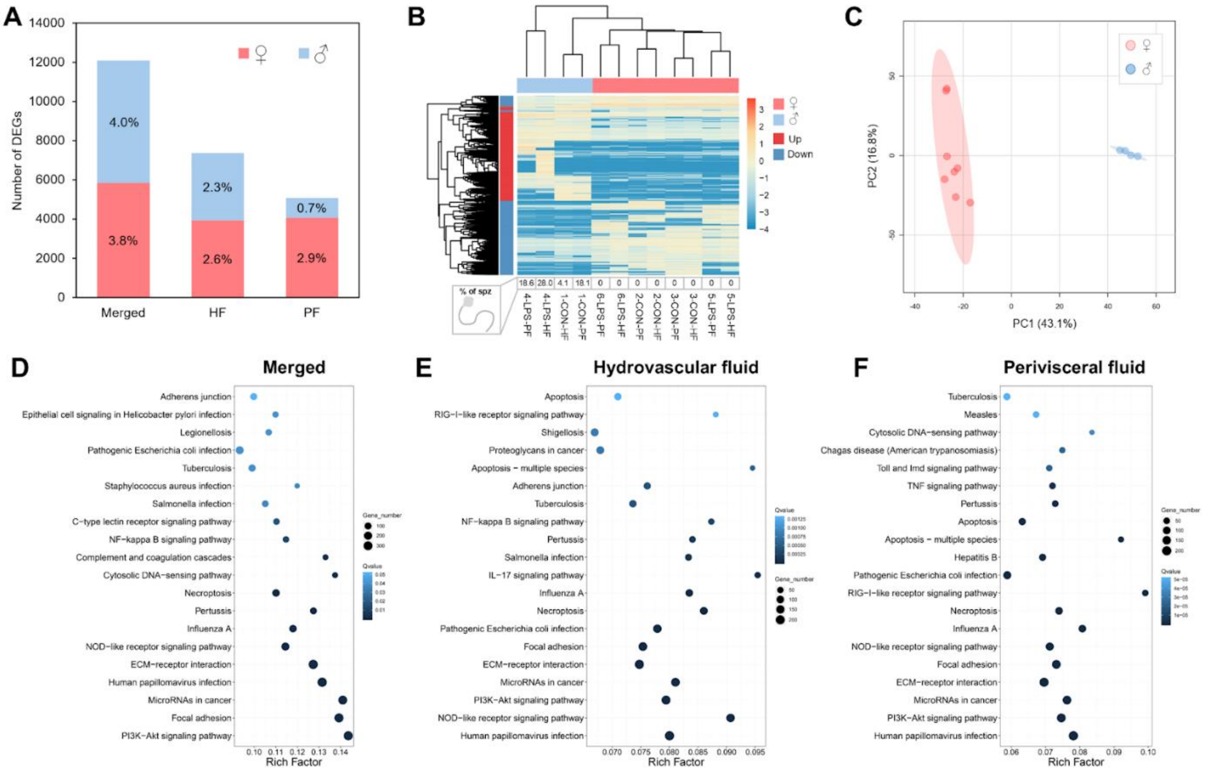


**Fig. S4.** Differential expression analysis of coelomocytes between males and females in *Holothuria forskali*. **A.** Number of differentially expressed genes between males () and females (). The percentage of DEGs is included on the graph. The analysis was done regardless of the body fluid (merged), in hyrdrovascular fluid (HF) and perivisceral fluid (PF). **B.** Heatmap based on the DEGs of the merged analysis: female samples form an accurate cluster distinct from males. The percentage of spermatozoa (% of spz) in each sample is displayed at the bottom of the heatmap, they were present only in male samples. C. a PCA analysis based on the DEGs of the merged analysis (based on the 25% most informative genes filtered by interquartile), female and male samples form distinct clusters with a higher variability in the expression within females. **D-F.** KEGG enrichment analyses showing the twenty most enriched pathways between the two sexes in the merged analysis, the HF and the PF, respectively. Colour scale represents the degree of enrichment, size of the dot, the number of genes for each pathway and rich factor (X axis), the number of DEGs enriched in the pathway out of the number of genes expressed in the pathway in the transcriptome.
