## Supplementary material for "Carotenoid-based immune response in sea cucumbers relies on newly identified coelomocytes – the carotenocytes": Fig. S5

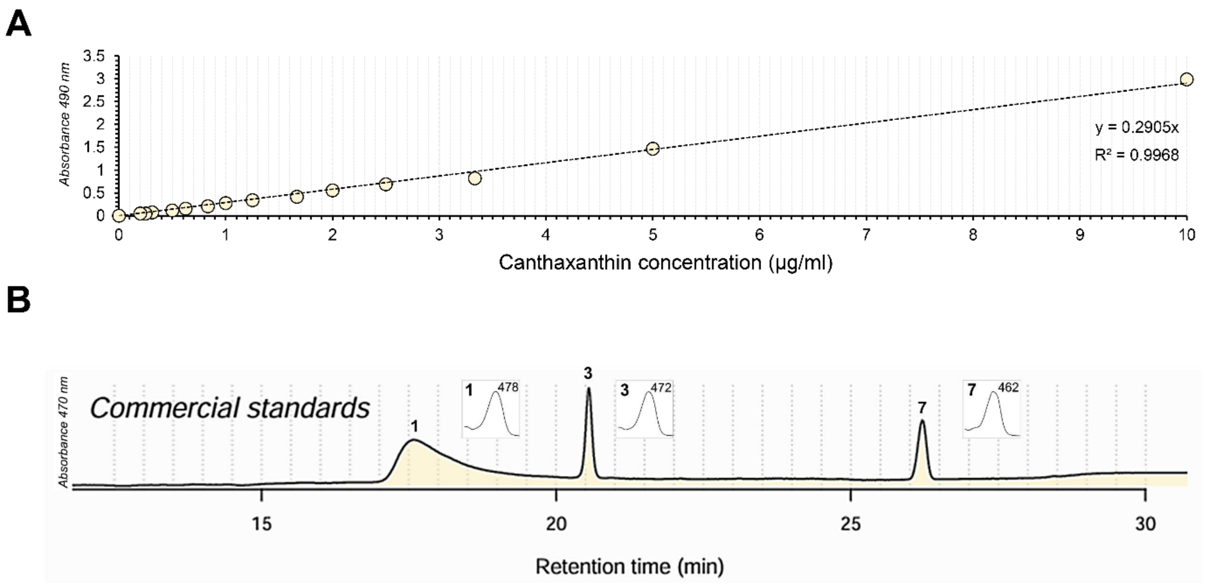


**Fig. S5.** Identification of pigment based on commercial standards. A. Calibration curve of pure canthaxanthin pigment (canthaxanthin_trans; Merck n°11775) of known concentration. The absorbance of the canthaxanthin standard was measured by spectrophotometry at increasing dilutions in chloroform (λ = 490 nm). The equation of the linear regression and the determination coefficient ($R^2$) indicated on the graph, this equation was used for estimating the carotenoid mass in coelomocyte extracts. B. Calibration curve in high-performance liquid chromatography (HPLC). The peaks 1, 3 and 6 on the HPLC spectra correspond to the commercial standards for astaxanthin, canthaxanthin and all-trans-echinenone, respectively (DHI Laboratory Products). The profiles of each peak (300–600 nm) are presented, confirming their identification.
