## Supplementary material for "Carotenoid-based immune response in sea cucumbers relies on newly identified coelomocytes – the carotenocytes": Fig. S6

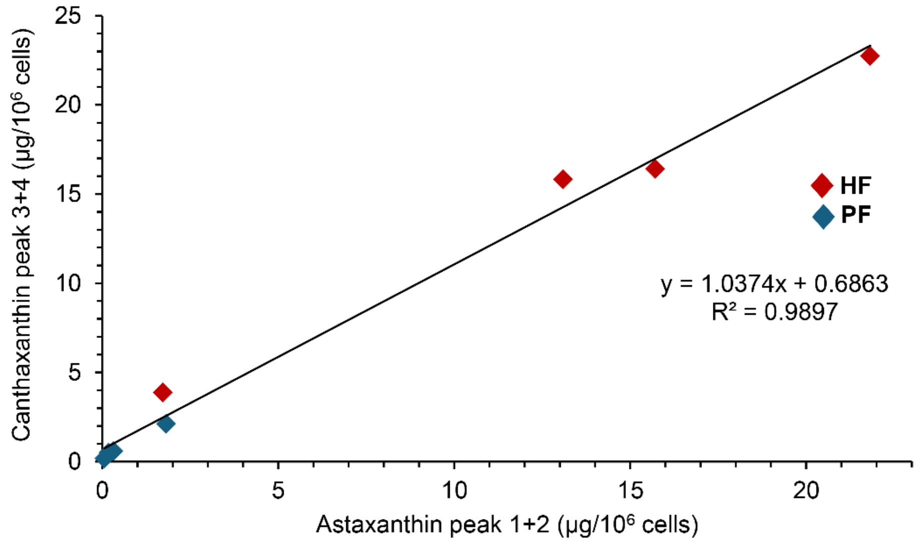


**Fig. S6.** Linear relationship between astaxanthin and canthaxanthin concentrations as determined by high-performance liquid chromatography (HPLC), in perivisceral fluid and in hydrovascular fluid of *Holothuria forskali*. Peaks refer to the spectra shown in Figure 6F. Peak 1+2: addition of the concentrations of the areas under the peaks (AUP) corresponding to astaxanthin. Peak 3+4: addition of the concentration of the areas under the peak (AUP) corresponding to canthaxanthin. The concentrations of these two types of carotenoids demonstrate a high coefficient of determination ($R^2$). The equation of the linear regression and the determination coefficient ($R^2$) are indicated on the graph. Legend: HF – hydrovascular fluid; PF – perivisceral fluid**.**
