## Supplementary material for "Carotenoid-based immune response in sea cucumbers relies on newly identified coelomocytes – the carotenocytes": Fig. S7

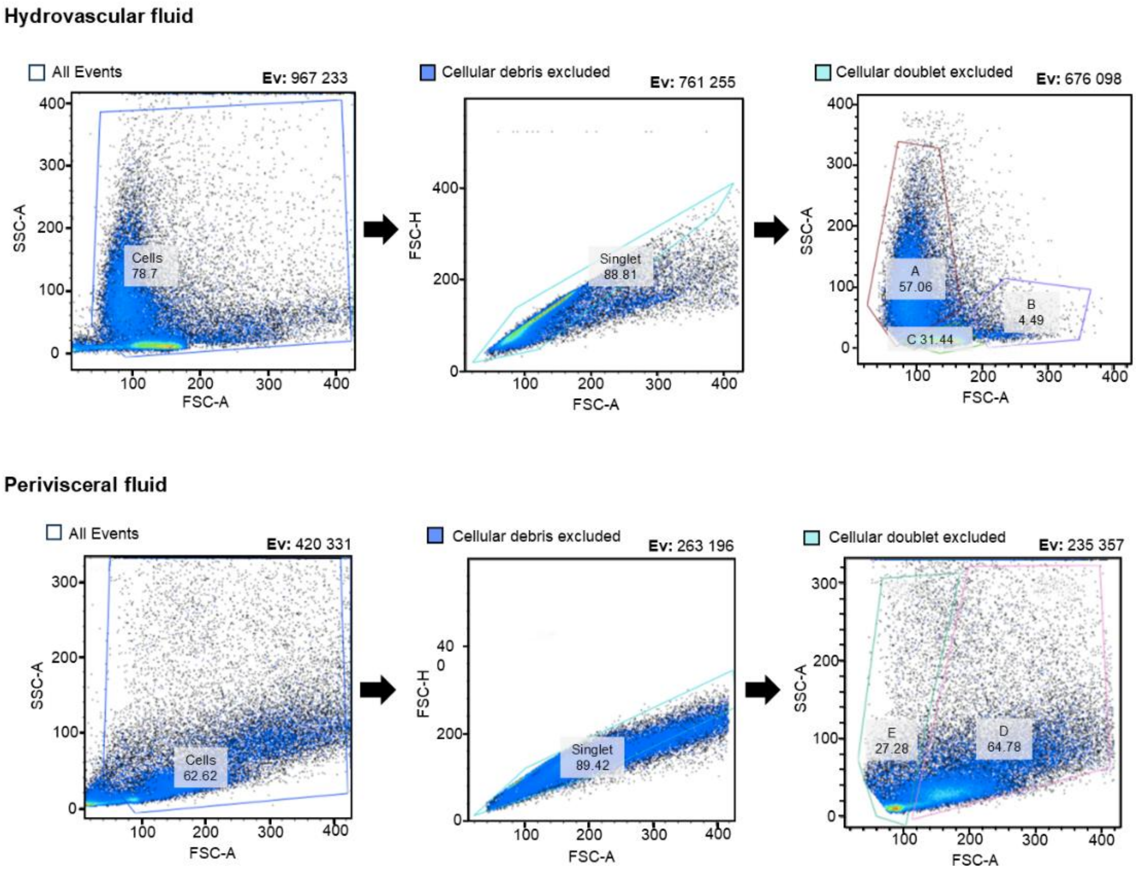


**Fig. S7.** Spectral flow cytometry gating strategy of coelomocytes in the hydrovascular (A) and perivisceral (B) fluids *of Holothuria forskali*. The percentages indicated correspond to the percentage of cells (cell events) selected. The size and granularity parameters (FSC-A and SSC-A) were utilised to eliminate cellular debris, which typically exhibits minimal levels of these parameters. Subsequently, the parameters of size (FSC-A) and height (FSC-H) were utilised to eliminate cell doublets, which possess an increased surface area. Finally, the selection of different populations was based on size and granularity parameters (FSC-A and SSC-A). Legend: Ev – number of cell events.
