## Supplementary material for "Carotenoid-based immune response in sea cucumbers relies on newly identified coelomocytes – the carotenocytes": Fig. S8

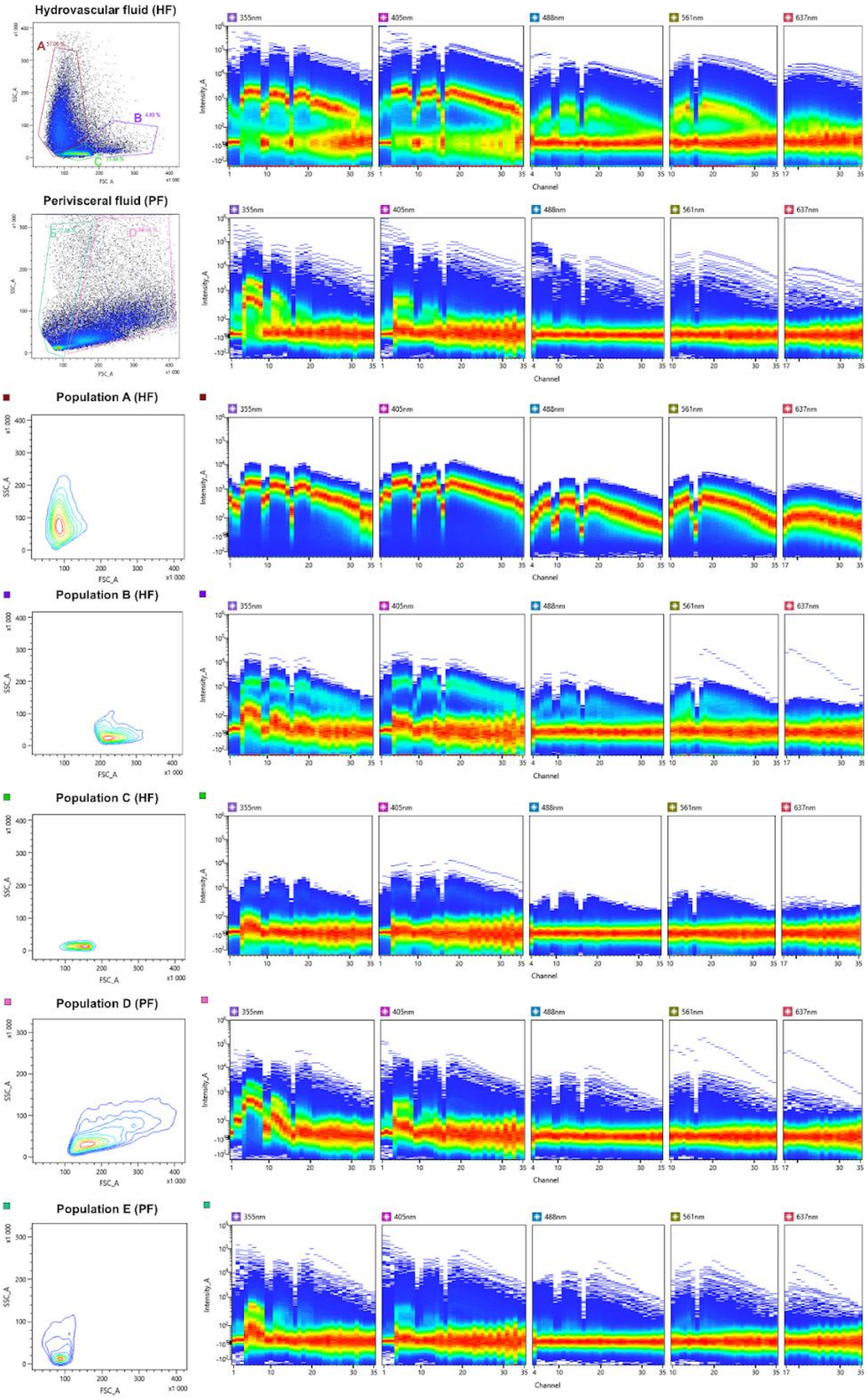


**Fig. S8.** Autofluorescence spectra of coelomocytes revealed by spectral flow cytometry. The first two spectra correspond to the two body fluids, respectively, perivisceral fluid (PF) and hydrovascular fluid (HF). The order corresponds to the different defined populations in Fig. 7. It is noticed that HF show a higher proportion of autofluorescent cells, and that among these population A, corresponding to Hemocyte-like cells (HELs), is the most autofluorescent population in all the lasers used.
