## Supplementary material for "Carotenoid-based immune response in sea cucumbers relies on newly identified coelomocytes – the carotenocytes": Fig. S9

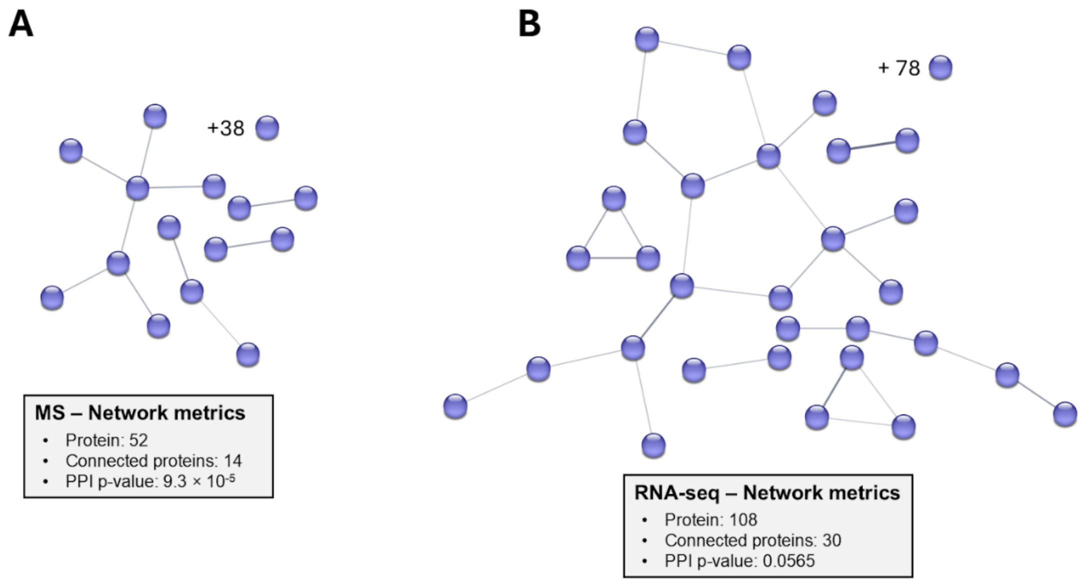


**Fig. S9.** String protein networks based on genes overexpressed in the perivisceral fluid (PF), in comparison to the hydrovascular fluid (HF: networks in Fig. 11). **A.** and **B.** are proteins identified based on mass-spectrometry (MS) analysis (fold change (HFvsPF) > 5) and RNA-sequencing (RNA-seq) analysis (fold change (HFvPF) > 2), respectively. It can be noticed that protein networks are smaller than those of HF, with lower protein-protein interaction p-values.
