## Supplementary material for "Carotenoid-based immune response in sea cucumbers relies on newly identified coelomocytes – the carotenocytes": Fig. S10

**A**

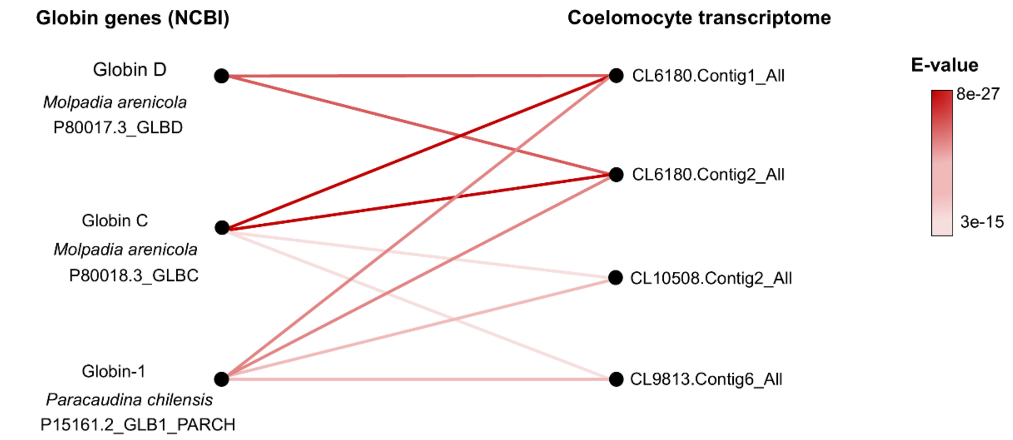


**B**

| GeneID | Annotation NR | E-value | $Log_2(FC)$ | FDR |
|---|---|---|---|---|
| CL6180.Contig1_All | Globin C, coelomic [*Holothuria leucospilota*] | $7 \times 10^{-112}$ | 2.14 | 0.12 |
| CL6180.Contig2_All | Globin C, coelomic [*Holothuria leucospilota*] | $4 \times 10^{-110}$ | 2.02 | 0.34 |
| CL10508.Contig2_All | Globin [*Holothuria leucospilota*] | $2 \times 10^{-120}$ | -0.33 | 1.00 |
| CL9813.Contig6_All | Globin [*Holothuria leucospilota*] | $3 \times 10^{-122}$ | 0.10 | 1.00 |

**C**

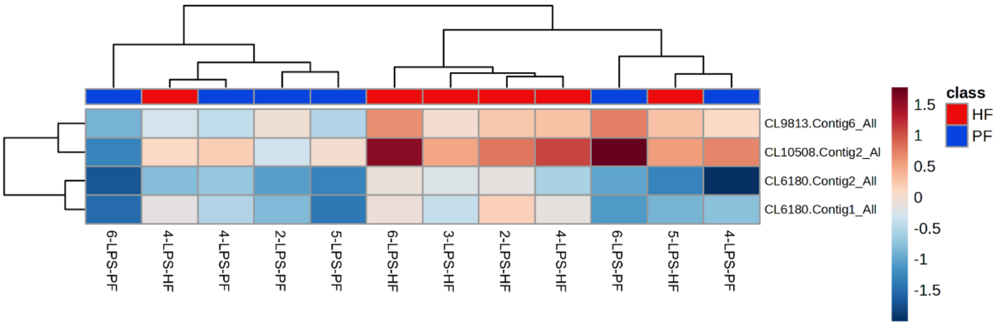


**Fig. S10.** Globin expression in coelomocytes of perivisceral and hydrovascular fluids. A. Relation between the unigenes expressed in *H. forskali* and the globin sequences expressed in the coelomocytes of other holothurian species (NCBI; according to Christensen et al., 2015). The degree of relation is assessed using the BLAST E-value, which indicates the statistical significance of the similarity between sequences. Four unigenes expressed in the *H. forskali* transcriptome showed a strong degree of correspondence with the holothurian intracellular globins (E-value $\leq 10^{-5}$). B. Table summarising the analysis of the expression of unigenes annotated as globins in holothurians (NCBI NR), indicating for each unigene of *H. forskali* transcriptome, the NR annotation and associated E-value, the Log2 (FC) and the FDR associated with unigene expression between the hydrovascular fluid (HF) and perivisceral fluid (PF). C. Heatmap representing the FPKM expression level of the unigenes expressed in the fluids. Only unigenes having a |log2(FC) value| ≥ 1 and an FDR ≤ 5% were considered as significantly differentially expressed. The unigenes corresponding to globins were not found to be differentially expressed between the PF and HF samples. The Padj values for each unigene did not demonstrate significant differences between the two fluids ($P \geq 0.05$), and the heatmaps illustrate a clustering that does not fit the PF and HF distinction. Legend: CON – control condition; GeneID – gene identification code; IND = individual; $Log_2$ (FC) – Fold Change value under $log_2$ transformation; LPS – immunostimulated condition. FDR – False Discovery Rate.
