## Supplementary material for "Carotenoid-based immune response in sea cucumbers relies on newly identified coelomocytes – the carotenocytes": Fig. S11

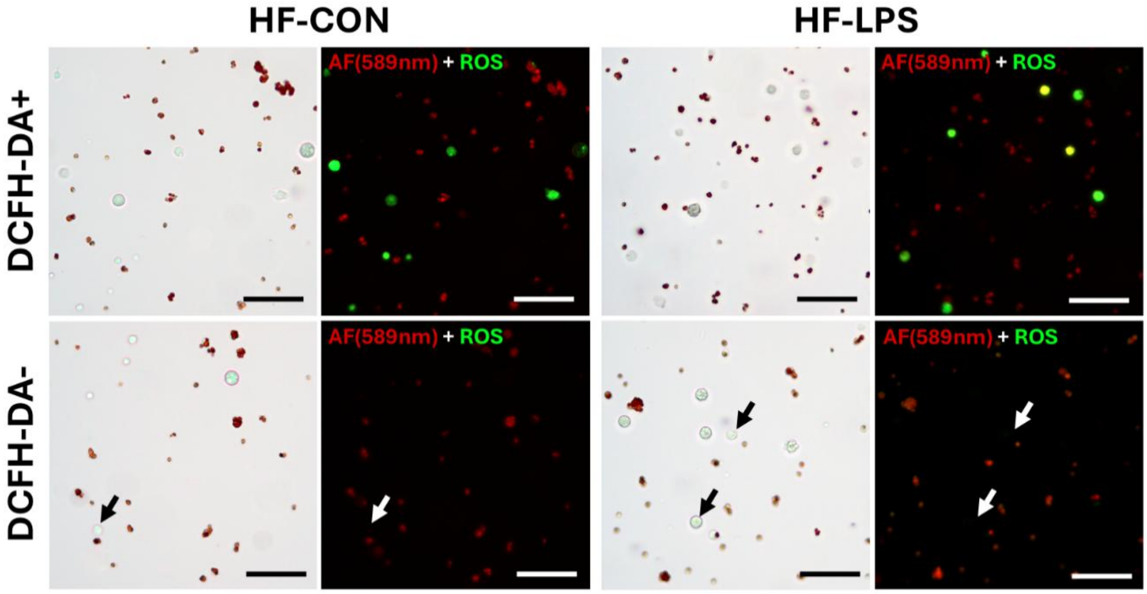


**Fig. S11.** Validation of DCFH-DA (2′−7′-dichlorodihydrofluorescein diacetate) labelling for reactive oxygen species (ROS) detection. Comparison of the same sample with and without the immunological stress (50µg/ml of lipopolysaccharides), and with the labelling of DCFH-DA (above) and without DCFH-DA (negative control – below). Positive cells (in green) are only observed in the presence of DCFH-DA. Autofluorescence at 589 nm was used to highlight the presence of hemocyte-like cells (HELS – in red). Note that these are not positive to ROS (the arrow indicates where cells are when they are not visible in fluorescent microscopy). The scale bar represents 50 µm (CON – control; HF – hydrovascular fluid; LPS – lipopolysaccharide-exposure).
