## Supplementary material for "Carotenoid-based immune response in sea cucumbers relies on newly identified coelomocytes – the carotenocytes": Fig. S12

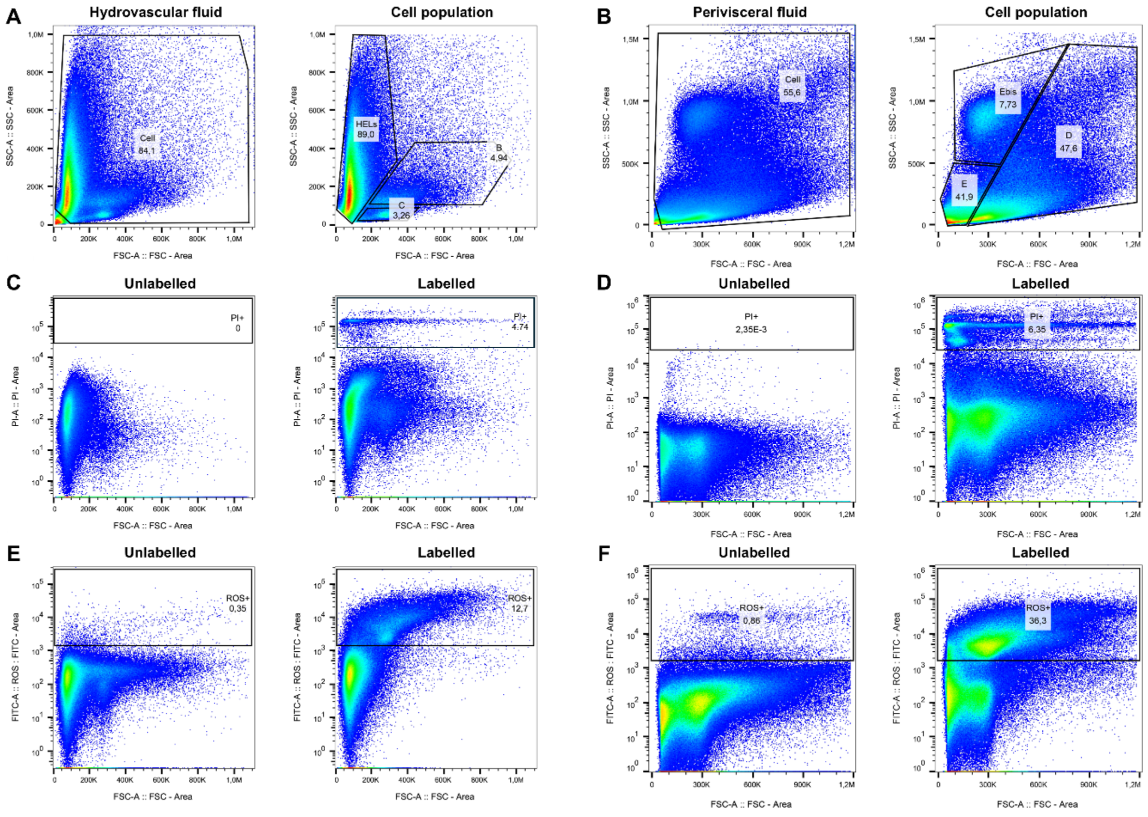


**Fig. S12.** Flow cytometry gating strategy of ROS production analysis. A. and B. Flow cytometry profile of hydrovascular (HF) and perivisceral fluids (PF), respectively. C. and D. Cell mortality analysis using propidium iodide labelling in the HF and the PF, respectively. Negative control (i.e., unlabelled sample) is shown for each fluid. E. and F. ROS production analysis using propidium iodide labelling in the HF and the PF, respectively. Negative control (i.e., unlabelled sample) is shown for each fluid.
