## Supplementary material for "Carotenoid-based immune response in sea cucumbers relies on newly identified coelomocytes – the carotenocytes": Fig. S13

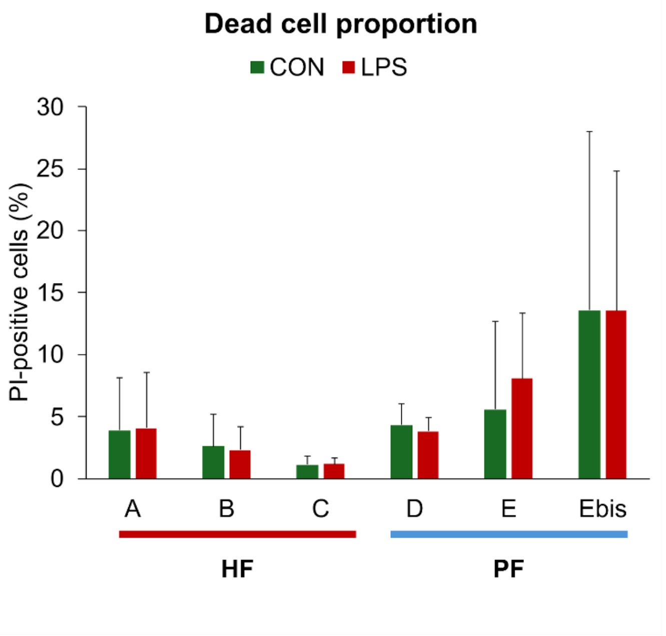


**Fig. S13.** Estimation of cell mortality using propidium iodide labelling and flow cytometry. Cell mortality proportion was low, with no clear difference between populations or between coelomocytes exposed or not to lipopolysaccharides. Different populations correspond to those defined in Fig. 13. Legend: CON – control; HF – hydrovascular fluid; LPS – lipopolysaccharide-exposure; PF – perivisceral fluid.
