## Supplementary material for "Carotenoid-based immune response in sea cucumbers relies on newly identified coelomocytes – the carotenocytes": Table S1

**Tables S1.** Statistical analysis results between the hydrovascular fluid (HF) and perivisceral fluid (PF), in specimens of normal condition (no injection). Results are formulated as mean ± SD (n = 6), and p-values (P) show significant differences between the two fluids (Wilcoxon signed rank test; significant differences are in bold; n.a. means not applicable).

| | Concentration (cells ml$^{-1}$) | | | | Proportion (%) | | | |
|---|---|---|---|---|---|---|---|---|
| **Coelomocyte types** | **HF** | **PF** | **P** | **W** | **HF** | **PF** | **P** | **W** |
| **Phagocyte** | 9.28 ± 6.18 $10^5$ | 1.04 ± 0.45 $10^6$ | 1 | -1 | 10.52 ± 11.33 | 30.98 ± 10.07 | **0.031** | **-21** |
| **Small spherulocyte** | 3.27 ± 1.44 $10^5$ | 1.19 ± 0.41 $10^6$ | **0.035** | **-21** | 2.91 ± 1.45 | 36.04 ± 5.90 | **0.031** | **-21** |
| **Large spherulocyte** | 1.05 ± 1.28 $10^5$ | 1.41 ± 1.16 $10^5$ | 0.27 | -7 | 1.07 ± 1.12 | 5.05 ± 3.14 | **0.031** | **-21** |
| **Small round cell** | 5.78 ± 4.16 $10^5$ | 7.37 ± 1.61 $10^5$ | 0.31 | -11 | 4.49 ± 2.69 | 23.44 ± 7.36 | **0.031** | **-21** |
| **Hemocyte-like cell** | 1.20 ± 0.96 $10^7$ | 2.33 ± 2.34 $10^4$ | **0.031** | **21** | 80.62 ± 5.14 | 0.75 ± 0.71 | 0.031 | 21 |
| **Fusiform cell** | 5.67 ± 5.99 $10^4$ | 1 ± 1.37 $10^5$ | 0.78 | -3 | 0.38 ± 0.35 | 3.09 ± 3.75 | **0.19** | **-11** |
| **Crystal cell** | 3.33 ± 8.16 $10^3$ | 2.0 ± 0.0 $10^4$ | **0.037** | **-15** | 0.01 ± 0.03 | 0.64 ± 0.15 | **0.031** | **-21** |
| **Total (all types)** | 1.40 ± 0.99 $10^7$ | 3.28 ± 0.83 $10^6$ | **0.031** | **21** | n.a. | n.a. | n.a. | n.a. |
