## Supplementary material for "Carotenoid-based immune response in sea cucumbers relies on newly identified coelomocytes – the carotenocytes": Table S2

**Table S2.** Statistical analysis results of concentration and proportion of coelomocyte populations between control and LPS-injected individuals (Mann-Whitney U test; significant differences are in bold).

| | Perivisceral fluid (PF) | | | | Hydrovascular fluid (HF) | | | |
|---|---|---|---|---|---|---|---|---|
| **Cell types** | **Control** | **LPS** | **P** | **U** | **Control** | **LPS** | **P** | **U** |
| | **Concentration** | | | | | | | |
| **Phagocyte** | $1.6 \pm 4.4\ 10^6$ | $3.7 \pm 3.8\ 10^6$ | 0.7308 | 18 | $1.7 \pm 1.3\ 10^6$ | $1.8 \pm 1.2\ 10^6$ | 0.7206 | 18 |
| **Small spherulocyte** | $1.1 \pm 0.4\ 10^6$ | $2.6 \pm 2.2\ 10^6$ | 0.4452 | 15 | $1 \pm 1.2\ 10^6$ | $8.9 \pm 6.2\ 10^5$ | 1 | 20.5 |
| **Large spherulocyte** | $2.5 \pm 1.3\ 10^5$ | $1 \pm 0.1\ 10^6$ | 0.2234 | 12 | $2.9 \pm 4\ 10^5$ | $8.3 \pm 9.2\ 10^5$ | 0.1 | 9 |
| **Small round cell** | $9.5 \pm 11.6\ 10^5$ | $1.1 \pm 1.1\ 10^6$ | 0.7206 | 18 | $1.8 \pm 3.6\ 10^6$ | $1 \pm 1.1\ 10^6$ | 0.3518 | 14 |
| **Hemocyte** | $1.7 \pm 4\ 10^4$ | $2.3 \pm 4.3\ 10^5$ | 0.4314 | 16 | $3.2 \pm 3.2\ 10^6$ | $3.1 \pm 2.1\ 10^7$ | **0.0012** | **0** |
| **Fusiform cell** | $2.4 \pm 2.8\ 10^5$ | $5.2 \pm 7.9\ 10^5$ | 0.6678 | 17.5 | $2.2 \pm 2.2\ 10^5$ | $1.8 \pm 2.3\ 10^5$ | 0.828 | 19 |
| **Crystal cell** | $3.3 \pm 6.2\ 10^4$ | $4.9 \pm 6\ 10^4$ | 0.2328 | 12.5 | $1.7 \pm 2.3\ 10^4$ | $2 \pm 2.6\ 10^4$ | 1 | 20.5 |
| **Total** | $4.2 \pm 1.8\ 10^6$ | $8.9 \pm 7.6\ 10^6$ | 0.7308 | 18 | $8.2 \pm 8.1\ 10^6$ | $3.5 \pm 2.3\ 10^7$ | **0.0082** | **3** |
| | **Proportion** | | | | | | | |
| **Phagocyte** | 42.1 ± 14 % | 41.3 ± 10.7 % | 0.8301 | 19 | 25.6 ± 11.6 % | 5.7 ± 3.4 % | **0.0053** | 1 |
| **Small spherulocyte** | 28 ± 9.7 % | 27.9 ± 9.3 % | 0.9452 | 20 | 13.3 ± 6.1 % | 2.8 ± 1.8 % | **0.0023** | 1 |
| **Large spherulocyte** | 11 ± 8.8 % | 5.6 ± 1.4 % | 0.2949 | 13 | 3.6 ± 2.1 % | 2.3 ± 2.7 % | 0.3525 | 14 |
| **Small round cell** | 12.9 ± 8.5 % | 18.2 ± 14.5 % | 0.6282 | 17 | 12.8 ± 13 % | 2.9 ± 2.3 % | **0.014** | 4 |
| **Hemocyte** | 0.2 ± 0.3 % | 0.6 ± 1.6 % | 0.5414 | 17 | 41 ± 25.1 % | 85.3 ± 6.6 % | **0.0047** | 2 |
| **Fusiform cell** | 4.9 ± 6.5 % | 5.3 ± 5.2 % | 0.7748 | 18.5 | 3.4 ± 3.9 % | 0.5 ± 0.5 % | **0.0123** | 3 |
| **Crystal cell** | 0.5 ± 0.3 % | 0.6 ± 0.9 % | 0.6621 | 17.5 | 0.1 ± 0.2 % | 0 ± 0.1 % | 0.5855 | 17 |
