## Supplementary material for "Carotenoid-based immune response in sea cucumbers relies on newly identified coelomocytes – the carotenocytes": Table S3

**Table S3**. Metrics of the *de novo* RNA-sequencing in *Holothuria forskali*. For each individual transcriptome and the merged transcriptome, the table provides: the sex of the individuals, metrics on raw data (reads metrics) and assembled data (assembly metrics). Read metrics include the total number of raw reads, the total number of clean reads, the total number of clean bases, the percentage of bases whose quality is greater than 20 in clean reads (Q20), and the percentage of clean reads. Assembly metrics include: the total number of unigenes; the total length of the transcriptome; the mean length of unigenes; the N50; and the ratio of GC bases. Legend: ♂ – male; ♀ – female; n.a. – not applicable; n.d. – non-determined; CON – control individual; HF – hydrovascular fluid sample; LPS – lipopolysaccharide individual; PF – perivisceral fluid sample.

| Samples | | Read metrics | | | | | Assembly metrics | | | | |
|---|---|---|---|---|---|---|---|---|---|---|---|
| **Name** | **Sex** | **Raw reads (M)** | **Clean reads (M)** | **Clean bases (Gb)** | **Q20 (%)** | **Clean reads (%)** | **Total number** | **Total length (M)** | **Mean length** | **N50** | **GC (%)** |
| **1-CON-PF** | ♂ | 233.89 | 221.38 | 22.14 | 99.0 | 94.7 | 42,190 | 48.85 | 1,157 | 2,054 | 38.4 |
| **1-CON-HF** | ♂ | 236.37 | 221.93 | 22.19 | 99.0 | 93.9 | 59,097 | 77.01 | 1,303 | 2,490 | 39.1 |
| **2-CON-PF** | ♀ | 241.35 | 221.49 | 22.15 | 99.1 | 91.8 | 33,934 | 33.40 | 984 | 1,500 | 38.0 |
| **2-CON-HF** | ♀ | 243.84 | 221.00 | 22.10 | 99.0 | 90.6 | 57,175 | 70.75 | 1,237 | 2,301 | 38.9 |
| **3-CON-PF** | ♀ | 236.37 | 221.78 | 22.18 | 99.1 | 93.8 | 56,407 | 81.03 | 1,436 | 2,740 | 39.7 |
| **3-CON-HF** | ♀ | 233.89 | 220.49 | 22.05 | 99.0 | 94.3 | 55,941 | 82.70 | 1,478 | 2,817 | 39.7 |
| **4-LPS-PF** | ♂ | 233.89 | 221.74 | 22.17 | 99.0 | 94.8 | 42,789 | 47.42 | 1,108 | 1,965 | 38.5 |
| **4-LPS-HF** | ♂ | 241.35 | 220.62 | 22.06 | 98.9 | 91.4 | 46,473 | 41.56 | 894 | 1,447 | 38.1 |
| **5-LPS-PF** | ♀ | 241.35 | 221.50 | 22.15 | 98.9 | 91.8 | 37,473 | 39.18 | 1,045 | 1,773 | 38.2 |
| **5-LPS-HF** | ♀ | 236.37 | 220.75 | 22.08 | 98.8 | 93.4 | 57,538 | 75.27 | 1,308 | 2,490 | 39.2 |
| **6-LPS-PF** | ♀ | 241.35 | 221.87 | 22.19 | 98.8 | 91.9 | 30,861 | 27.73 | 898 | 1,269 | 38.1 |
| **6-LPS-HF** | ♀ | 236.37 | 222.51 | 22.25 | 98.8 | 94.1 | 55,386 | 77.09 | 1,391 | 2,629 | 39.4 |
| **Stone canal** | n.d. | 241.35 | 220.28 | 22.03 | 99.0 | 91.3 | 58,800 | 71.97 | 1,223 | 2,267 | 38.8 |
| **Podia** | n.d. | 233.89 | 220.25 | 22.02 | 99.1 | 94.2 | 73,673 | 95.07 | 1,290 | 2,425 | 39.4 |
| **Total** | n.a. | 3331.63 | 3097.59 | 309.76 | 99.0 | 93.0 | 167,199 | 337.36 | 2,017 | 3,807 | 38.9 |
